# Cortical Thickness and Curvature in Autism and ADHD: A Mega-Analysis

**DOI:** 10.64898/2026.01.05.697782

**Authors:** Maryam Mahmoudi, Lucille A. Moore, Jacob Lundquist, Andrew Stier, Michael Anderson, Begim Fayzullobekova, Robert Hermosillo, Audrey Houghton, Thomas J. Madison, rae McCollum, Kimberly B. Weldon, Steve Nelson, Amy Esler, Oscar Miranda-Dominguez, Damien A. Fair, Brenden Tervo-Clemmens, Eric Feczko

## Abstract

**Background:** Existing evidence suggests cortical morphometric alterations occur in people with autism and ADHD. However, these findings remain tentative due to small sample sizes, heterogeneous imaging pipelines, varied statistical approaches, and limited harmonization across acquisition sites. Few studies have applied standardized processing to large, clinically enriched datasets or addressed site-related batch effects.

**Methods:** We leveraged six large-scale brain imaging datasets (n = 9,647; male=5,835; female=3,812; ages 5–64 years), including 1,533 individuals with ADHD, 1,080 with autism spectrum disorder, and 7,034 matched controls. All imaging data were processed using the validated ABCD-HCP pipeline, with cortical parcellation into 360 regions based on the Human Connectome Project (HCP) atlas, and ComBat harmonization was applied to account for variability across 67 acquisition sites. Group-level differences in cortical thickness and sulcal curvature were examined with ANCOVAs, controlling for covariates and using Bonferroni correction for multiple comparisons.

**Results:** Our analyses revealed distinct neuroanatomical signatures for both autism and ADHD. Individuals with autism exhibited regionally thinner cortex and curvature alterations particularly in the Cingulo-Opercular network. In contrast, individuals with ADHD displayed regionally thicker cortex, particularly in the default mode and somatomotor networks, alongside curvature differences. Control participants showed intermediate patterns, suggesting that autism and ADHD may represent diverging extremes of cortical maturation.

**Conclusions:** Cortical thickness and curvature emerge as potential biomarkers that can advance understanding of neurodevelopmental conditions and disentangle heterogeneity across diagnostic groups. These findings highlight the value of harmonized, large-scale, standardized analyses for resolving inconsistencies in the literature.

## Neurodevelopmental disabilities and overlapping symptoms, but distinct profiles

Neurodevelopmental disabilities (NDDs) diagnoses are highly prevalent^1,2^. Autism and ADHD are among the most common, affecting approximately 1 in 31 children and 11.4% of youth, respectively^1,3^. While autism and ADHD frequently co-occur and share symptoms, each presents distinct cognitive, emotional and behavioral profiles^1,4^. A systematic review highlights both overlapping and distinct patterns across domains such as executive function, attention, sensory processing, and face processing^5^. For example, autistic individuals show greater impairments in visual and sustained attention, while individuals with ADHD often show greater difficulties in auditory attention^6^.

Structural imaging provides a complementary, developmental perspective on brain maturation by exploring cortical morphometry, revealing both shared and distinct alterations across NDDs. Transmodal association cortices, such as the default mode and frontoparietal networks, develop later than primary sensory regions and remain plastic into adolescence^7^. Autism is associated with early cortical overgrowth, accelerated thinning during adolescence, and relative stabilization in adulthood^8^, while ADHD is characterized by delayed cortical maturation potentially due to slower synaptic pruning or microstructural maturation^9–11^. Both conditions may therefore disrupt similar late-developing networks but follow opposite structural trajectories, accelerated in autism and delayed in ADHD.

## Structural neuroimaging evidence for different neural substrates in autism and ADHD

Neuroanatomical studies of autism and ADHD reveal a complex and sometimes contradictory picture. Inconsistent findings may be due in part to small sample sizes and variable analytic approaches, which limit reproducibility and increase the risk of statistical false positives ^12–14^. Earlier studies relied on lower-resolution, coarser atlases, such as Desikan-Killiany (DK) atlas, and reported hemispheric averages, potentially obscuring subtle, region-specific or lateralized effects ^8,15,16^.

In autism, large-scale studies report increased cortical thickness in frontal regions and cortical thinning in temporal areas, the cingulate, inferior frontal cortex, inferior temporal sulcus, and parietal lobules^15,17–23^. Additionally, alterations in asymmetry of cortical thickness and orbitofrontal surface areas have been described ^24^. Longitudinal evidence suggests an atypical developmental trajectory, with increased thickness in early childhood, followed by accelerated thinning but with partial normalization during adolescence^8^.

In ADHD, cortical alterations are generally more subtle and heterogeneous. Increased cortical thickness has been reported in the somatosensory cortex, anterior cingulate, anterior insula, pre-SMA, and occipital cortices during adolescence, with fewer changes observed in adulthood^25,26^. Meta-analysis reveals inconsistent convergence and divergence, with both autism and ADHD exhibiting cortical thinning in the right temporoparietal junction, but divergent effects in motor areas (thinning in ADHD and thickening in autism^27^.

## Curvature & cortical folding as markers of NDDs

Cortical curvature, expressed as inward folds (sulci) and outward folds (gyri), provides additional insight into atypical development. Curvature reflects the interplay of biological and mechanical forces shaping cortical expansion, synaptic pruning, and myelination^28,29^.

Alterations in sulcal and gyral morphology have been reported in autism and ADHD, particularly in regions supporting sensory, motor, auditory, and salience-related processing. Autism shows atypical folding patterns in the anterior insula, frontal operculum, and temporoparietal junction^30^, and increased gyrification in visual, sensorimotor, and frontotemporal cortices^19,20(p202),31^. ADHD reveals increased gyrification in the Rolandic operculum, superior temporal gyrus, fusiform, cuneus, and orbitofrontal regions, with some effects linked to stimulant exposure^32^. Together, these findings suggest cortical complexity and folding morphometry may differentiate NDDs. However, large-scale, systematic comparisons remain limited due to lack of harmonized, curated, processed data across clinically enriched and unenriched datasets.

### Current Gaps and Study Aims

The present study is the first to characterize both cortical curvature and probabilistic atlas cortical thickness morphometric signatures of autism and ADHD by leveraging multiple large-scale datasets processed through a harmonized and reproducible pipeline. Datasets include the Oregon-ADHD-1000, OHSU-ASD, Autism Brain Imaging Data Exchange I & II (ABIDE I & II), Healthy Brain Network (HBN), and the Adolescent Brain Cognitive Development (ABCD) study (Supplementary section 1.1). This integrated approach enables both clinical and normative developmental analyses and supports examination of autism, ADHD and transdiagnostic morphological patterns. Ultimately, through this approach, we aim to improve understanding of the structural neurobiological mechanisms underlying autism and ADHD.

### Methods and Materials

#### Participants

We analyzed participants with autism, ADHD, and age-matched controls from six large-scale, publicly available neuroimaging datasets: HBN^33^, Oregon-ADHD-1000 and OHSU-ASD ^34^, ABIDE I & II^35,36^, and ABCD^37^. Full dataset descriptions and inclusion criteria are provided in the Supplementary Materials (Section 1.1–1.2). Participants were characterized using formal psychiatric diagnosis based on DSM-IV-TR or DSM-5 criteria (see Supplement 1.1). To minimize heterogeneity and diagnostic confounds, individuals with psychiatric diagnoses other than autism or ADHD were classified as ‘clinical’ and excluded from both the harmonization process and subsequent ANCOVA analyses. The final sample included 9,647 individuals aged 5-64 years across 67 sites: 1,533 with ADHD, 1,080 with autism, and 7,034 controls without either diagnosis.

#### Study Design

This study employed a between-subjects mega-analysis design, integrating data from multiple cohorts to compare cortical morphology across three diagnostic groups: autism, ADHD, and controls. Our primary variables of interest were cortical thickness and cortical curvature, with surface area analyses reported separately in the Supplementary Materials (Section 3.3). Group comparisons were conducted while controlling for demographic covariates (age and sex).

Morphometric values were parcellated using the Human Connectome Project (HCP) multimodal parcellation^38^, comprising 360 symmetrical ROIs. To place these fine-grained structural measures in a broader functional context, each HCP ROI was mapped onto the large-scale networks defined in the Cole-Anticevic Brain-wide Network Partition (CAB-NP)^39^. This resting-state-drive partition provides a robust framework for interpreting ROIs within distributed brain networks.

In parallel, cortical thickness was also parcellated using the MIDB Network Precision Atlas^40^, which defines 15 large-scale functional networks, along with an ROI-level parcellation of 71 parcels (SupplementarylJ2.2.1). MIDB atlas was included because it is specifically optimized to capture individual variability in brain organization. This framework also enables assessment of whether structural alterations in autism and ADHD align with network-level systems implicated in social-cognitive, attentional, and sensorimotor processes, such as the recently defined Somato-Cognitive Action Network (SCAN)^41^. Additionally, MIDB-based analyses (Supplementary 2.2.1) were conducted as sensitivity tests for the HCP results and to provide an additional network-level perspective on diagnostic group differences.

### MRI Processing and Cortical Morphometry

All structural MRI data were processed using the standardized ABCD-HCP pipeline^37^ to ensure consistent cortical surface reconstruction, alignment, and feature extraction across datasets. Cortical morphometric measures (FreeSurfer ∼ 60K grayordinate outputs), including cortical thickness and curvature, were parcellated using the HCP atlas template in 360 ROIs^38^ (see Supplementary Materials, Figure S3) using Connectome Workbench tools, optimized for high-performance computing.

Following a previously described methodology^30^, we used the midthickness surface for cortical surface outcome measures in each ROI. The midthickness surface is calculated by determining the midpoint between the white and pial surfaces. Midthickness surfaces were utilized for registration to align individual surfaces to the atlas target (fsaverage atlas and fs_LR atlas) and to measure cortical surface area. The midthickness surface is often used in surface-based analyses as it provides a geometrically unbiased representation of the cortical surface, positioned midway between the white and pial surfaces. The midthickness surface allows for a more accurate representation of the overall cortical geometry, capturing information from both the inner and outer cortical boundaries. Sulcal depth is computed as the distance between each vertex on the midthickness surface and the closest vertex on the hull surface. The computation excludes distances where the deviation from the surface normal is 90° or more.

#### Data Aggregation and Harmonization

Following processing, we aggregated structural MRI data across six datasets: HBN, Oregon-ADHD-1000 and OHSU-ASD, ABIDE I, ABIDE II, and the ABCD study. The multi-site and multi-cohort nature of these datasets, inter-site variability, and non-biological sources of variance (e.g., scanner manufacturer, acquisition protocols) posed potential confounds. To mitigate these issues, we employed NeuroCombat^42,43^, a widely used harmonization tool that corrects for batch effects while preserving biological signals in cortical thickness and curvature across 360 ROIs. NeuroCombat uses an empirical Bayes framework to estimate and adjust for non-biological variance associated with data collection sites, referred to as “batches.” In our study, 67 site-specific batches were included across the six datasets (See supplementary information 2.3). As is standard practice, to preserve biologically meaningful variance, the harmonization model incorporated covariates including age, sex, autism diagnosis, and ADHD diagnosis, ensuring that batch adjustments did not remove effects related to these variables (see Suppl. 2.3).

To assess and visualize site-related variance before and after harmonization, we applied Principal Component Analysis (PCA) to cortical thickness and curvature features across the 360 ROIs (Figure 1).

**Figure 1.**
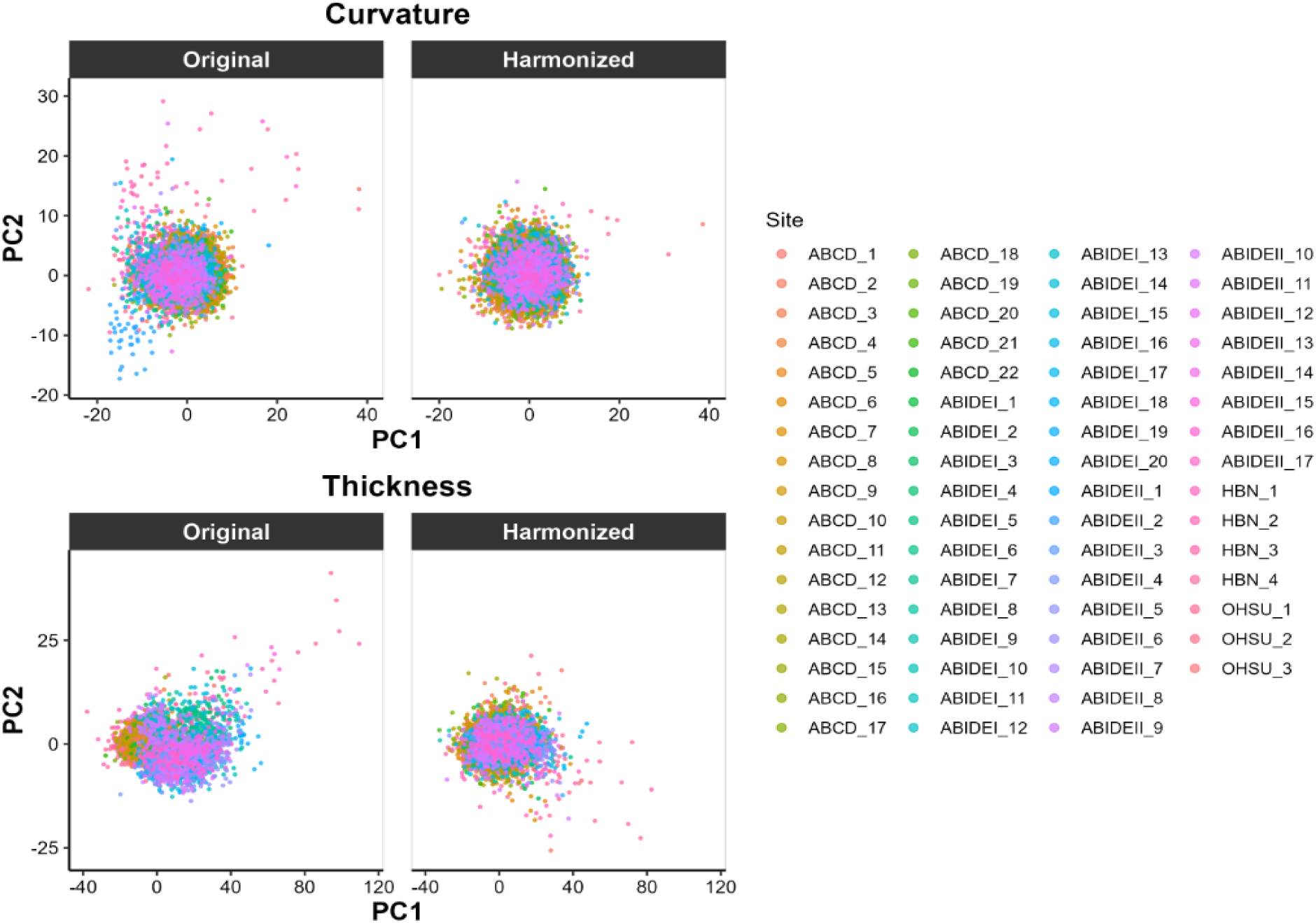
Principal Component Analysis (PCA) plots of cortical thickness and curvature before and after harmonization with NeuroCombat. Each point represents an individual, color-coded by the acquisition site. In the original data, cortical thickness shows site-related clustering, which is reduced after harmonization. Curvature shows less pronounced site clustering compared to thickness.

### Statistical Analysis

Following data harmonization, we examined diagnostic-group differences in cortical morphometry using region-wise Analyses of Covariance (ANCOVAs) for each of the ROIs. Separate sets of models were fit for cortical thickness and cortical curvature. In each model, the diagnostic group (Autism, ADHD, or control) was an independent factor, and age and sex were included as covariates to account for potential confounding effects.

To control the family-wise error rate across ROIs, we applied a Bonferroni correction to the significance threshold for the main effect of group. For ROIs that survived correction, we performed a post hoc pairwise t-test (Bonferroni adjusted) to identify group contrasts. Effect sizes (Cohen’s *d)* were calculated to quantify the magnitude and direction of these effects. These analyses were performed on the original (unharmonized) and harmonized datasets. Analyses on unharmonized data provided baseline group comparisons, whereas harmonized analyses allowed us to isolate biologically meaningful effects while reducing site-related variance.

## Results

### Harmonization of Cortical Thickness and Curvature Corrects Batch Effect

Below, we show data harmonization’s effects on cortical thickness and curvature from the HCP atlas parcellation. Results for cortical thickness at different parcellations (MIDB-parcel ROI and MIDB-parcel network), and for surface area within each HCP-parcel ROI, are shown in Supplementary Materials (section 3). To mitigate inter-site variability and improve data consistency, we applied NeuroCombat separately to each morphometric feature. Consistent with our hypothesis, site batch effects were markedly reduced following harmonization—most prominently for cortical thickness.

In the top row (pre-harmonization/original data), data points cluster by site, with clear separation, particularly for cortical thickness. After harmonization (bottom row), this site-specific clustering is substantially diminished for thickness, indicating effective removal of site effects. Curvature, by contrast, exhibited minimal site-driven clustering even before harmonization, suggesting that batch effects were less pronounced for this feature. Together, these findings confirm that NeuroCombat successfully attenuates site-related variance—especially in cortical thickness—thus improving cross-site comparability and setting a strong foundation for subsequent group-level analyses.

### Cortical Thickness and Curvature Differ in Autism, ADHD, and Control Groups

We hypothesized that autistic and ADHD individuals exhibit distinct differences in cortical thickness and curvature relative to controls. To test this, we performed ANCOVAs across diagnostic groups for both original and harmonized data, with age and sex included as covariates. The results showed widespread group differences across multiple cortical regions, which remained significant after applying a stringent Bonferroni correction for 360 cortical parcels (p < 0.00013). Pairwise comparisons and group-specific “brain signatures” (defined as regions showing distinctive morphological alterations within each diagnostic group), are shown in Figure 2 and detailed in Supplementary (section 3.1.2). Additionally, significantly different ROIs for each feature were mapped to their corresponding functional networks using the Cole-Anticevic Brain-wide Network (Figure 3).

**Figure 2.**
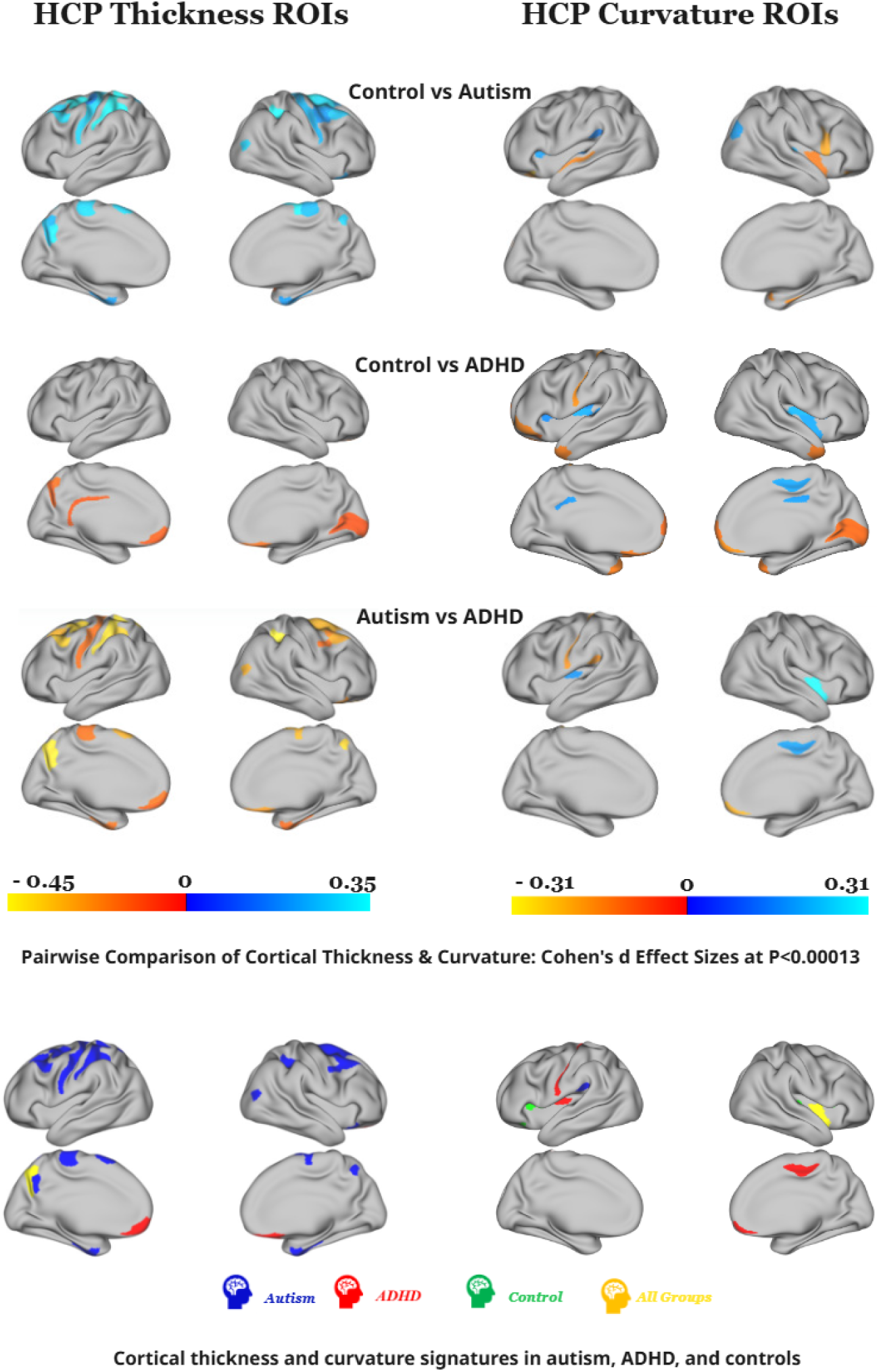
Cortical thickness and curvature alterations in autism, ADHD, and controls. Cohen’s d values in each pairwise comparison (top three rows) are shown for cortical thickness (left column) and curvature (right column). Warm colors indicate higher values in the second group, and cool colors indicate higher values in the first. The bottom row highlights group signature regions (blue = autism, red = ADHD, green = controls, yellow = transdiagnostic). Autism shows widespread cortical thinning relative, ADHD shows cortical thickening within DMN regions, and controls show intermediate thickness values. Curvature differences include a deeper PFcm sulcus (CO network) in autism and variations in alterations in ADHD, particularly deepened (Ig) and shallower (3a, 24dd) sulci, and a more prominent 10v gyrus.

**Figure 3.**
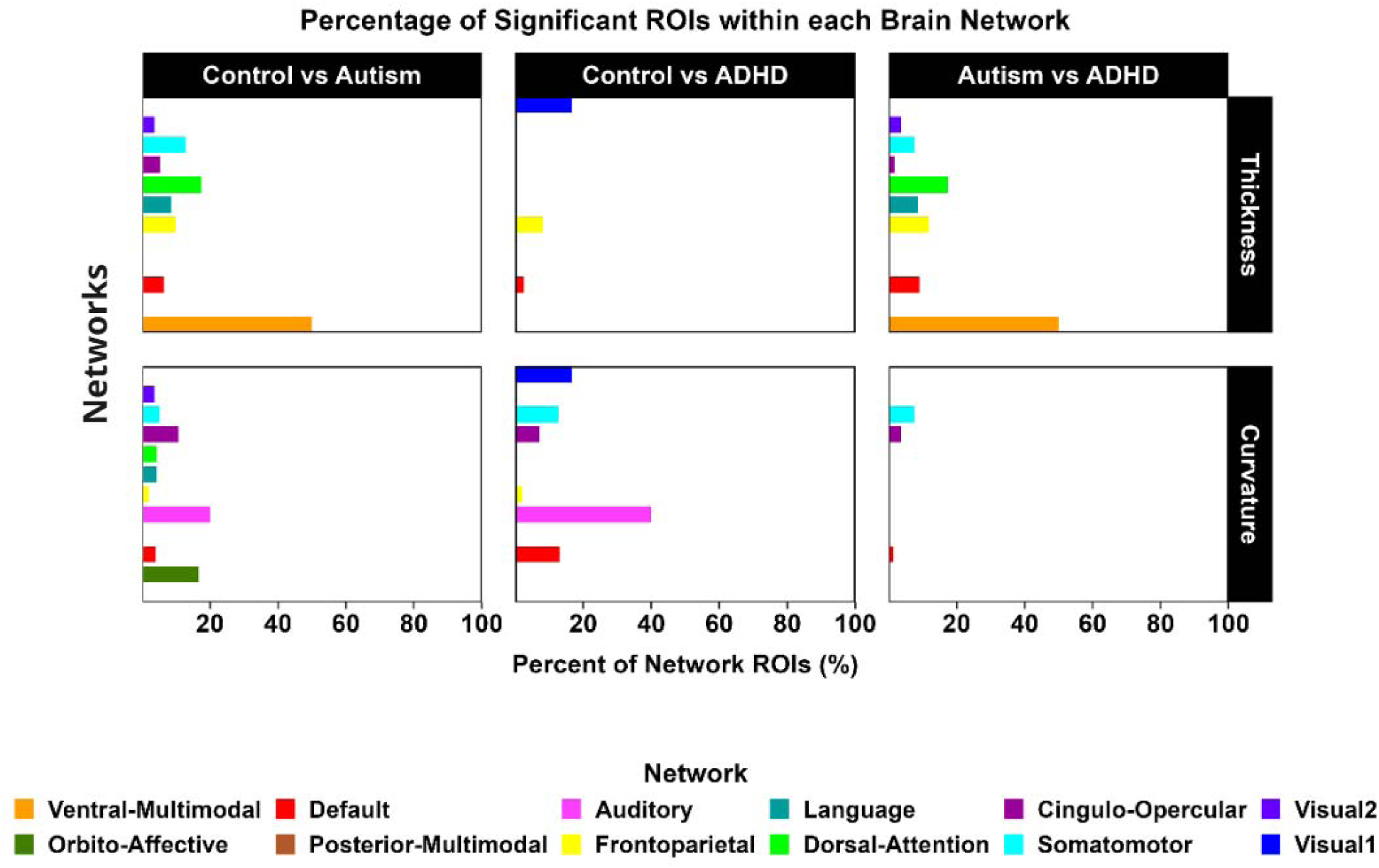
Network Distribution of Cortical regions showing group difference. Bar plots depict the percentage of ROIs within each network that showed significant group differences. Networks were defined using the Cole-Anticevic Brain-wide Network Partition (CAB-NP)^46^. For cortical thickness, the Ventral-Multimodal network showed the highest percentage of affected ROIs, while for curvature, the Auditory network was the most affected.

Unless otherwise noted, description of ROIs and their corresponding cortex in the following sections follows the HCP-MMP1.0 atlas guidance^38^.

### Cortical thickness signatures

#### Autism

Autistic participants showed a distributed pattern of cortical thinning across multiple large-scale networks compared to both control and ADHD. Cortical thinning was observed in Language (55b and SFL), Frontoparietal (7Pm, i6–8, and s6–8), Cingulo-Opercular (right 6ma), Somatomotor (area 4, 2, 6mp), DMN (8Av, 8Ad, 47s, 7m), Dorsal Attention (6a, AIP), Ventral Multimodal (PeEc) and Visual2 (LO3, VIP) networks. Anatomically, these regions are located within dorsolateral and premotor prefrontal cortices, mid-cingulate and supplementary motor areas, posterior cingulate, superior parietal and occipital cortices, and primary sensorimotor areas^38^ (detailed in supplementary section 3.1.2). Thinning in occipital regions (LO3 and VIP, parts of the visual 2 network) aligns with prior reports of altered visual integration^44,45^. Overall, these findings indicate widespread cortical thinning across networks supporting language, attention, sensorimotor, and visual processing.

#### ADHD

Structural analyses revealed ADHD-specific cortical thickening primarily within the DMN, including the right orbitofrontal cortex (OFC), left ventromedial prefrontal cortex (10v), and left anterior rostral prefrontal cortex (10r)^38^. Our findings are consistent with the evidence linking DMN dysfunction to impaired self-regulation, executive control, and altered integration of sensory, reward, and emotional inputs that shape reward-based decision-making^38,47,48^ (details in supplementary section 6-Box 1).

#### All groups

Across all diagnostic groups, significant cortical thickness differences emerged in the Fronto-Parietal Network, particularly in left POS2 and left area 7Pm, where autism showed the thinnest cortex and ADHD the thickest (Figure 2). These alterations suggest that fronto-parietal regions can represent a transdiagnostic locus of structural variation across autism, ADHD, and control groups. The left POS2 is a key integrative hub engaged in diverse cognitive operations^38^, while the left area 7Pm is implicated in complex cognitive processing and executive control^49^.

### Cortical Curvature Signature

#### Autism

Autism showed increased sulcal depth in the left PFcm sulcus (CO network) relative to ADHD and controls. This region is functionally related to language processing & sensory motor-related situated within the posterior opercular cortex^38^, consistent with prior reports of atypical sulcal morphology in autism^50–52^.

#### ADHD

In ADHD, we identified distinctive alterations within the somatomotor network, most prominently a shallower left area 3a sulcus relative to both autism and controls, alongside a deeper left Ig sulcus and a shallower right 24dd sulcus. Within the DMN, ADHD individuals also showed a more prominent right area 10v gyrus compared to the other groups.

#### Control

Controls exhibited unique sulcal curvature patterns across multiple network systems, with the most pronounced differences in the auditory, cingulo-opercular, and DMN networks. Specifically, we found more shallow sulci in right Area 52 and MBelt, as well as bilaterally in FOP3 and in the left FOP4 relative to both ADHD and autism. These regions support sensory integration, motor planning, and language-related^38^ processes.

In contrast, the left area 47m showed the opposite pattern, showing a deeper sulcus in controls, intermediate depth in ADHD, and a more shallow sulcus in autism across all pairwise group comparisons. Area 47m is notable for its heightened responsiveness to faces and social stimuli^38^.

#### All pairwise groups

PoI2, a posterior insular region affiliated with the Cingulo-Opercular (CO) Network, showed significant sulcal curvature differences across all pairwise group comparisons, with the deepest sulcus observed in the ADHD group. This region is functionally linked to motor tasks, tongue movement. However, known surface reconstruction artifacts in PoI2, particularly in the right hemisphere^38^, may impact this finding.

## Discussion

For the first time via a large-scale, multi-site morphometric study, we aimed to uncover unique cortical curvature and thickness brain signatures of autism and ADHD by aggregating, curating and harmonizing large clinically enriched and unenriched datasets. For methodological rigor and reproducibility, all MRI data were processed using the standardized ABCD-HCP pipeline, and inter-site variability was mitigated using NeuroCombat harmonization. All findings were mapped onto their network affiliations based on the Cole-Anticevic Brain-wide Network Partition^46^ to contextualize regional, structural findings within distributed functional networks.

Our cortical thickness comparisons are consistent with prior literature finding differences among autism, ADHD, and typical groups. Autism compared to typicals showed widespread cortical thickness reductions across temporal, cingulate, frontal, and parietal cortices consistent with prior findings^8,17–20,27,53–56^. Increased parietal and occipital thickness observed in ADHD compared to typical groups is consistent with prior findings^25^. Autism showed generally reduced cortical thickness compared to ADHD, consistent when synthesizing ADHD and ASD large-scale study findings^15,25,27,56^. Together, our findings support a growing literature emphasizing dissociable neurodevelopmental trajectories in autism and ADHD, particularly in networks supporting attentional control, executive function, and self-referential processing. The consistently thinner cortex in autism across these regions may contribute to its more pronounced cognitive and sensory processing impairments. We do not have the space to discuss here, a complete discussion of these findings can be found in the supplemental materials (section 3 & 4). Our curvature comparisons are somewhat consistent with prior studies comparing curvature between autism, ADHD, and typical groups. Autism showed altered curvature for frontal, temporal, parietal, and insula cortices compared to ADHD, consistent with prior reports^19,20,30,31,50,52,57,58^. Our observed widespread differences between ADHD and typicals, and ADHD and autism, have not been observed in prior studies^20,59–61^; this may reflect our additional power to detect small effects, given that our sample size is much larger than these prior studies. A complete discussion of these findings can be found in the supplemental materials.

Our unique curvature signatures reflect findings within the literature. A unique autism cortical signature was reflected in the CO network, consistent with prior evidence of atypical sulcal morphology and frontotemporal hypo-gyrification in autism^50–52^, while a unique ADHD cortical signature was found across motor and default mode systems, consistent with emerging evidence linking ADHD to atypical maturation in these same networks^32^. Further details contextualizing these differences can be found in the supplemental materials.

Collectively, our findings fit within a unified neurodevelopmental framework, where divergent neurodevelopmental trajectories can occur late into adolescence as described below.

### A Unifying Neurodevelopmental Framework: Shared Vulnerability and Divergent Trajectories

Our findings can be explained with the model of divergent developmental trajectories within the <u>hierarchical and</u> <u>protracted maturation of association cortex</u> (e.g., DMN, FPN)^7^. These evolutionarily expanded regions, which underlie cognition, social behavior, and self-regulation, remain plastic well into young adulthood due to delayed inhibitory maturation and extended synaptic remodeling. Furthermore, brain structure shows moderate associations with genetics^62,63^, supporting the morphogenetic hypothesis. Such prolonged developmental windows make them particularly vulnerable to genetic, molecular, and environmental perturbations^7^.

Consistent with this framework, we found transdiagnostic convergence in the Fronto-Parietal Network, where cortical thickness differs across all groups. This suggests a shared area of vulnerability in late-maturation association systems where each group follows a distinct maturational pathway. Within this shared vulnerability, autism and ADHD seem to follow extreme but opposing neurodevelopmental trajectories. Autism may involve early cortical overgrowth, accelerated thinning during adolescence, and relative stabilization in adulthood^8^ while ADHD shows evidence of delayed cortical maturation and prolonged cortical thickening^9–11^, characterized by relatively thicker cortex^15,25,26^. These opposite timing patterns, i.e., accelerated versus delayed refinement, align with the broader concept of atypical timing of synaptic pruning and myelination across neurodevelopmental conditions. Cortical thinning in autism has been interpreted as tissue loss, but may also reflect increased intracortical myelination, which shortens T1 relaxation times and reduces apparent gray matter thickness without actual neuronal loss^29^.

Curvature findings further illuminate these divergent morphogenetic mechanisms. The deeper PFcm sulcus in autism might suggest altered mechanical forces during development, such as atypical surface area expansion, dendritic arborization, or local axonal tension^29,64–66^. Conversely, some shallower sulci in ADHD may reflect reduced cortical tension or delayed mechanical folding, aligning with broader evidence of a maturational lag in somatomotor regions. These divergent maturational patterns within shared association networks highlight how differences in developmental timing might produce distinct cortical architectures.

Together, these morphometric patterns are best understood within a unifying neurodevelopmental framework. Autism and ADHD share a vulnerability within high-plasticity transmodal systems but diverge in the direction and timing of cortical maturation. The opposing profiles of cortical thickness and curvature observed here, accelerated refinement in autism and delayed specialization in ADHD, highlight maturational timing as a central principle of neurodevelopmental divergence. These findings underscore the need for longitudinal and multimodal studies to clarify how structural timing differences within association cortex translate into functional outcomes and behavioral phenotypes across neurodevelopment.

### Methodological Considerations and Limitations in Large-Scale Neuroimaging

The main limitation to this study reflects broader challenges in the field: lack of harmonized clinical and behavioral phenotyping across large data collections. Key diagnostic tools such as the Autism Diagnostic Observation Schedule (ADOS), the Conners Rating Scale for ADHD, and IQ scales are not consistently available across the datasets, precluding a comprehensive analysis of brain-behavior relationships. This limits the ability to interpret anatomical differences in the context of cognitive functioning, an important consideration given the association between cortical thickness and cognitive performance^67,68^. Medication status was also inconsistently available, preventing us from controlling for its potential effects. This reflects a broader issue, as many studies fail to implement dual diagnostic evaluations for autism and ADHD despite their frequent co-occurrence^69^.

Furthermore, our findings are not generalizable to individuals with the highest support needs, such as those with co-occurring intellectual disability, who remain profoundly underrepresented in neuroimaging research. This highlights a critical need for the development and widespread adoption of inclusive and accessible MRI protocols to ensure that future work is representative of the full neurodevelopmental spectrum.

Additionally, given that both autism and ADHD are highly heterogeneous conditions^70,71^, our group-level approach cannot capture the full range of neurobiological variability. Finally, while we implemented rigorous quality assurance, the absence of quality control metadata across all datasets remains a constraint on interpretability and replication. Addressing these issues by establishing standardized phenotyping and QC frameworks in future data-sharing initiatives will be essential for advancing the field.

### Future Directions

One should remain cautious when interpreting the cortical signatures observed here. Both autism and ADHD may reflect a combination of strengths and weaknesses^53,72^, and this study does not elucidate how parts of these signatures may reflect behavioral strengths and/or weaknesses. The field should move beyond deficit-focused models toward identifying strength-based associations, such as brain associations with enhanced visuospatial skills in some autistic individuals^72^. By recognizing that every individual possesses unique capacities, future studies may pave the way for precision medicine that amplifies inherent strengths and supports weaknesses.

The most crucial future direction, however, is the implementation of large-scale autism and ADHD focused longitudinal designs. By tracking cortical maturation trajectories over time, we can test the proposed models of atypical developmental divergence in autism and ADHD more directly. Ultimately, the clinical utility of these findings lies in their potential to guide personalized interventions. Our findings align with the embodied cognition framework, suggesting that interventions engaging motor systems may enhance language and social cognition^73,74^. Additionally, this study provides a harmonized, large-scale morphometric dataset processed using standardized pipelines. We are preparing a companion manuscript to make these curated and harmonized data publicly available to the research community to promote transparent, reproducible, and multi-sites neuroimaging research and advance science in neurodevelopmental and clinical conditions.

## Supporting information

This supplementary document provides the expanded technical framework and detailed results for our study

## Acknowledgments

We thank our colleagues at Oregon Health & Science University (OHSU) for their assistance in providing additional information and support throughout the study. We also extend our gratitude to our collaborators at the Child Mind Institute for sharing additional details related to the HBN dataset, which greatly contributed to this research. We are also deeply grateful to Theresa Vaughan at the National Center for Adaptive Neurotechnologies for her invaluable guidance and mentorship throughout this project. This work was supported by the National Institutes of Health (NIH) grant K23 DA057486 (PI: Tervo-Clemmens) and National Institute of Biomedical Imaging and Bioengineering (NIBIB-NIH) grant P41EB018783 (PI: Wolpaw). Lastly, we acknowledge the use of artificial intelligence (AI) tools to enhance the clarity, organization, and writing style of this manuscript.

## References

1. Danielson ML, Claussen AH, Bitsko RH, et al. ADHD Prevalence Among U.S. Children and Adolescents in 2022: Diagnosis, Severity, Co-Occurring Disorders, and Treatment. J Clin Child Adolesc Psychol Off J Soc Clin Child Adolesc Psychol Am Psychol Assoc Div 53. 2024;53(3):343–360. doi:10.1080/15374416.2024.2335625

2. Maenner MJ, Warren Z, Williams AR, et al. Prevalence and Characteristics of Autism Spectrum Disorder Among Children Aged 8 Years — Autism and Developmental Disabilities Monitoring Network, 11 Sites, United States, 2020. MMWR Surveill Summ. 2023;72(2):1. doi:10.15585/mmwr.ss7202a1

3. Shaw KA. Prevalence and Early Identification of Autism Spectrum Disorder Among Children Aged 4 and 8 Years — Autism and Developmental Disabilities Monitoring Network, 16 Sites, United States, 2022. MMWR Surveill Summ. 2025;74. doi:10.15585/mmwr.ss7402a1

4. Doernberg E, Hollander E. Neurodevelopmental Disorders (ASD and ADHD): DSM-5, ICD-10, and ICD-11. CNS Spectr. 2016;21(4):295–299. doi:10.1017/S1092852916000262

5. Lau-Zhu A, Fritz A, McLoughlin G. Overlaps and distinctions between attention deficit/hyperactivity disorder and autism spectrum disorder in young adulthood: Systematic review and guiding framework for EEG-imaging research. Neurosci Biobehav Rev. 2019;96:93. doi:10.1016/j.neubiorev.2018.10.009

6. Albajara Sáenz A, Septier M, Van Schuerbeek P, et al. ADHD and ASD: distinct brain patterns of inhibition-related activation? Transl Psychiatry. 2020;10(1):1–10. doi:10.1038/s41398-020-0707-z

7. Sydnor VJ, Larsen B, Bassett DS, et al. Neurodevelopment of the association cortices: Patterns, mechanisms, and implications for psychopathology. Neuron. 2021;109(18):2820–2846. doi:10.1016/j.neuron.2021.06.016

8. Zielinski BA, Prigge MBD, Nielsen JA, et al. Longitudinal changes in cortical thickness in autism and typical development. Brain. 2014;137(6):1799–1812. doi:10.1093/brain/awu083

9. Berger I, Slobodin O, Aboud M, Melamed J, Cassuto H. Maturational delay in ADHD: evidence from CPT. Front Hum Neurosci. 2013;7:691. doi:10.3389/fnhum.2013.00691

10. Shaw P, Eckstrand K, Sharp W, et al. Attention-deficit/hyperactivity disorder is characterized by a delay in cortical maturation. Proc Natl Acad Sci. 2007;104(49):19649–19654. doi:10.1073/pnas.0707741104

11. Shaw P, Malek M, Watson B, Sharp W, Evans A, Greenstein D. Development of Cortical Surface Area and Gyrification in Attention-Deficit/Hyperactivity Disorder. Biol Psychiatry. 2012;72(3):191–197. doi:10.1016/j.biopsych.2012.01.031

12. Collins FS, Tabak LA. Policy: NIH plans to enhance reproducibility. Nature. 2014;505(7485):612–613. doi:10.1038/505612a

13. Marek S, Tervo-Clemmens B, Calabro FJ, et al. Reproducible brain-wide association studies require thousands of individuals. Nature. 2022;603(7902):654–660. doi:10.1038/s41586-022-04492-9

14. Poldrack RA, Baker CI, Durnez J, et al. Scanning the horizon: towards transparent and reproducible neuroimaging research. Nat Rev Neurosci. 2017;18(2):115–126. doi:10.1038/nrn.2016.167

15. Bedford SA, Lai MC, Lombardo MV, et al. Brain-charting autism and attention deficit hyperactivity disorder reveals distinct and overlapping neurobiology. Biol Psychiatry. Published online August 14, 2024. doi:10.1016/j.biopsych.2024.07.024

16. Raznahan A, Toro R, Daly E, et al. Cortical anatomy in autism spectrum disorder: an in vivo MRI study on the effect of age. Cereb Cortex N Y N 1991. 2010;20(6):1332–1340. doi:10.1093/cercor/bhp198

17. Braden BB, Riecken C. Thinning faster? Age-related cortical thickness differences in adults with autism spectrum disorder. Res Autism Spectr Disord. 2019;64:31–38. doi:10.1016/j.rasd.2019.03.005

18. Hadjikhani N, Joseph RM, Snyder J, Tager-Flusberg H. Anatomical differences in the mirror neuron system and social cognition network in autism. Cereb Cortex N Y N 1991. 2006;16(9):1276–1282. doi:10.1093/cercor/bhj069

19. Pereira AM, Campos BM, Coan AC, et al. Differences in Cortical Structure and Functional MRI Connectivity in High Functioning Autism. Front Neurol. 2018;9:539. doi:10.3389/fneur.2018.00539

20. Zoltowski AR, Lyu I, Failla M, et al. Cortical Morphology in Autism: Findings from a Cortical Shape-Adaptive Approach to Local Gyrification Indexing. Cereb Cortex N Y NY. 2021;31(11):5188. doi:10.1093/cercor/bhab151

21. Bedford SA, Park MTM, Devenyi GA, et al. Large-scale analyses of the relationship between sex, age and intelligence quotient heterogeneity and cortical morphometry in autism spectrum disorder. Mol Psychiatry. 2020;25(3):614–628. doi:10.1038/s41380-019-0420-6

22. Boedhoe PSW, van Rooij D, Hoogman M, et al. Subcortical brain volume, regional cortical thickness and cortical surface area across attention-deficit/hyperactivity disorder (ADHD), autism spectrum disorder (ASD), and obsessive-compulsive disorder (OCD). Am J Psychiatry. 2020;177(9):834–843. doi:10.1176/appi.ajp.2020.19030331

23. van Rooij D, Anagnostou E, Arango C, et al. Cortical and Subcortical Brain Morphometry Differences Between Patients With Autism Spectrum Disorder and Healthy Individuals Across the Lifespan: Results From the ENIGMA ASD Working Group. Am J Psychiatry. 2018;175(4):359–369. doi:10.1176/appi.ajp.2017.17010100

24. Postema MC, van Rooij D, Anagnostou E, et al. Altered structural brain asymmetry in autism spectrum disorder in a study of 54 datasets. Nat Commun. 2019;10(1):4958. doi:10.1038/s41467-019-13005-8

25. Almeida Montes LG, Prado Alcántara H, Martínez García RB, De La Torre LB, Ávila Acosta D, Duarte MG. Brain Cortical Thickness in ADHD: Age, Sex, and Clinical Correlations. J Atten Disord. 2013;17(8):641–654. doi:10.1177/1087054711434351

26. Duerden EG, Tannock R, Dockstader C. Altered cortical morphology in sensorimotor processing regions in adolescents and adults with attention-deficit/hyperactivity disorder. Brain Res. 2012;1445:82–91. doi:10.1016/j.brainres.2012.01.034

27. You W, Li Q, Chen L, et al. Common and distinct cortical thickness alterations in youth with autism spectrum disorder and attention-deficit/hyperactivity disorder. BMC Med. 2024;22(1):92. doi:10.1186/s12916-024-03313-2

28. Mercadante AA, Tadi P. Neuroanatomy, Gray Matter. In: StatPearls. StatPearls Publishing; 2024. Accessed September 4, 2024. http://www.ncbi.nlm.nih.gov/books/NBK553239/

29. Natu VS, Gomez J, Barnett M, et al. Apparent thinning of human visual cortex during childhood is associated with myelination. Proc Natl Acad Sci U S A. 2019;116(41):20750. doi:10.1073/pnas.1904931116

30. Dierker DL, Feczko E, John R. Pruett J, et al. Analysis of Cortical Shape in Children with Simplex Autism. Cereb Cortex N Y NY. 2015;25(4):1042. doi:10.1093/cercor/bht294

31. Kohli JS, Kinnear MK, Fong CH, Fishman I, Carper RA, Müller RA. Local Cortical Gyrification is Increased in Children With Autism Spectrum Disorders, but Decreases Rapidly in Adolescents. Cereb Cortex N Y NY. 2018;29(6):2412. doi:10.1093/cercor/bhy111

32. Ghozy S, Meiza J, Morsy A, et al. How psychostimulant treatment changes the brain morphometry in adults with ADHD: sMRI Comparison study to medication-naïve adults with ADHD. Psychiatry Res Neuroimaging. 2025;349:111992. doi:10.1016/j.pscychresns.2025.111992

33. Alexander LM, Escalera J, Ai L, et al. An open resource for transdiagnostic research in pediatric mental health and learning disorders. Sci Data. 2017;4(1):170181. doi:10.1038/sdata.2017.181

34. Nigg JT, Karalunas SL, Mooney MA, et al. The Oregon ADHD-1000: A new longitudinal data resource enriched for clinical cases and multiple levels of analysis. Dev Cogn Neurosci. 2023;60:101222. doi:10.1016/j.dcn.2023.101222

35. Di Martino A, Yan CG, Li Q, et al. The autism brain imaging data exchange: towards a large-scale evaluation of the intrinsic brain architecture in autism. Mol Psychiatry. 2014;19(6):659–667. doi:10.1038/mp.2013.78

36. Di Martino A, O’Connor D, Chen B, et al. Enhancing studies of the connectome in autism using the autism brain imaging data exchange II. Sci Data. 2017;4(1):170010. doi:10.1038/sdata.2017.10

37. Feczko E, Conan G, Marek S, et al. Adolescent Brain Cognitive Development (ABCD) Community MRI Collection and Utilities. Published online July 11, 2021:2021.07.09.451638. doi:10.1101/2021.07.09.451638

38. Glasser MF, Coalson TS, Robinson EC, et al. A multi-modal parcellation of human cerebral cortex. Nature. 2016;536(7615):171–178. doi:10.1038/nature18933

39. Ji JL, Spronk M, Kulkarni K, Repovš G, Anticevic A, Cole MW. Mapping the human brain’s cortical-subcortical functional network organization. NeuroImage. 2019;185:35–57. doi:10.1016/j.neuroimage.2018.10.006

40. Hermosillo RJM, Moore LA, Feczko E, et al. A precision functional atlas of personalized network topography and probabilities. Nat Neurosci. 2024;27(5):1000–1013. doi:10.1038/s41593-024-01596-5

41. Gordon EM, Chauvin RJ, Van AN, et al. A somato-cognitive action network alternates with effector regions in motor cortex. Nature. 2023;617(7960):351–359. doi:10.1038/s41586-023-05964-2

42. Fortin JP. Jfortin1/neuroCombat_Rpackage. Published online November 30, 2023. Accessed April 30, 2024. https://github.com/Jfortin1/neuroCombat_Rpackage

43. Fortin JP, Cullen N, Sheline YI, et al. Harmonization of cortical thickness measurements across scanners and sites. NeuroImage. 2018;167:104–120. doi:10.1016/j.neuroimage.2017.11.024

44. Sapey-Triomphe LA, Boets B, Eylen LV, et al. Ventral stream hierarchy underlying perceptual organization in adolescents with autism. NeuroImage Clin. 2020;25:102197. doi:10.1016/j.nicl.2020.102197

45. Bölte S, Hubl D, Dierks T, Holtmann M, Poustka F. An fMRI-study of locally oriented perception in autism: altered early visual processing of the block design test. J Neural Transm. 2008;115(3):545–552. doi:10.1007/s00702-007-0850-1

46. Ji JL, Spronk M, Kulkarni K, Repovš G, Anticevic A, Cole MW. Mapping the human brain’s cortical-subcortical functional network organization. NeuroImage. 2018;185:35. doi:10.1016/j.neuroimage.2018.10.006

47. Bludau S, Eickhoff SB, Mohlberg H, et al. Cytoarchitecture, probability maps and functions of the human frontal pole. NeuroImage. 2014;93 Pt 2(Pt 2):260-275. doi:10.1016/j.neuroimage.2013.05.052

48. Ongür D, Ferry AT, Price JL. Architectonic subdivision of the human orbital and medial prefrontal cortex. J Comp Neurol. 2003;460(3):425–449. doi:10.1002/cne.10609

49. Baker CM, Burks JD, Briggs RG, et al. A Connectomic Atlas of the Human Cerebrum—Chapter 2: The Lateral Frontal Lobe. Oper Neurosurg. 2018;15(Suppl 1):S10–S74. doi:10.1093/ons/opy254

50. Levitt JG, Blanton RE, Smalley S, et al. Cortical sulcal maps in autism. Cereb Cortex N Y N 1991. 2003;13(7):728-735. doi:10.1093/cercor/13.7.728

51. Nordahl CW, Dierker D, Mostafavi I, et al. Cortical folding abnormalities in autism revealed by surface-based morphometry. J Neurosci Off J Soc Neurosci. 2007;27(43):11725–11735. doi:10.1523/JNEUROSCI.0777-07.2007

52. Knaus TA, Tager-Flusberg H, Foundas AL. Sylvian fissure and parietal anatomy in children with autism spectrum disorder. Behav Neurol. 2012;25(4):327–339. doi:10.3233/BEN-2012-110214

53. Ecker C, Murphy D. Neuroimaging in autism—from basic science to translational research. Nat Rev Neurol. 2014;10(2):82–91. doi:10.1038/nrneurol.2013.276

54. Wallace GL, Dankner N, Kenworthy L, Giedd JN, Martin A. Age-related temporal and parietal cortical thinning in autism spectrum disorders. Brain. 2010;133(12):3745–3754. doi:10.1093/brain/awq279

55. Laidi C, Boisgontier J, de Pierrefeu A, et al. Decreased Cortical Thickness in the Anterior Cingulate Cortex in Adults with Autism. J Autism Dev Disord. 2019;49(4):1402–1409. doi:10.1007/s10803-018-3807-3

56. Richter J, Henze R, Vomstein K, et al. Reduced cortical thickness and its association with social reactivity in children with autism spectrum disorder. Psychiatry Res Neuroimaging. 2015;234(1):15–24. doi:10.1016/j.pscychresns.2015.06.011

57. Auzias G, Viellard M, Takerkart S, et al. Atypical sulcal anatomy in young children with autism spectrum disorder. NeuroImage Clin. 2014;4:593–603. doi:10.1016/j.nicl.2014.03.008

58. Sasabayashi D, Takahashi T, Takayanagi Y, Suzuki M. Anomalous brain gyrification patterns in major psychiatric disorders: a systematic review and transdiagnostic integration. Transl Psychiatry. 2021;11(1):1–12. doi:10.1038/s41398-021-01297-8

59. Forde NJ, Ronan L, Zwiers MP, et al. No Association between Cortical Gyrification or Intrinsic Curvature and Attention-deficit/Hyperactivity Disorder in Adolescents and Young Adults. Front Neurosci. 2017;11. doi:10.3389/fnins.2017.00218

60. Gharehgazlou A, Freitas C, Ameis SH, et al. Cortical Gyrification Morphology in Individuals with ASD and ADHD across the Lifespan: A Systematic Review and Meta-Analysis. Cereb Cortex. 2021;31(5):2653–2669. doi:10.1093/cercor/bhaa381

61. Gharehgazlou A, Vandewouw M, Ziolkowski J, et al. Cortical Gyrification Morphology in ASD and ADHD: Implication for Further Similarities or Disorder-Specific Features? Cereb Cortex. 2022;32(11):2332–2342. doi:10.1093/cercor/bhab326

62. Coffman C, Feczko E, Larsen B, et al. Heritability estimation of subcortical volumes in a multi-ethnic multi-site cohort study. Published online January 12, 2024:2024.01.11.575231. doi:10.1101/2024.01.11.575231

63. Elliott LT, Sharp K, Alfaro-Almagro F, et al. Genome-wide association studies of brain imaging phenotypes in UK Biobank. Nature. 2018;562(7726):210–216. doi:10.1038/s41586-018-0571-7

64. Essen DCV. A tension-based theory of morphogenesis and compact wiring in the central nervous system. Nature. 1997;385(6614):313–318. doi:10.1038/385313a0

65. Rash BG, Duque A, Morozov YM, Arellano JI, Micali N, Rakic P. Gliogenesis in the outer subventricular zone promotes enlargement and gyrification of the primate cerebrum. Proc Natl Acad Sci U S A. 2019;116(14):7089–7094. doi:10.1073/pnas.1822169116

66. Mota B, Herculano-Houzel S. BRAIN STRUCTURE. Cortical folding scales universally with surface area and thickness, not number of neurons. Science. 2015;349(6243):74–77. doi:10.1126/science.aaa9101

67. Menary K, Collins PF, Porter JN, et al. Associations between cortical thickness and general intelligence in children, adolescents and young adults. Intelligence. 2013;41(5):597–606. doi:10.1016/j.intell.2013.07.010

68. Schnack HG, van Haren NEM, Brouwer RM, et al. Changes in Thickness and Surface Area of the Human Cortex and Their Relationship with Intelligence. Cereb Cortex. 2015;25(6):1608–1617. doi:10.1093/cercor/bht357

69. Harikumar A, Evans DW, Dougherty CC, Carpenter KL, Michael AM. A Review of the Default Mode Network in Autism Spectrum Disorders and Attention Deficit Hyperactivity Disorder. Brain Connect. 2021;11(4):253. doi:10.1089/brain.2020.0865

70. Feczko E, Miranda-Dominguez O, Marr M, Graham AM, Nigg JT, Fair DA. The Heterogeneity problem: Approaches to identify psychiatric subtypes. Trends Cogn Sci. 2019;23(7):584–601. doi:10.1016/j.tics.2019.03.009

71. Lenroot RK, Yeung PK. Heterogeneity within Autism Spectrum Disorders: What have We Learned from Neuroimaging Studies? Front Hum Neurosci. 2013;7:733. doi:10.3389/fnhum.2013.00733

72. Menon V. Large-scale brain networks and psychopathology: a unifying triple network model. Trends Cogn Sci. 2011;15(10):483–506. doi:10.1016/j.tics.2011.08.003

73. Eigsti IM. A Review of Embodiment in Autism Spectrum Disorders. Front Psychol. 2013;4:224. doi:10.3389/fpsyg.2013.00224

74. Moseley RL, Pulvermüller F. What can autism teach us about the role of sensorimotor systems in higher cognition? New clues from studies on language, action semantics, and abstract emotional concept processing. Cortex. 2018;100:149–190. doi:10.1016/j.cortex.2017.11.019

