## Supplementary material for "Cortical Thickness and Curvature in Autism and ADHD: A Mega-Analysis": This supplementary document provides the expanded technical framework and detailed results for our study

This supplementary document provides the expanded technical framework and detailed results for our study. It details our complete methodology—from participant selection and neuroimaging processing to data harmonization and statistical modeling. While the main manuscript focuses on our primary findings for HCP-based cortical thickness and curvature, this supplement contains the full statistical outputs for all morphometric measures (including surface area). This includes ANCOVA results, Bonferroni-corrected pairwise comparisons, and Cohen's d effect sizes, which were omitted from the main text due to space constraints.

To establish the generalizability of our primary findings, we also present parallel analyses for cortical thickness using the MIDB atlas at both network (15 networks) and ROI levels (17 parcels). The complete diagnostic "signature" regions for autism, ADHD, and control groups are fully tabulated and functionally characterized. Finally, a dedicated supplementary discussion elaborates on the main manuscript findings. While these interpretations provide additional depth and context, they are less central to the primary narrative and are therefore presented here.

### 1. Descriptive Statistics

#### 1.1 Participants & Datasets Descriptions

Our study included 9,647 participants (ages 5–64) selected from six datasets: five clinical cohorts and one representing typically developing U.S. adolescents. The sample comprised 1,533 individuals with ADHD, 1,080 autistic individuals, and 7,034 controls (Table 1). Details for each dataset, including specific inclusion and exclusion criteria, are provided in the subsequent sections. Sex distribution across groups and datasets is shown in Figure S 1, and age distribution is shown in Figure S 2.

Table 1. Demographic characteristics of participants in the cortical thickness dataset

| Group | Sample | Mean Thickness | SD | Mean Age | SD Age | Male | Female |
| --- | --- | --- | --- | --- | --- | --- | --- |
| Control | 7034 | 3.01 | 0.45 | 12.07 | 4.12 | 3950 | 3084 |
| Autism | 1080 | 2.97 | 0.46 | 15.78 | 8.66 | 918 | 162 |
| ADHD | 1533 | 3.03 | 0.45 | 10.46 | 2.45 | 967 | 566 |
| Total | 9647 |  |  |  |  | 5835 | 3812 |

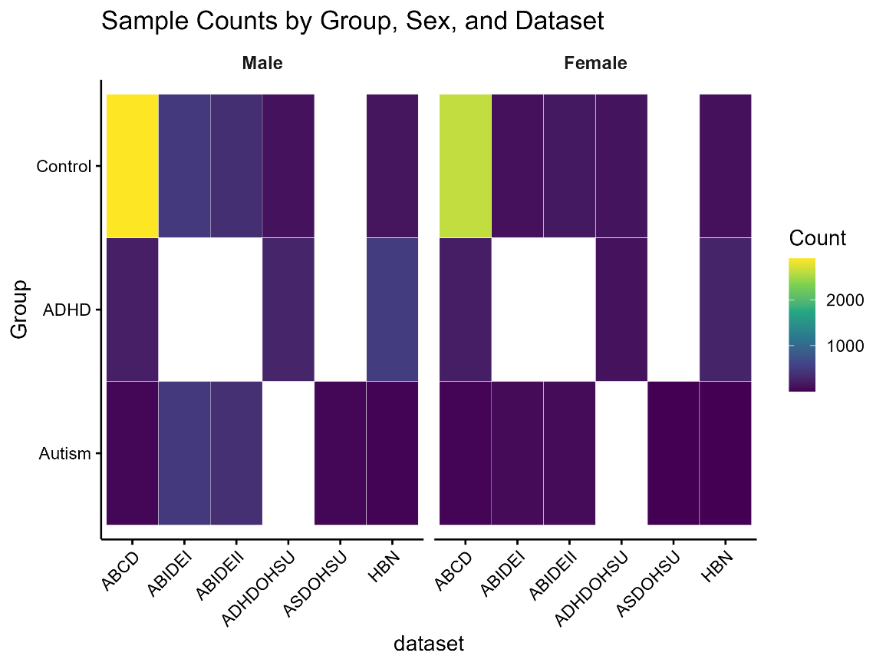

Figure S 1.Participants Number by Group, Sex, & Datasets

**Healthy Brain Networks (HBN).** This unique dataset has an enriched clinical population from over 3,000 children and young adults 5-21 years old from four sites in the US, measuring mental health outcomes from psychiatric, behavioral, to cognitive, and lifestyle phenotypes, in addition to biological measures including multimodal brain imaging, EEG, eye-tracking, voice and video recordings, genetics and actigraphy^1^.

We used the diagnosis labels provided for autism, ADHD, and control groups based on multi-step diagnostic procedure involving clinician-administered assessments, DSM-5 semi-structured interviews (KSADS-COMP), and targeted follow-up assessments (e.g., ADOS, CELF). All assessments are conducted by licensed clinicians, and checked by a second evaluator^1^.

**Adolescent Brain Cognitive Development (ABCD).** This large longitudinal dataset includes data from 9–11-year-old children collected across 22 United States sites. Here we used data from the ABCD Baseline dataset (release 2.0) with 11,873 children age 9-10 and a one-year follow-up including 4,951 participants age 10-11 ^2^. For this study, 6,000 participants were selected from the ABCD cohort, comprising two large, matched groups whose imaging data met high-quality control standards ^3^. ABCD measures and imaging data and phenotypes from multiple respondents (parent, youth, teacher) including clinical, educational, cultural, environmental, behavioral and gender, sexual and physical health as well as genetic data.

**Autism.** Autistic participants in the ABCD dataset were selected based on a single item from the ABCD screening survey for  parent-reported history of autism diagnosis. Parent reports are considered highly valid, as shown by SPARK researchers who found that ASD diagnosis was confirmed in 98.8% of cases using EMRs. Core clinical features recorded in EMRs were typical of autism samples and showed strong agreement with SPARK cohort data, further supporting the validity of clinical information in the SPARK database^4^.

Although the KSADS tool is available, it provides incomplete and inaccurate data for autism diagnosis, with release 5 containing incorrect and missing summary scores. Additionally, the only relevant diagnosis identified was “Other Specified Neurodevelopmental Disorder, Autism Spectrum Disorder full criteria not assessed (F88.0),” which does not reflect confirmed autism diagnosis and may include individuals with subthreshold presentations of autism.

**ADHD**. For ADHD diagnosis, the ABCD dataset lacks reliable measures, such as the Conners score. Further, despite the availability of the diagnosis for ADHD based on KSADS in ABCD, release 5 wiki (*ABCD Study*, n.d.) reports that data is incorrect. To address the limitations, we used the least stringent category of ADHD diagnosis, based on polygenic neuro risk factors study ^5^. This category (tier 1) meets *current* criteria of ADHD based on K-SADS-COMP (DSM-5) while exclude the participants who only meet *past* criteria for ADHD ^5^. Further, the ADHD prevalence in tier 1 is close to the national rate.

**Oregon-ADHD-1000 and OHSU-ASD cohorts.** The Oregon ADHD includes participants with ADHD (n = 739), controls (n = 434), and subthreshold/NOS presentations (n = 310). Longitudinal scans and extensive phenotypic data were collected, including clinical, cognitive, behavioral, temperament, and ADHD-related measures. Additionally, the OHSU-ASD cohort includes 109 autistic participants aged 7–16 years. For the present study, we included only baseline data and excluded longitudinal participants.

Diagnostic determinations in these datasets were made by two experienced clinicians (a child psychiatrist and a child psychologist) who independently reviewed all available materials, including parent and teacher reports, clinical interviews (K-SADS or SCID), and written observations from both the clinician and psychometrician. Participants with IQ scores below 75 were excluded^6^. We used the diagnostic labels provided by the dataset for autism, ADHD, and control groups, excluding those who did not meet full diagnostic criteria. Participants with dual diagnoses of ADHD and autism (due to small sample size), or with subthreshold/NOS classifications, were also excluded from our analyses. For ease of reference, we refer to these datasets as OHSU-ADHD and OHSU-ASD throughout the Supplementary Material.

**Autism Brain Imaging Data Exchange (ABIDE I & II)** ^7,8^. These datasets collect structural and resting-state functional MRI data from 2156 autistic and control individuals age matched (ABIDE I: 5-64) across 24 international sites.

We used the phenotypic diagnosis provided for autism and control groups. Diagnosis of autism was based on DSM-IV-TR criteria and determined by a clinician based on the Autism Diagnostic Observation Schedule (ADOS) and Autism Diagnostic Interview-Revised (ADI-R). Control groups were matched at the group level to autism for age, sex, handedness, and full-scale IQ, and had no self-reported history of ASD or any psychiatric or neurological condition ^7–9^.

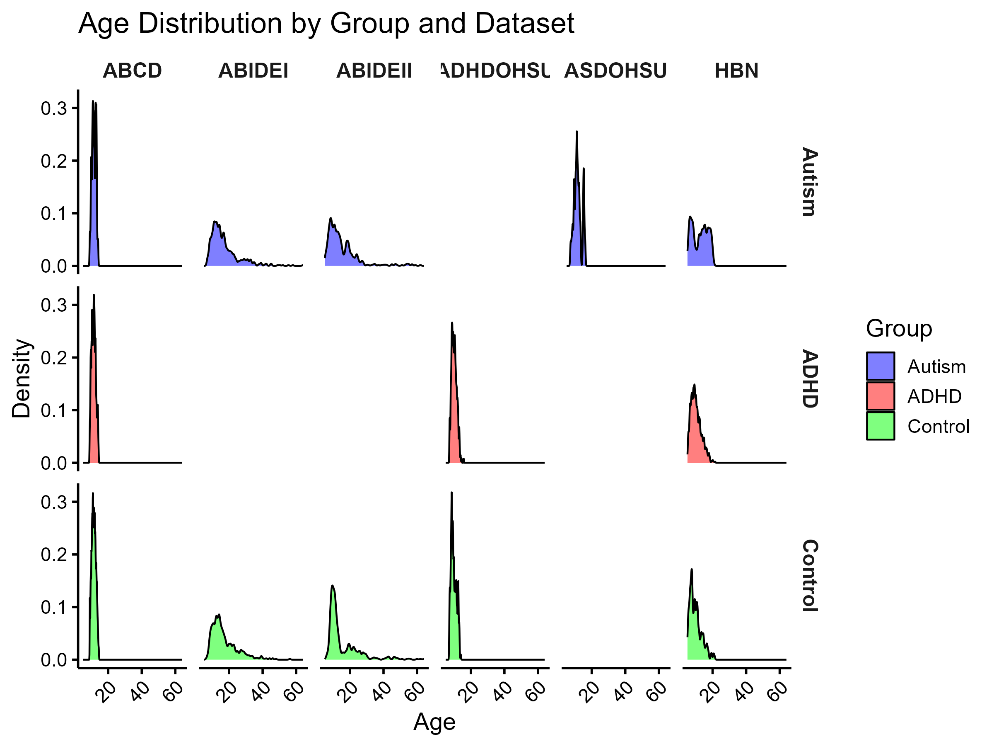

Figure S 2. Density Plots: Age Distribution By Group & Datasets

#### 1.2 Inclusion and exclusion criteria

Inclusion criteria. Our study includes individuals with confirmed diagnoses of autism or ADHD, as well as those classified as control (i.e., not diagnosed with autism or ADHD). Diagnosis labels provided within the datasets are utilized to identify eligible participants. We classified HBN participants into control, autism, ADHD, and a “clinical” group that includes all other clinical diagnoses. The age range of participants included in the study is not restricted, allowing for a broad representation of developmental stages.

Exclusion criteria. Our study excludes individuals categorized as subthreshold or not otherwise specified (NOS) for autism or ADHD, as determined by the diagnostic labels provided by the datasets. Participants with primary diagnosis of intellectual disability are excluded. Participants falling under this category are excluded from the study to ensure the homogeneity of the diagnostic groups. Further, we excluded subjects identified with both ADHD from polygenic study and autism from history item obtained from parent report in ABCD dataset due to the fact lack of such data in other datasets we used.

### 2. MRI Data Processing & Cortical Morphometry Features Computation

A schematic overview of the full analysis pipeline is provided in Figure S 3. Below we outline the key methodological steps.

#### 2.1 Neuroimaging Processing

All structural MRI datasets are processed using the standardized ABCD-HCP pipeline^3^, ensuring consistent preprocessing resources and parameters. This pipeline includes cortical surface reconstruction, alignment, and morphometric feature extraction, implemented via FreeSurfer and Workbench-compatible tools. The consistent application of this pipeline across datasets minimizes methodological variability and facilitates robust cross-sample comparisons.

#### *2.2* Cortical Morphometry Extraction

We computed three primary structural features for each region of interest (ROI): **Cortical Thickness** (measure of mean distance between the inner white matter surface and the outer pial surface within a ROI), **Cortical Curvature** (measure of mean curvature per ROI— quantifies the average degree of folding of the cortical sheet in each region, indicating the shape of gyri and sulci), and **Surface Area** (measure of the total expanse of the midthickness cortical surface for each ROI). Morphometric measurements were derived using Connectome Workbench commands, optimized for high-performance computing with parallelized SLURM workflows.

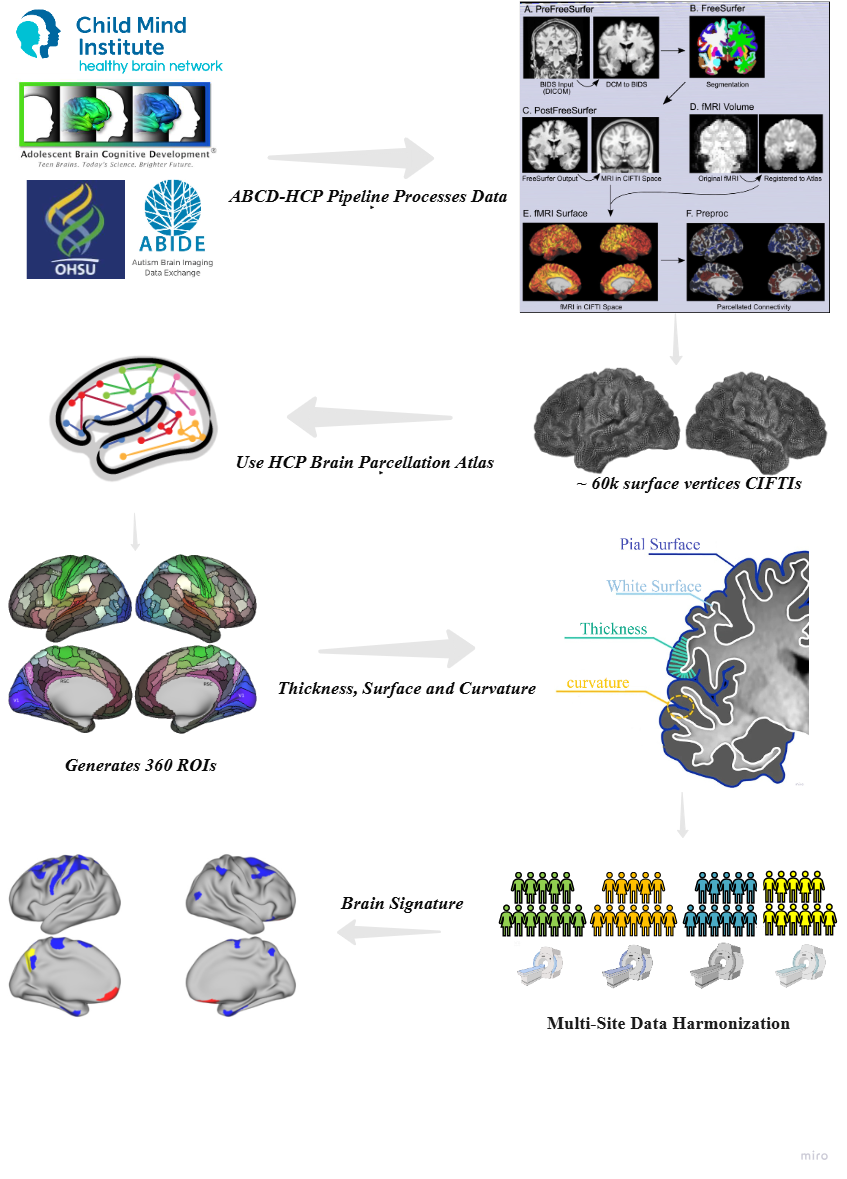

Figure S 3. HCP Cortical Morphometry features calculation workflow

Our workflow starts by aggregating data from multiple datasets: HBN, OHSU-ADHD, OHSU-ASD, ABIDE-1, ABIDE-2 and ABCD. Each dataset was processed using a standard FreeSurfer-based imaging pipeline (ABCD-HCP pipeline). The pipeline output, which consists of approximately 60,000 cortical grayordinates for each cortical morphometry feature, was then parcellated using a 360-region Human Connectome Project (HCP) template. This step gives thickness, surface area, and curvature values for each of the 360 regions of interest (ROIs) across the whole brain for every participant. To correct for batch (site) effects arising from the 67 data collection sites, the data were harmonized using the NeuroCombat tool. The final, harmonized data were then used to identify distinctive brain signature regions for autism and ADHD.

Morphometric measures were extracted using Connectome Workbench commands optimized for high-performance computing via parallelized SLURM workflows. Following the methodology of Dierker et al. (2015), cortical surface measures were computed on the midthickness surface, defined as the average position between the white and pial surfaces ^10^. Midthickness surfaces were utilized for registration to align individual surfaces to the atlas target (fsaverage atlas and fs_LR atlas) and to measure cortical surface area. The midthickness surface is often used in surface-based analyses as it provides a geometrically unbiased representation of the cortical surface, positioned midway between the white and pial surfaces. The midthickness surface allows for a more accurate representation of the overall cortical geometry, capturing information from both the inner and outer cortical boundaries. Sulcal depth is computed as the distance between each vertex on the midthickness surface and the closest vertex on the hull surface. The computation excludes distances where the deviation from the surface normal is 90° or more.

##### 2.2.1 Brain Atlas Templates

We utilized two cortical parcellation templates for morphometric analysis. The HCP atlas provided 360 cortical ROIs, from which cortical thickness, curvature, and surface area were computed. The MIDB atlas was used to compute cortical thickness only, at both the network level (15 large-scale networks) and ROI level (71 parcels). The inclusion of the MIDB parcellation served as a complementary analysis to evaluate the generalizability and robustness of observed group differences across atlases, and to examine whether specific networks (e.g., the SCAN network, relevant to social-cognitive processing in autism and ADHD) exhibit effects in different groups.

###### HCP Brain template

HCP-MMP1.0 Atlas ^11^. The Human Connectome Project Multi-Modal Parcellation (HCP-MMP1.0) defines 180 cortical areas per hemisphere (360 total ROIs) using high-resolution multimodal MRI from 210 healthy young adults (Figure S 4). Boundaries were delineated via a semi-automated, gradient-based approach applied to surface-aligned data (MSMAll), integrating cortical architecture, myelin content, functional connectivity, and task activation. Candidate borders were identified from spatial gradient “ridges” across multiple feature maps, verified by neuroanatomists, and named with reference to the neuroanatomical literature. Overall, 97 previously undescribed areas were defined alongside 83 established regions. The atlas demonstrates high accuracy, with a machine-learning classifier trained on multimodal “fingerprints” capable of automatically detecting cortical areas in new subjects with 96.6% accuracy, including in atypical configurations. This atlas provides a reproducible, anatomically grounded population parcellation optimized for surface-based analysis.

We selected the Human Connectome Project (HCP) parcellation due to its hemispherically symmetric, high-resolution (360-area) atlas derived from multimodal data. This template has finer anatomical precision and functional validity than coarser atlases such as Desikan–Killiany, which averages across broader gyral boundaries and is based on a unihemispheric model. Additionally, an important consideration for neurodevelopmental conditions is that hemispheric asymmetries are frequently observed, the HCP atlas preserves bilateral specificity, enabling sensitive detection of regional and lateralized differences.

Its derivation from independent structural and functional modalities also reduces the risk of overfitting because the parcellation is not tuned to our specific dataset but is grounded in reproducible anatomical landmarks and cross-validated in independent cohorts. The atlas further benefits from machine-learning–based boundary identification with 96.6% detection accuracy in new subjects, including atypical cortical configurations that this point supports its generalizability and reliability for morphometric analyses.

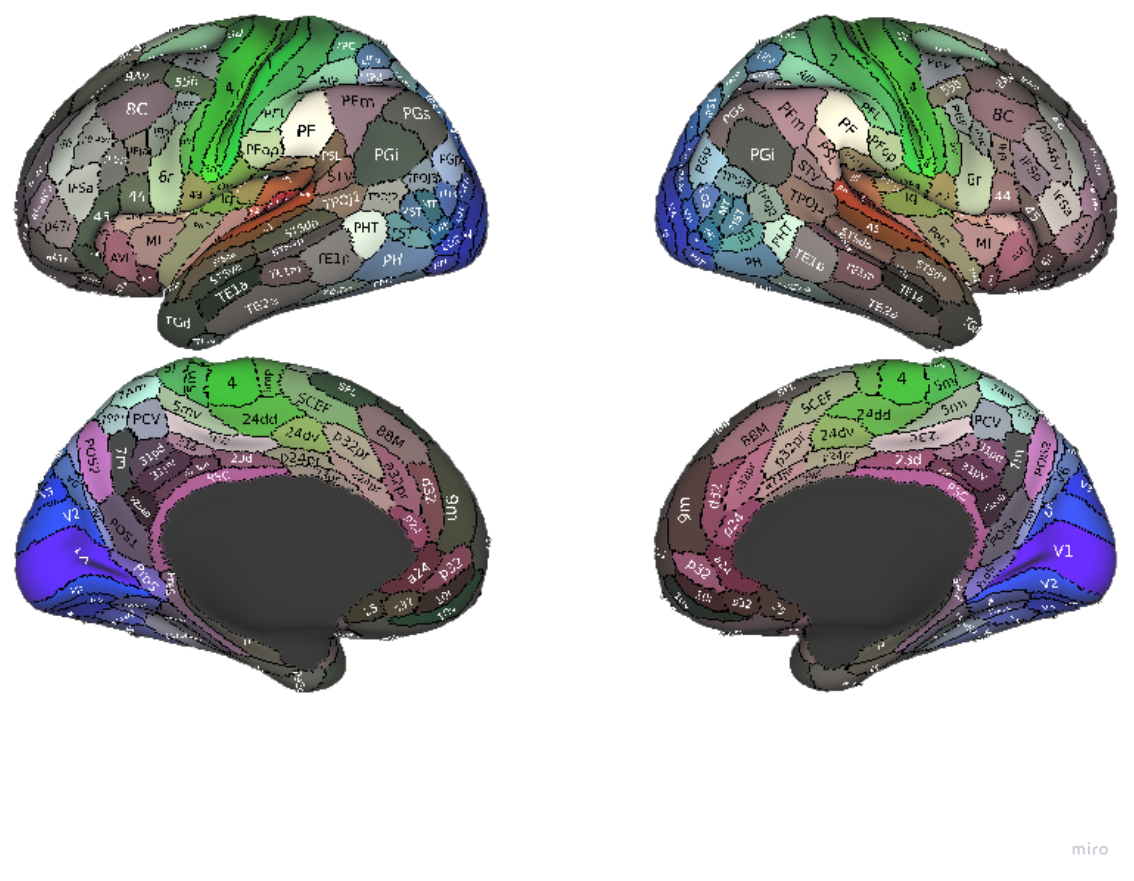

Figure S 4. HCP Brain Atlas Template

**Cole-Anticevic Network Parcellation** ^12^. While our analyses focus on structural cortical measures, we interpret the HCP ROI findings within a functional network framework to ground our results in the brain's systems-level organization. There is a well-established coupling between brain structure and function, wherein the anatomical architecture of the cortex is intrinsically organized to support dynamic functional specialization. By projecting our fine-grained, region-specific structural results onto the Cole-Anticevic Brain-wide Network Partition (CAB-NP), we can move beyond interpreting disparate regional effects. This approach provides further insights to ascertain whether the observed structural patterns conform to large-scale functional systems, providing a more integrated and neurobiologically meaningful understanding of how structural alterations relate to the brain's canonical network architecture in our target groups.

###### MIDB Brain template

**MIDB Brain Network template atlas**^13^. The Masonic Institute for the Developing Brain (MIDB) (Figure S 5) Precision Brain Atlas is a large-scale, open-source resource designed to capture individual variability in functional network organization. Built from over 53,000 individual-specific network maps in more than 9,900 participants from developmental cohorts (e.g., ABCD, Developing Human Connectome Project), it integrates precision resting-state fMRI mapping with probabilistic network templates. Networks were identified using multiple complementary methods—Infomap, template matching, non-negative matrix factorization, and the novel Overlapping MultiNetwork Imaging (OMNI) technique—enabling robust delineation of canonical networks and overlapping subnetwork zones.

The atlas delineates Default Mode (DMN), Visual (Vis), Frontoparietal (FPN), Dorsal Attention Network (DAN), Ventral Attention Network (VAN), Salience (Sal), Cingulo-Opercular (CO) network, Sensorimotor Dorsal (SMd), Sensorimotor Lateral (SMl), Auditory (Aud), Anterior Medial Temporal, Posterior Medial Temporal (post-MTL), Parieto-Occipital (PON), and Parietal Medial (PMN) networks. For our study, we use a slightly different MIDB template version that includes Somato-Cognitive Action (SCAN) ^14^ network, as well.

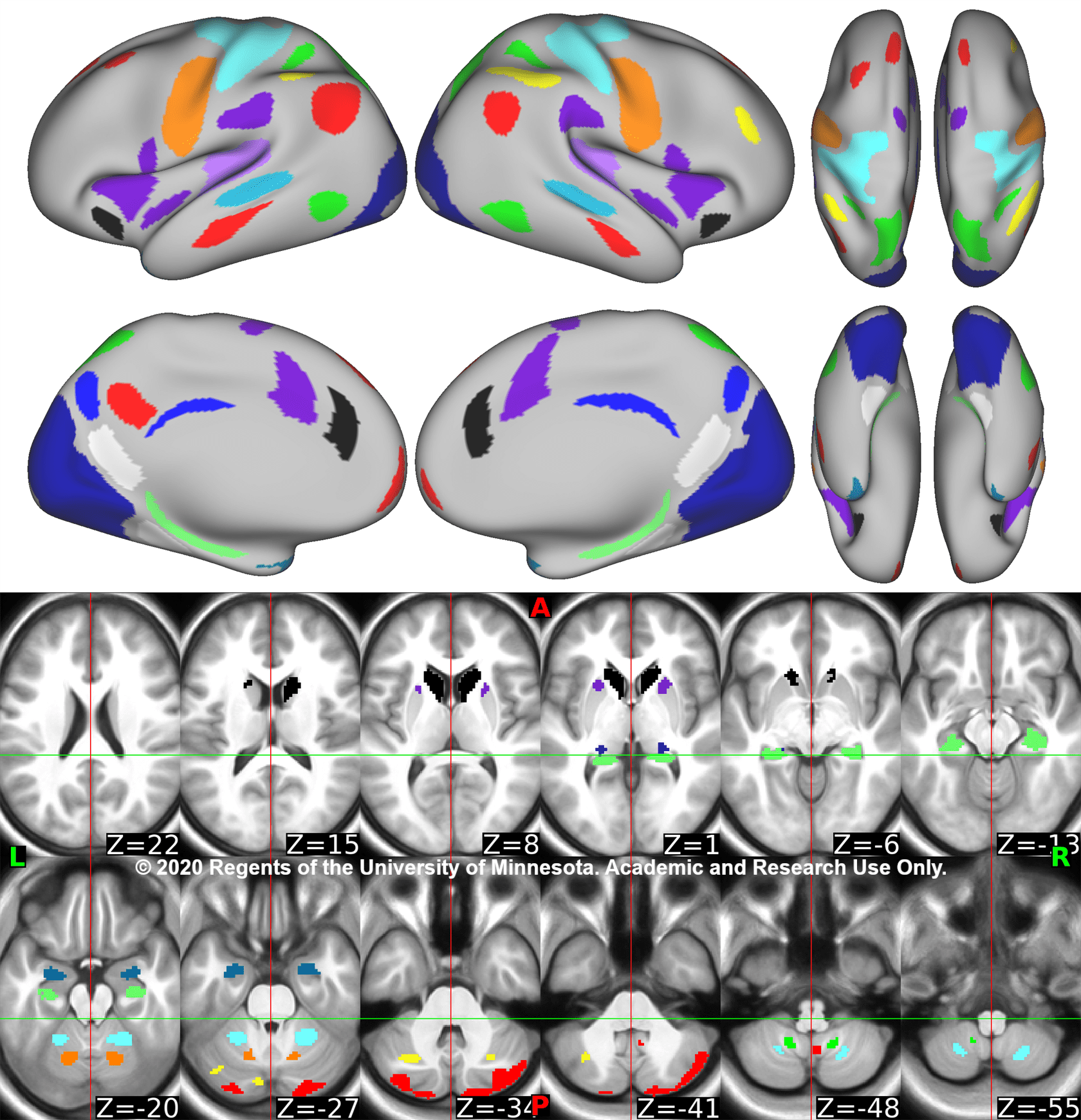

Figure S 5. MIDB Brain Atlas Template

**MIDB Brain ROI Atlas**. In addition to the 15-network probabilistic atlas, we created a 71-region-of-interest (ROI) parcellation from the MIDB Precision Brain Atlas to enable finer-grained analyses for this study. ROIs were generated using a patch-based refinement of the probabilistic network maps, starting from individual-specific parcellations aligned to the group-level template via the template matching procedure. Regions were defined based on spatially contiguous clusters of high network consensus (≥75% agreement across participants). Small, spatially isolated patches below a predefined minimum surface area threshold (30 vertices) were excluded to minimize noise and improve anatomical stability, with such vertices reassigned to the nearest large-scale network by geodesic proximity.

Left and right hemisphere cortical surfaces were processed separately to preserve hemispheric specificity, and volume data were excluded to produce a surface-only parcellation. Final ROI definitions were generated by combining left and right surface labels into a single CIFTI label file, resulting in 71 cortical ROIs that preserve the topographic fidelity of the MIDB probabilistic atlas while providing a resolution suitable for network-level and region-level morphometric analyses.

##### 2.2.2 Cortical Morphometry features Distribution

The distribution of cortical mean values across each feature is presented for the HCP brain template (Figure S 6) and the MIDB brain template (Figure S 7). In each figure, rows correspond to each cortical morphometry measures. For visualization of age-related trends in cortical values, only subjects within the shared age range were included. The first column of plots displays cortical values against age, stratified by autism, ADHD, and control groups. The second column presents these values separated by sex.

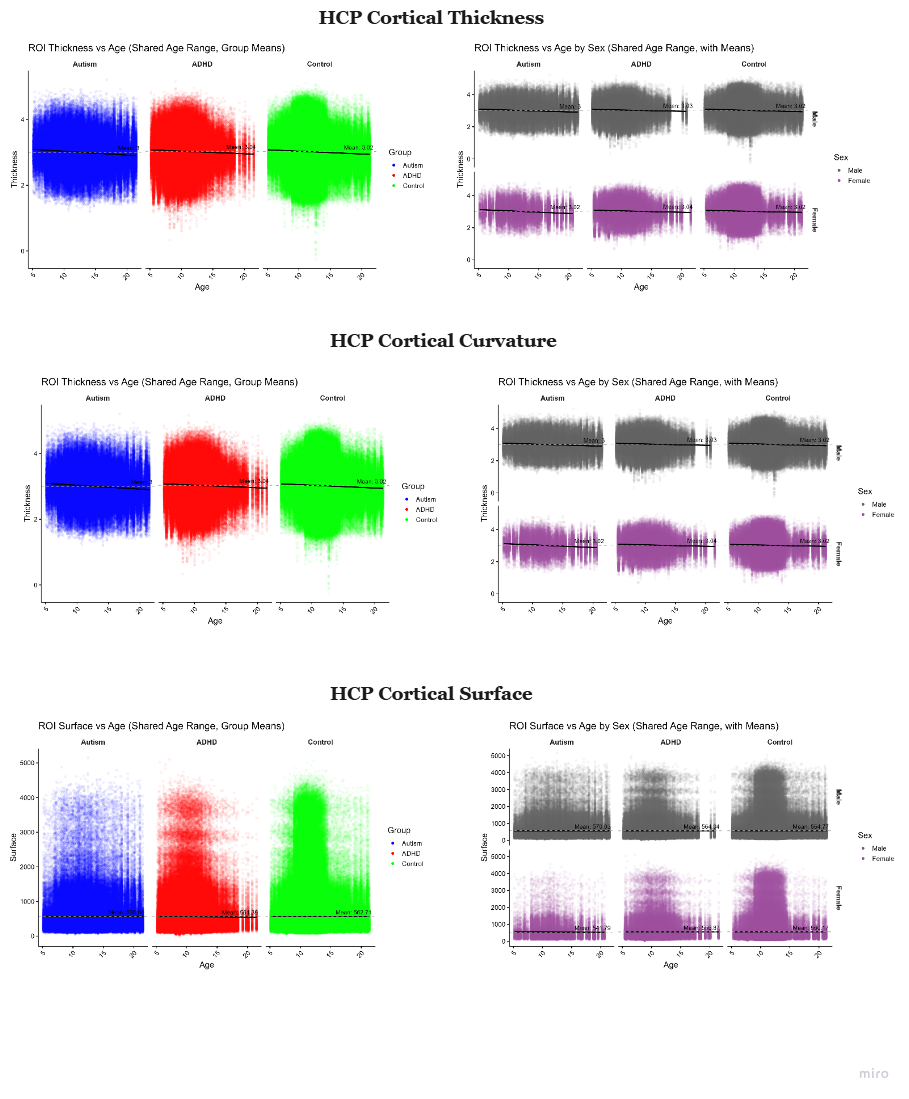

Figure S 6. HCP Cortical Morphometry Mean Values vs age by group | by Sex

Plots show cortical thickness (top row), cortical curvature (middle row), and surface area (bottom row) mean values across the shared age range for participants with autism (blue), ADHD (red), and controls (green). Thickness and curvature show a slight declining trend with age in both males and females, whereas surface area shows no noticeable age-related trend. The right-hand plots additionally present data separated by sex.

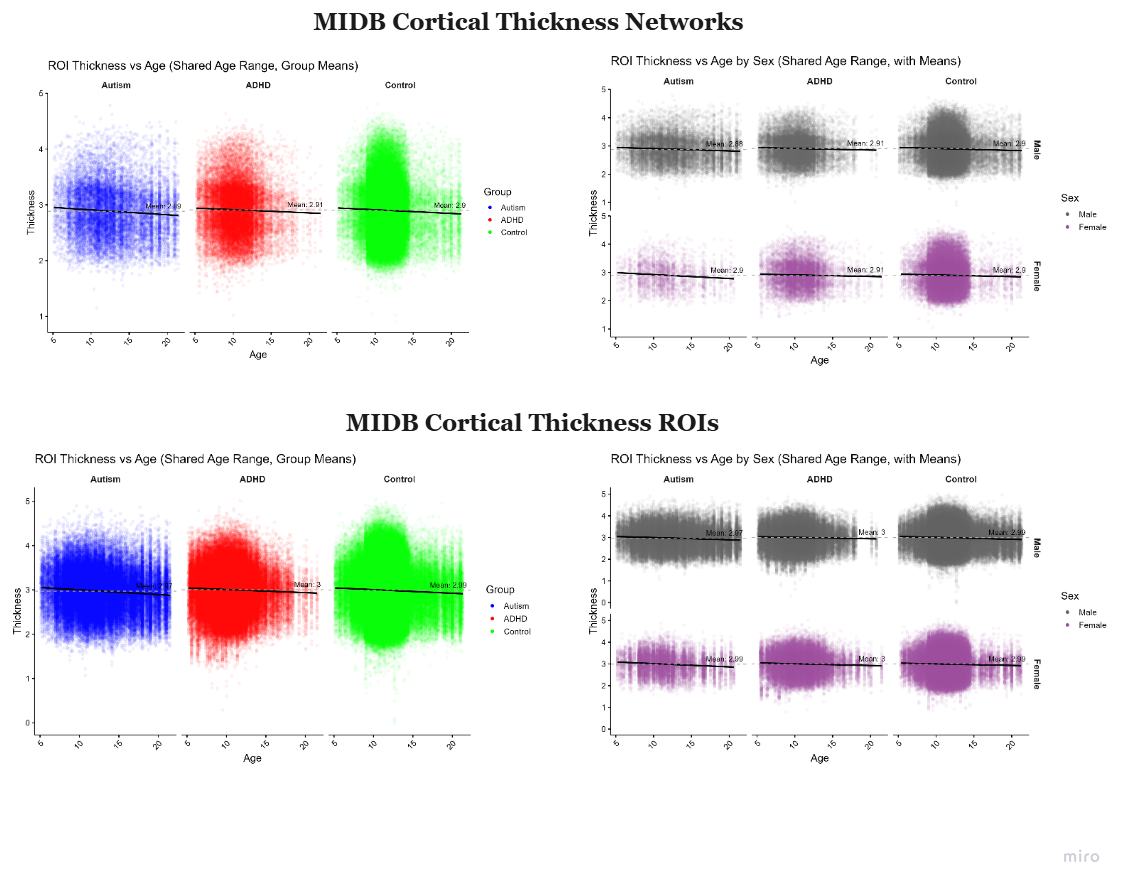

Figure S 7. MIDB Cortical Thickness Mean Values vs age by group | by Sex

Plots show cortical thickness mean values across the shared age range for participants with autism (blue), ADHD (red), and controls (green). The right-hand plots additionally display data separated by sex. Overall, cortical thickness shows a slight declining trend with age.

#### 2.3 Data harmonization

Data harmonization was performed to account for non-biological variability introduced by multi-site data acquisition. We utilized the NeuroCombat algorithm^15,16^, an established method for correcting batch effects in imaging data. The harmonization was applied to cortical morphometry features (mean cortical thickness, surface area, and curvature) for each of 360 predefined regions of interest (ROIs). Our combined dataset comprised 67 unique site-specific batches. A crucial aspect of this procedure was the preservation of variance attributable to biological factors. Therefore, the NeuroCombat design matrix was specified to include age, sex, and diagnostic status for both autism and ADHD as covariates, ensuring that statistical adjustments targeted site-related artifacts while leaving the variance associated with these key variables intact.

The harmonization followed the standard NeuroCombat pipeline:

1. Standardization of each ROI feature across subjects to ensure comparable scaling.
2. Estimation of batch effects via a location/scale (L/S) model with site-specific parameters (α: location, δ: scale), along with covariate effects (β).
3. Empirical Bayes estimation of prior distributions for site parameters to improve robustness in smaller sites.
4. Adjustment of individual feature values to remove estimated batch effects while retaining variance associated with biological covariates.

Formally, the harmonized values for each ROI ν\nuν, participant jjj, and site iii were computed using:  yComBatijν=yijν−αˆν−Xijβˆν−γ∗iνδ∗iν+αˆν+Xijβˆν ^15,16^ where yijνy_{ij}^{\nu}yijν is the original measurement, α^ν\hat{\alpha}^{\nu}α^ν and β^ν\hat{\beta}^{\nu}β^ν are the estimated intercept and covariate effects, and γiν∗\gamma^*_{i\nu}γiν∗, δiν∗\delta^*_{i\nu}δiν∗ are the empirical Bayes–estimated site-specific location and scale parameters ^15,16^.

By applying this harmonization framework, we substantially reduced non-biological variability while preserving variance attributable to diagnosis and demographics. This ensures greater reliability and generalizability in our group comparisons across diagnostic categories.

To visualize and qualitatively assess the effectiveness of the harmonization procedure, we performed Principal Component Analysis (PCA) on each cortical morphometry features (thickness, surface area, and curvature) across the 360 HCP regions (Figure S 8). The same was performed for cortical thickness across 15 networks and 71 ROIs (Figure S 9). PCA was applied separately to the unharmonized and harmonized data. In each case, we projected the high-dimensional data onto the first two principal components (PC1 and PC2), which capture the largest variance in the dataset. Prior to harmonization, the data showed clear clustering by sites—indicating strong site-related variance, particularly for cortical thickness. These PCA plots illustrate an intuitive low-dimensional summary of the harmonization impact across all morphometric features.

#### 2.4 Cortical Surface, Thickness & Curvature PCA Plots

In the following sections, PCA plots for cortical surface, thickness and curvature based on HCP atlas parcellations (Figure S 8) and cortical thickness in network and ROI parcels based on MIDB atlas (Figure S 9) before (first row) and after (second row) data harmonization are presented. Total of 67 acquisition sites were included across all datasets. For interpretability, site labels were renamed to reflect their corresponding dataset affiliations (e.g., HBN, ABIDE, OHSU, ABCD). The original site identifiers and their dataset mappings are provided in the following section (Table S 2). Some sites, such as OHSU, appear under multiple site names due to changes in MRI scanner models over time. As both PCA figures share the same legend, it is presented only once for clarity. Considering both figures have the same legend, we bring them only once.

#### 2.5 Mean Values for Significantly Different Regions

We present the mean values for each cortical morphometry feature (Figure S 10), focusing specifically on regions or networks that exhibited significant differences in at least one pairwise group comparison. For clarity, only the brain maps from the control group are displayed, as the subtle inter-group differences are not readily discernible in these visualizations.

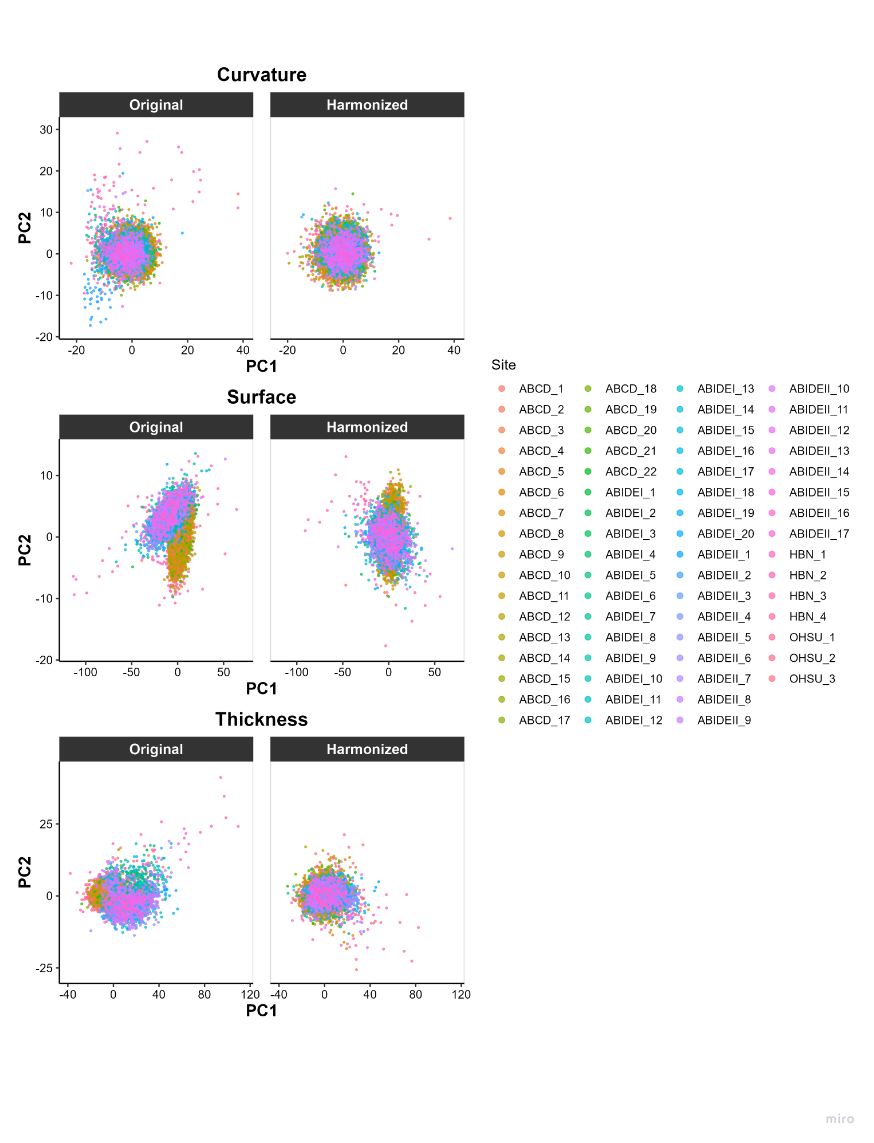

Figure S 8. Data Harmonization for HCP ROIs in Cortical surface, Thickness, and curvature

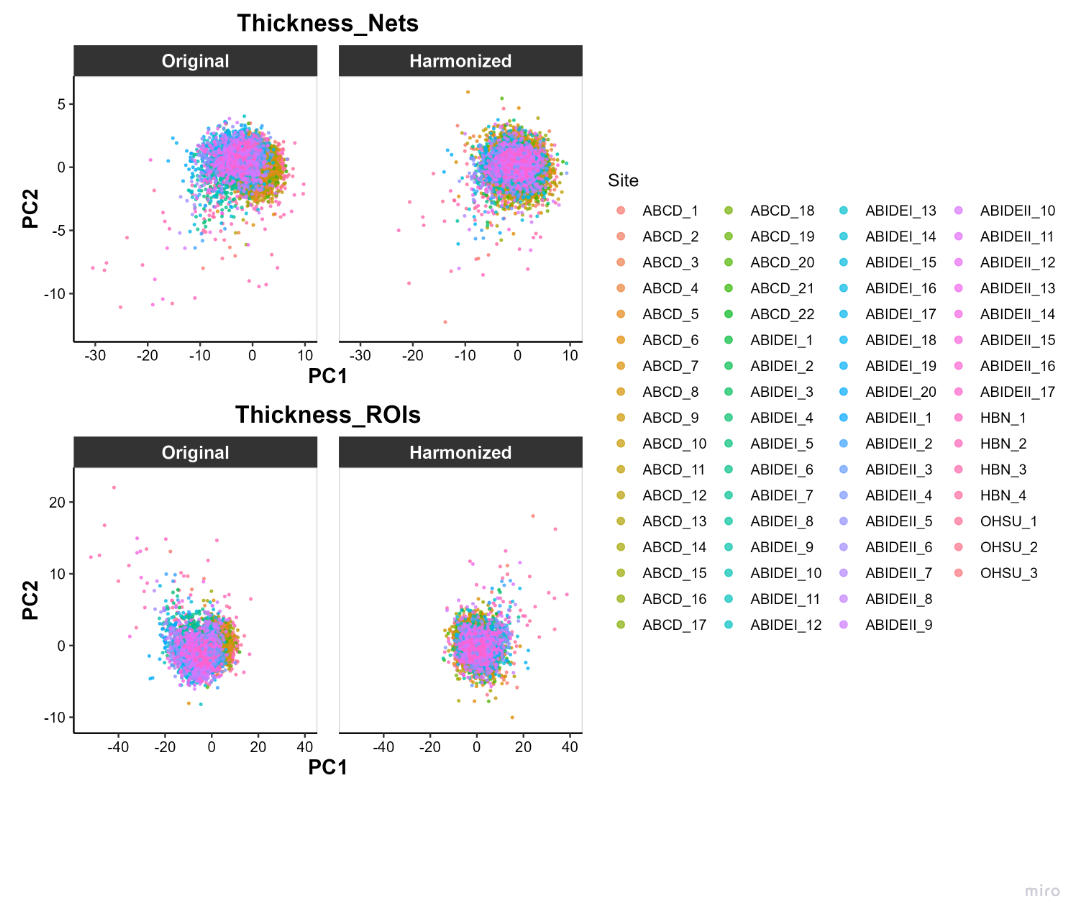

Figure S 9. Data Harmonization for MIDB Cortical thickness for network and ROI level parcels

Table S 2. Original Site names along with their renamed sites

| Datasets | Original Site Names | Renamed Sites | Datasets | Original Site Names | Renamed Sites |
| --- | --- | --- | --- | --- | --- |
| HBN | SI | HBN_1 | **abide1** | TRINITY | ABIDEI_5 |
| HBN | RUBIC | HBN_2 | **abide1** | UM_1 | ABIDEI_6 |
| HBN | CUNY | HBN_3 | **abide1** | UM_2 | ABIDEI_7 |
| HBN | CBIC | HBN_4 | **abide1** | USM | ABIDEI_8 |
| OHSUASD | TrioSingleBand | OHSU_1 | **abide1** | YALE | ABIDEI_9 |
| OHSUADHD | TrioSingleBand | OHSU_1 | **abide1** | CMU | ABIDEI_10 |
| OHSUADHD | Multi_Prisma | OHSU_2 | **abide1** | LEUVEN_1 | ABIDEI_11 |
| OHSUADHD | Single_Prisma | OHSU_3 | **abide1** | LEUVEN_2 | ABIDEI_12 |
| ABCD | site20 | ABCD_1 | **abide1** | KKI | ABIDEI_13 |
| ABCD | site12 | ABCD_2 | **abide1** | NYU | ABIDEI_14 |
| ABCD | site03 | ABCD_3 | **abide1** | STANFORD | ABIDEI_15 |
| ABCD | site11 | ABCD_4 | **abide1** | UCLA_1 | ABIDEI_16 |
| ABCD | site17 | ABCD_5 | **abide1** | UCLA_2 | ABIDEI_17 |
| ABCD | site06 | ABCD_6 | **abide1** | MAX_MUN | ABIDEI_18 |
| ABCD | site10 | ABCD_7 | **abide1** | CALTECH | ABIDEI_19 |
| ABCD | site21 | ABCD_8 | **abide1** | SBL | ABIDEI_20 |
| ABCD | site07 | ABCD_9 | **abide2** | ABIDEII-OILH_2 | ABIDEII_1 |
| ABCD | site16 | ABCD_10 | **abide2** | ABIDEII-GU_1 | ABIDEII_2 |
| ABCD | site04 | ABCD_11 | **abide2** | ABIDEII-SDSU_1 | ABIDEII_3 |
| ABCD | site19 | ABCD_12 | **abide2** | ABIDEII-OHSU_1 | ABIDEII_4 |
| ABCD | site02 | ABCD_13 | **abide2** | ABIDEII-BNI_1 | ABIDEII_5 |
| ABCD | site14 | ABCD_14 | **abide2** | ABIDEII-ETH_1 | ABIDEII_6 |
| ABCD | site13 | ABCD_15 | **abide2** | ABIDEII-TCD_1 | ABIDEII_7 |
| ABCD | site05 | ABCD_16 | **abide2** | ABIDEII-NYU_2 | ABIDEII_8 |
| ABCD | site01 | ABCD_17 | **abide2** | ABIDEII-NYU_1 | ABIDEII_9 |
| ABCD | site08 | ABCD_18 | **abide2** | ABIDEII-KKI_1 | ABIDEII_10 |
| ABCD | site09 | ABCD_19 | **abide2** | ABIDEII-USM_1 | ABIDEII_11 |
| ABCD | site15 | ABCD_20 | **abide2** | ABIDEII-IU_1 | ABIDEII_12 |
| ABCD | site18 | ABCD_21 | **abide2** | ABIDEII-IP_1 | ABIDEII_13 |
| ABCD | site22 | ABCD_22 | **abide2** | ABIDEII-KUL_3 | ABIDEII_14 |
| abide1 | PITT | ABIDEI_1 | **abide2** | ABIDEII-UCLA_1 | ABIDEII_15 |
| abide1 | OLIN | ABIDEI_2 | **abide2** | ABIDEII-EMC_1 | ABIDEII_16 |
| abide1 | OHSU | ABIDEI_3 | **abide2** | ABIDEII-UCD_1 | ABIDEII_17 |
| abide1 | SDSU | ABIDEI_4 |  |  |  |

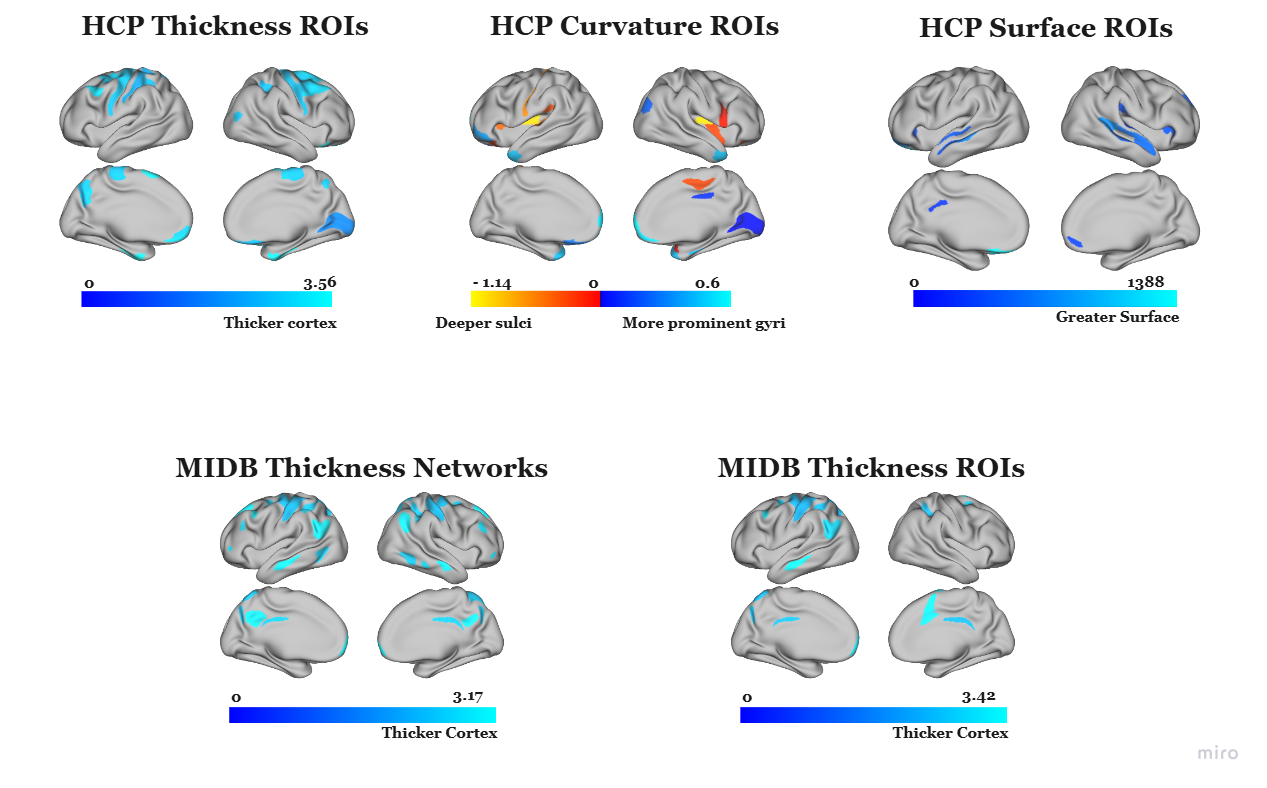

Figure S 10. Mean Values for Cortical Areas with Significant Pairwise Group Differences

Mean values are presented for ROIs that exhibited significant pairwise group differences. The color scale represents the mean value, with lighter colors indicating higher values. For cortical curvature, the yellow-orange spectrum denotes sulci (identified by negative values), while the blue spectrum represents gyri. Lighter shades within each spectrum correspond to deeper sulci or more prominent gyri. To ensure clarity and focus, these maps are derived solely from the control group data, as visual inspection of mean maps from other groups revealed no discernible qualitative differences.

### 3. Inferential Statistics: ANCOVA analysis

To investigate diagnosis-related variations in cortical morphology, we conducted our statistical analyses on two versions of the data independently: the original (unharmonized) and harmonized data to mitigate site-related variance (batch effects).

We performed a series of region-wise Analyses of Covariance (ANCOVAs) for each region of interest (ROI). These models tested for a main effect of diagnostic group (Autism, ADHD, or typically developing control) on morphometric features, while including age and sex as covariates. We applied a Bonferroni correction to the significance threshold (p<0.05) to control for the family-wise error rate across all comparisons. In regions where a significant main effect of group was found, we performed post-hoc pairwise t-tests with a Bonferroni adjustment to identify specific between-group differences. To quantify the magnitude of these effects, we calculated Cohen’s d effect sizes.

To give an overall view regarding these results within a functional framework, we aggregated the ROIs exhibiting significant differences according to their canonical network assignments. Figure S 11 presents box plots of these significant differences for cortical thickness (top row), surface area (middle row), and curvature (bottom row). Complementing this, Figure S 12 visualizes the distribution of Cohen’s d effect sizes for these same ROIs, which are mapped within each network by effect size.

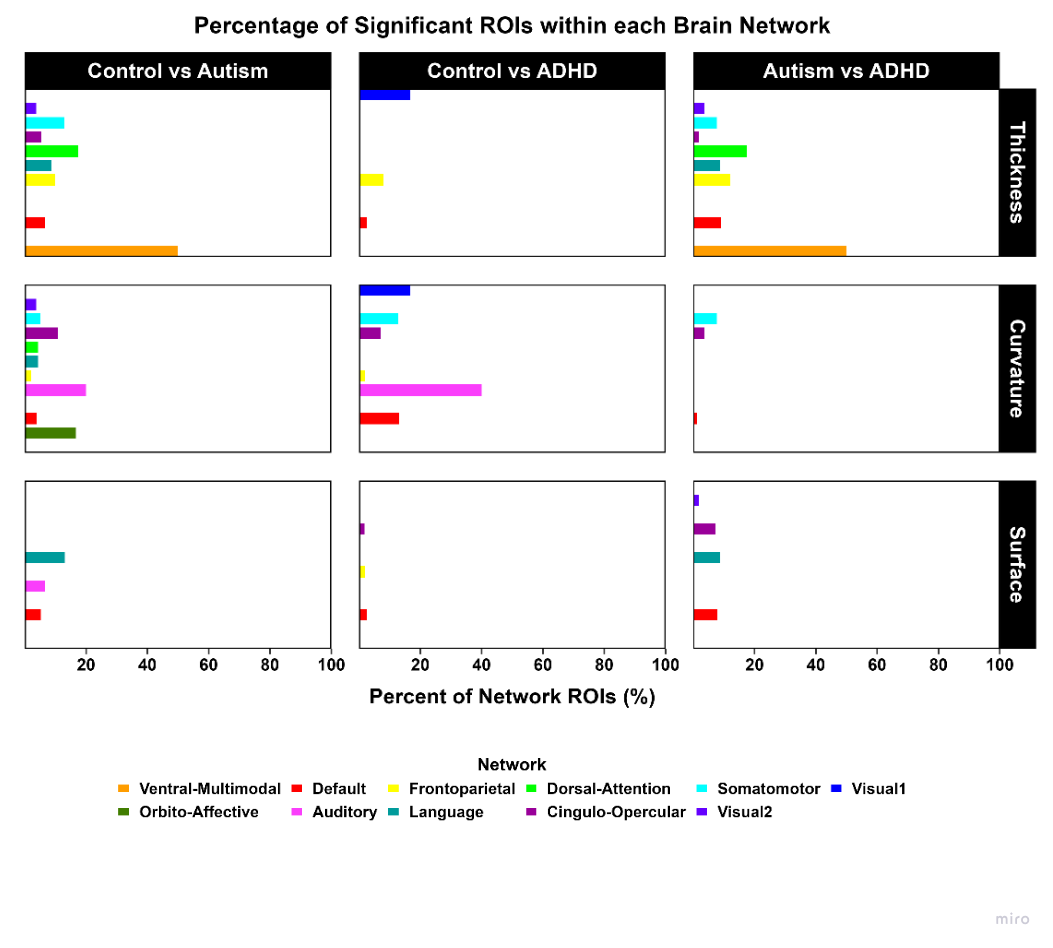

Figure S 11. Proportion of significantly altered ROIs across CAB-NP networks

Proportion of significantly altered ROIs across networks, mapped using the Cole-Anticevic Brain-wide Network Partition (CAB-NP)^12^. Rows correspond to cortical metrics (top = cortical thickness, middle = curvature, bottom = surface area), and columns correspond to pairwise group comparisons (Controls vs. Autism, Controls vs. ADHD, Autism vs. ADHD). The Ventral-Multimodal network exhibits the highest proportion of altered ROIs for cortical thickness, whereas the Auditory network contributed most for curvature.

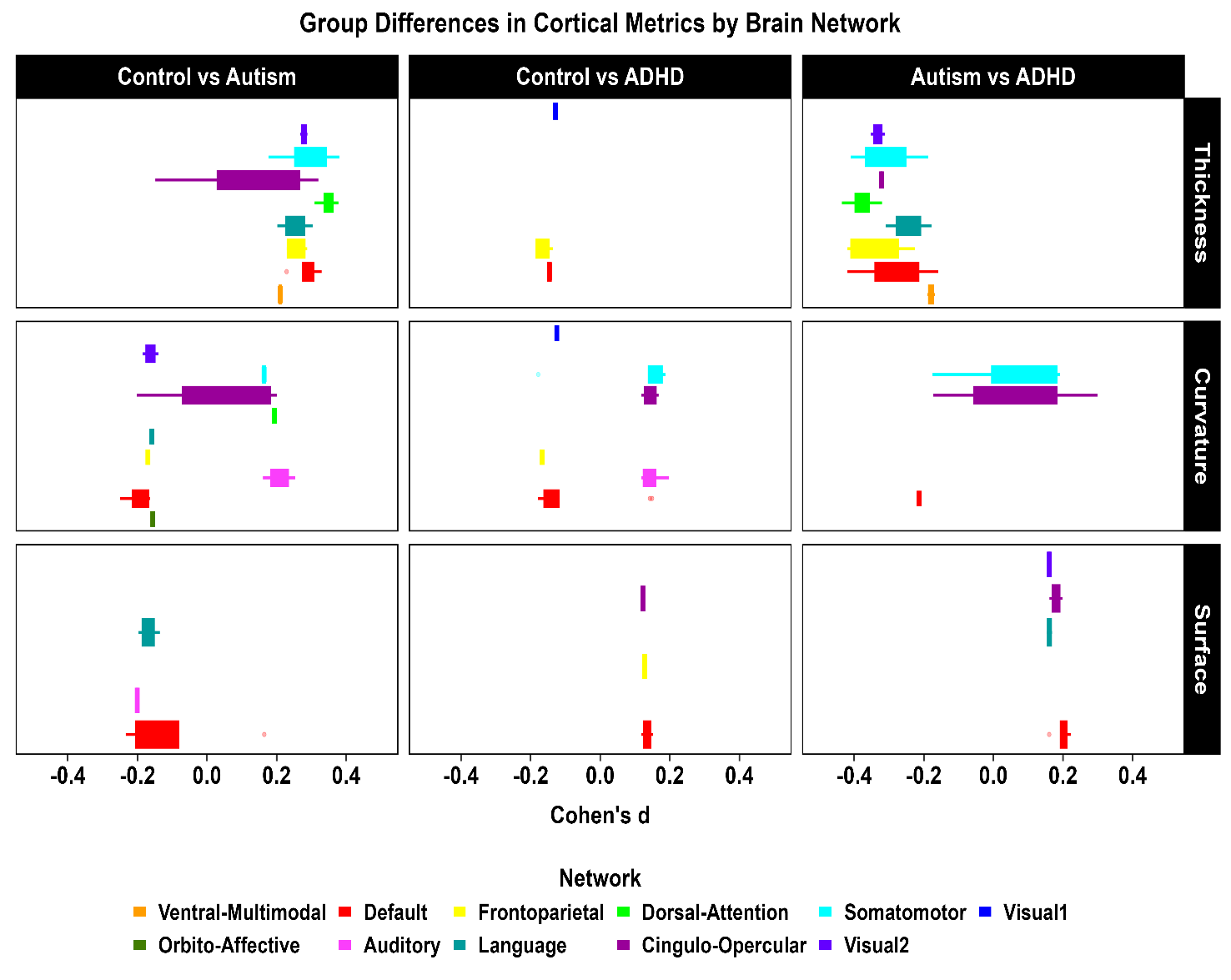

Figure S 12. Cohen's d Box plot for each HCP region with their corresponding HCP network system

This figure illustrates the magnitude and direction of group differences in cortical morphometry by showing the distribution of Cohen's d effect sizes. The effect sizes for all ROIs are grouped by their HCP functional network to reveal system-level patterns. The results are organized with rows corresponding to cortical thickness (top), curvature (middle), and surface area (bottom). The columns represent the three pairwise group comparisons: Autism vs. ADHD, Controls vs. ADHD, and Control vs. Autism.

Additionally, we generated cortical surface maps for each morphometric measure from both the HCP and MIDB atlases presented in the following sections. These figures display the log₁₀-transformed p-values for the main group-level effect (top row), followed by the corresponding Cohen’s d values for each pairwise group comparison (rows 2-4).

#### 3.1 HCP Cortical Thickness results in original and harmonized data

**In the following section, ANCOVA results for cortical thickness ROIs are presented, beginning with pairwise group comparisons, followed by a summary of signature regions—regions that significantly differ in one group and distinguish it from the other two.**

##### 3.1.1 HCP Cortical thickness pairwise group comparison

Our findings revealed that, on average, individuals with autism exhibited the thinnest cortex in the following regions, while those with ADHD showed the thickest cortical profiles relative to the other groups (Figure S 13). Below, we present significant group-level differences from the harmonized dataset across the 360 cortical regions of the HCP-MMP1.0 atlas. A summary table accompanies these results that outline each pairwise comparison, including the ROIs, their corresponding HCP functional networks, the range of Cohen’s d values (reported as absolute values), and the number of ROIs involved.

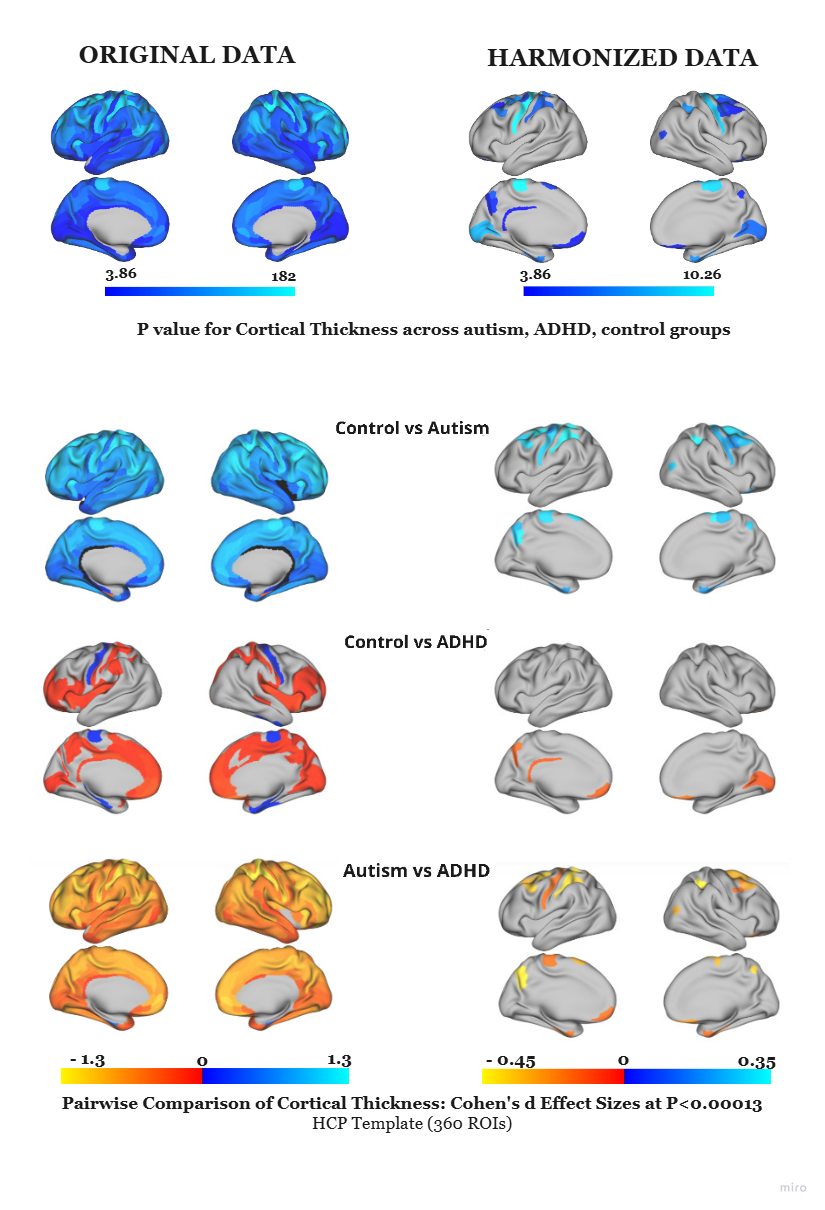

Figure S 13. HCP Cortical Thickness in Main Group and Pairwise Group Comparison

Results from ANCOVA are displayed, with the first row showing the main group effect as log_10_(p) values (p<0.05), corrected for comparisons). Subsequent rows present pairwise post-hoc comparisons for the significant regions, reported as Cohen’s d effect sizes. The first column of brain maps displays results from the original (non-harmonized) data, and the second column displays results after harmonization. Harmonization notably reduced false positives and inflated effect sizes, indicating its importance for robust cross-site analysis.

###### Control vs Autism

Compared to control participants, autistic individuals show significantly reduced cortical thickness across several functional networks (Table S 3). The most pronounced differences emerged in the DMN, then Fronto-Parietal (FPN); Somatomotor; and DAN have largest differences in this comparison. A few regions were also appeared in CO network; language; Ventral-Multimodal; Visual 2. Notably, R_PI was the only region exhibiting greater cortical thickness in autism relative to controls.

Table S 3. Significantly Different Cortical thickness ROIs in Control Vs, Autism comparison

| Network | n | | min_Cohens’ d | max_Cohens’ d | ROIs |
| --- | --- | --- | --- | --- | --- |
| DMN | 5 | | 0.228724 | 0.329284 | R_8Av, R_47s, L_7m, L_8Av, L_8Ad |
| FPN | 5 | | 0.228885 | 0.28807 | R_7Pm, R_i6-8, R_s6-8, L_POS2, L_7Pm |
| Somatomotor | 5 | | 0.176725 | 0.380443 | R_4, R_6mp, L_4, L_2, L_6d |
| DAN | 4 | | 0.309343 | 0.377227 | R_6a, R_AIP, L_6a, L_AIP |
| CO network | 3 | 0.148344 | | 0.319769 | R_FEF, R_6ma, R_PI |
| Language | 2 | 0.202053 | | 0.304019 | R_55b, L_SFL |
| Ventral-Multimodal | 2 | 0.208177 | | 0.212661 | R_PeEc, L_PeEc |
| Visual 2 | 2 | 0.268565 | | 0.288479 | R_LO3, L_VIP |

These widespread reductions in cortical thickness align with prior studies reporting thinning in autism across temporal, frontal, and parietal cortices^17–20^. Specific convergence was observed with previous reports of thinning in the inferior frontal cortex (IFC), inferior parietal lobule (IPL), and superior temporal sulcus (STS)—regions implicated in social cognition and emotion processing^21^. Reduced thickness in the left fusiform and inferior temporal sulcus has also been previously noted ^22^, as well as the right caudal anterior cingulate cortex^23^.

Our findings align with previous large-scale studies reporting reduced cortical thickness in autism across frontal, temporal, parietal, and occipital lobes, particularly within regions corresponding to the heteromodal association cortex (HASC)^24^. These areas, including the superior and middle frontal gyri, inferior parietal lobule, and superior temporal gyrus, are implicated in higher-order cognitive processes such as theory of mind (ToM), language, attention, and executive function. Notably, several of these regions overlap with key ToM-related hubs involved in social inference and self–other distinction, including the temporoparietal junction (TPJ), where we observed thinning consistent with meta-analytic findings^25^.

From a developmental perspective, our results also align with longitudinal evidence suggesting that autism is characterized by accelerated cortical thinning during mid-childhood and adolescence, particularly in frontal and parietal cortices^26^. This atypical trajectory—often interpreted as early cortical overgrowth followed by premature or excessive pruning—may underlie deficits in social and cognitive domains supported by these association areas^26^.

While cortical thinning is the dominant pattern observed in our findings, prior studies have reported increased cortical thickness in autism within specific regions, such as the superior temporal gyrus and inferior frontal sulcus^27,28^. These discrepancies may reflect regional heterogeneity, age-related variation, or methodological differences across studies.

###### Control vs ADHD

Our analysis revealed that individuals with ADHD exhibited significantly greater cortical thickness compared to controls in specific brain regions. These regions (Table S 4) are within the DMN, the FPN, and the primary visual network.

Table S 4. Significantly Different Cortical thickness ROIs in Control Vs, ADHD comparison

| Network | n | min_Cohens’ d | max_Cohens’ d | ROIs |
| --- | --- | --- | --- | --- |
| DMN | 3 | 0.139605 | 0.185474 | R_OFC, L_10r, L_10v |
| FPN | 3 | 0.135854 | 0.181974 | L_RSC, L_POS2, L_7Pm |
| Visual 1 | 1 | 0.12811 | 0.12811 | R_V1 |

This finding of localized cortical thickening is consistent with reports showing modest globally thicker cortex in ADHD^28^. The increased thickness we observed in posterior parietal and occipital regions specifically corroborates evidence ^29^, who reported a similar pattern in posterior locations in individuals with ADHD.

However, the topography of cortical alterations in ADHD is not uniform. Other studies also report thickening but in different regions, such as the somatosensory and anterior cingulate cortices, with effects that may vary by age^30^. Conversely, some research describes cortical *thinning*, particularly in motor-related brain areas ^25^. This suggests ADHD may be characterized by a mosaic of both cortical thickening and thinning across different functional systems or the contradiction could be due to methodological differences.

Taken together, our findings support the hypothesis that ADHD is characterized by altered cortical maturation. These alterations may reflect delayed synaptic pruning or region-specific disruptions in neurodevelopment, particularly in brain systems subserving cognitive control, visuospatial processing, and DMN activity.

###### Autism vs ADHD

In the direct comparison between autism and ADHD, all regions exhibiting significant differences showed increased cortical thickness in the ADHD group relative to the autism group. These regions are broadly distributed across several functional networks (Table S 5), including the DMN, then FPN; DAN; Somatomotor. A few regions were also appeared in language; Ventral-Multimodal; Visual 2; and CO network. Overall, ADHD shows thicker cortex in all above mentioned regions.

Table S 5. Significantly Different Cortical thickness ROIs in Autism Vs, ADHD comparison

| Network | N | min_Cohens’ d | max_Cohens’ d | ROIs |
| --- | --- | --- | --- | --- |
| DMN | 8 | 0.157792 | 0.419267 | R_8Av, R_OFC, R_47s, L_7m, L_10r, L_8Av, L_8Ad, L_10v |
| FPN | 5 | 0.22527 | 0.419756 | R_7Pm, R_i6-8, R_s6-8, L_POS2, L_7Pm |
| DAN | 4 | 0.320136 | 0.435252 | R_6a, R_AIP, L_6a, L_AIP |
| Somatomotor | 3 | 0.186859 | 0.409939 | R_6mp, L_4, L_2 |
| Language | 2 | 0.178047 | 0.310208 | R_55b, L_SFL |
| Ventral-Multimodal | 2 | 0.168291 | 0.189937 | R_PeEc, L_PeEc |
| Visual 2 | 2 | 0.311546 | 0.351794 | R_LO3, L_VIP |
| CO | 1 | 0.321428 | 0.321428 | R_6ma |

These findings suggest widespread cortical thinning in autism relative to ADHD, consistent with evidence that autism is associated with more pervasive neurodevelopmental alterations. Prior studies report reduced cortical thickness in autism in regions related to theory of mind, executive function, and sensory processing, particularly within heteromodal association areas ^24,25^. In contrast, ADHD is often associated with a delayed cortical maturation profile, characterized by relatively thicker cortex, particularly in frontal and posterior brain regions^28,29^.

Behaviorally, these morphological differences align with functional disparities observed between the disorders. For instance, autistic children show more severe impairments in visual attention and inhibitory control compared to both ADHD and typically developing peers^31^, possibly reflecting the thinner cortex observed in attention and sensorimotor regions in our results. Functional MRI findings further highlight distinct activation patterns: ADHD is characterized by right inferior parietal activation during inhibition tasks, whereas autism recruits frontal regions, including the middle frontal gyrus ^32^. These FPN differences may underlie the consistently thicker cortex we observed in ADHD across regions within these networks.

Together, our findings support a growing literature emphasizing dissociable neurodevelopmental trajectories in autism and ADHD, particularly in networks supporting attentional control, executive function, and self-referential processing. The consistently thinner cortex in autism across these regions may contribute to its more pronounced cognitive and sensory processing impairments.

##### 3.1.2 HCP thickness signatures

**The distinct, significantly different cortical thickness regions for each group** **are presented along with brief functional and structural descriptions.** Considering the detailed description of signature regions in the main manuscript, we provide only a general overview here. **A summary of all signature regions and their corresponding cortex is also provided in the accompanying tables for each group.**

###### Autism cortical thickness signature ROIs

The autism cohort exhibited a distinct cortical thickness signature encompassing a broad set of regions distributed across multiple large-scale functional networks (Table S 6). These ROIs were first characterized by their functional network affiliation and then by their anatomical localization, based on the HCP multimodal parcellation guide^11^.

Table S 6. Autism Cortical Thickness Signature

| ROI | Cortex Location | Network/System |
| --- | --- | --- |
| Bi-hemispheres | |  |
| 6a | Superior Premotor Cortex | DAN |
| 8Av | Dorsolateral Prefrontal Cortex (DLPFC) | DMN |
| AIP | Superior Parietal Cortex | DAN |
| PeEc | Inferior Medial Temporal Cortex | Ventral-Multimodal |
| Left Hemisphere | |  |
| 2 | Somatosensory Cortex (S1) | Somatomotor |
| 4 | Motor Cortex (M1, Primary Motor Cortex) | Somatomotor |
| 7m | Posterior Cingulate Cortex | DMN |
| 8Ad | Dorsolateral Prefrontal Cortex (DLPFC) | DMN |
| SFL | Dorsolateral Prefrontal Cortex (DLPFC) | Language |
| VIP | Superior Parietal Cortex | Visual (Visual 2) |
| Right Hemisphere | |  |
| 47s | Orbital and Polar Frontal Cortex | DMN |
| 55b | Premotor Cortex | Language |
| 6mp | Paracentral Lobular and Mid Cingulate Cortex (Supplementary Motor Area) | Somatomotor |
| 6ma | Paracentral Lobular and Mid Cingulate Cortex (Supplementary Motor Area) | CO |
| 7Pm | superior parietal cortex \| Posterior Superior Parietal lobule | FPN |
| i6-8 | Dorsolateral Prefrontal Cortex (DLPFC) | FPN |
| s6-8 | Dorsolateral Prefrontal Cortex (DLPFC) | FPN |
| LO3 | lateral occipital and posterior temporal cortex \| MT+ Complex and Neighboring Visual Areas | Visual (Visual 2) |

**Functional characterization**. Regions showing significant thickness differences were distributed across canonical large-scale networks. The DMN included bilateral 8Av (relational reasoning, executive function), left 8Ad (higher-order cognitive control), left 7m (working memory, theory of mind, emotion/language processing), and right 47s (language comprehension, narrative processing). FPN effects were observed in right 7Pm (complex cognitive processing), right i6-8 and s6-8 (working memory, cognitive control). The DAN included bilateral 6a (premotor planning, social cognition) and bilateral AIP (working memory). Language network alterations were noted in right 55b and left SFL (language processing, relational reasoning). Somatomotor Network differences involved right 6mp (motor planning, speech production), left area 4 (primary motor control), and left area 2 (primary somatosensory cortex). The CO network showed differences in right 6ma (task control, speech motor programming). The Ventral-Multimodal Network included bilateral PeEc (working memory, face processing), and the Visual Network (Visual II) showed alterations in right LO3 (peripheral visual processing) and left VIP (visuomotor integration).

**Anatomical characterization.** Anatomically, these ROIs localized to dorsolateral prefrontal cortex (8Av, 8Ad, SFL, i6-8, s6-8), superior premotor cortex (6a), premotor cortex (55b), paracentral lobular and mid-cingulate cortex / supplementary motor area (6mp, 6ma), orbital and polar frontal cortex (47s), primary motor cortex (area 4), superior parietal cortex (AIP, VIP), posterior superior parietal lobule (7Pm), posterior cingulate cortex (7m), primary somatosensory cortex (area 2), inferior medial temporal cortex (PeEc), and lateral occipital cortex (LO3).

**Hemispheric Distribution.** The thickness signature displayed both bilateral and lateralized effects. Bilateral alterations were observed in 6a, 8Av, AIP, and PeEc. Left-lateralized effects included area 2, area 4, 7m, 8Ad, SFL, and VIP. Right-lateralized effects were identified in 47s, 55b, 6ma, 6mp, 7Pm, i6-8, LO3, and s6-8. This distribution underscores both symmetric and asymmetric patterns of cortical alteration, consistent with known hemispheric specializations relevant to cognitive, language, and motor processes in autism.

The following sections describe these signature regions.

###### Bilateral ROIs Characterization

Area 6a (Superior Premotor Cortex): This region is functionally implicated in premotor planning and the preparation of voluntary movements ^33^. The Area 6a shows heightened activation during language, mathematical, and social cognition tasks^11^.

Area 8Av (Dorsolateral Prefrontal Cortex): As a subdivision of the DLPFC, Area 8Av is critical for higher-order cognitive control. Area 8Av, overall, is involved in higher-order cognitive control, particularly task-specific activation patterns linked to relational reasoning and executive processing, but lower activation in motor, memory, and social cognition tasks^11^. Specifically, Baker et al. refer to this area as being involved in the complex interpretation of visual information and attention^33^.

Area AIP (Anterior Intraparietal Area): Located in the superior parietal cortex, Area AIP exhibits a functional profile suggestive of a role in integrating sensory and cognitive information. It is more activated during social and emotional processing tasks, as well as during hand motor and working memory paradigms^11^. This pattern points to its involvement in processing salient environmental stimuli to guide motor action and cognitive operations.

Area PeEc (Perirhinal Ectorhinal Cortex): This area, located in the inferior medial temporal region, comprises the perirhinal (BA35) and ectorhinal (BA36) cortices. It demonstrates a complex functional profile with variable engagement across multiple cognitive domains. Specifically, Area PeEc shows significant activation during tasks involving face processing and gambling. Its involvement in working memory and theory of mind tasks appears to be context-dependent. Collectively, these findings highlight the engagement of PeEc in specialized memory and perceptual processes, particularly for relational reasoning and face recognition^11^.

###### Left Hemisphere ROIs Characterization:

Areas 2 and 4 (Somatosensory and Motor Cortices): These core regions of the somatomotor system showed distinct profiles. Area 4 (M1), located on the anterior bank of the central sulcus, is a heavily myelinated region central to voluntary motor control. Area 2, part of the primary somatosensory complex (S1), has a more varied functional profile, with reduced activation during motor cue and working memory contrasts but heightened activation during face-processing tasks. This suggests Area 2 is primarily involved in integrating tactile and proprioceptive information, with potential contributions to social-emotional processing^11^.

Area 7m (Posterior Cingulate Cortex): A key node of the DMN, Area 7m exhibits strong functional connectivity with lateral parietal areas. It is robustly activated during tasks requiring internal cognition, including working memory, theory of mind, relational reasoning, and emotional processing ^11^. Its differential activation in language tasks further supports its role in high-level, internally-directed cognitive processes ^11^.

Areas 8Ad and SFL (Dorsolateral Prefrontal Cortex): These two subdivisions of the DLPFC showed unique functional characteristics. Area 8Ad, in the posterior superior frontal gyrus, exhibit reduced activation during specific cognitive tasks (gambling, relational reasoning) and working memory, alongside deactivation during motor cue paradigms ^11^. Area SFL (Superior Frontal Language) demonstrated substantial functional lateralization. It is more highly activated in the left hemisphere during language and relational matching tasks, consistent with its role in the canonical language network. In contrast, the corresponding right hemisphere area is more engaged during theory of mind tasks ^11^, highlighting hemispheric specialization for language processing and relational reasoning.

Area VIP (Ventral Intraparietal Complex): Located on the medial bank of the intraparietal sulcus, Area VIP is critically involved in multisensory integration. It integrates visual and somatosensory information to support spatial awareness and action, and is crucial for distinguishing object motion from self-motion ^34^. While active during motor tasks, it shows greater activation during emotional processing and was also engaged during a range of cognitive paradigms including working memory, mathematical language, and theory of mind ^11^.

###### Right hemisphere: Characterization

Area 47s (Orbital and Polar Frontal Cortex): As the most anterior subregion of area 47 and part of the DMN, Area 47s is implicated in higher-order language functions. It is robustly activated during tasks requiring language comprehension and narrative processing (storytelling) ^11^.

Area 55b (Premotor Cortex): Situated immediately anterior to area 4, Area 55b is strongly associated with the language network ^11^. While functionally distinct from neighboring regions in classical area 6, recent work has identified the right hemisphere Area 55b as a potential core hub for music perception ^35^.

Areas 6mp & 6ma (Supplementary Motor Area): Located in the paracentral lobular and mid-cingulate cortex, these areas are involved in motor sequencing, movement planning, and speech motor programming. Area 6mp is cytoarchitectonically distinct, with higher myelination and a thinner cortex than its neighbors; altered thickness in this region may relate to deficits in motor planning, coordination, and executive function ^11,36^. Area 6ma is characterized by a unique functional profile, showing increased activity during motor cue processing but decreased activity during social cognition and relational reasoning tasks ^11^.

Area 7Pm (Medial Superior Parietal Cortex): This subdivision of area 7P functions as a hub for complex cognitive computations rather than direct sensory or motor processing. It is engaged during high-level tasks including working memory, mathematical and relational reasoning, spatial processing, and theory of mind, but shows deactivation during body-centered sensory tasks ^11^. In the right hemisphere, it is specifically implicated in processing visual motion and shape, motor learning, and attention^37^.

Areas i6-8 & s6-8 (Dorsolateral Prefrontal Cortex): These two regions occupy a transitional zone between classic Brodmann areas 6 and 8 ^11^. Area i6-8 is active during working memory and mathematical tasks but shows reduced involvement during tasks requiring narrative-mathematical integration. Area s6-8 is similarly engaged by working memory and motor cue tasks but shows less involvement in relational processing and narrative-mathematical integration, suggesting distinct roles in working memory and motor processing ^11^.

Area LO3 (Lateral Occipital Cortex): Located within the MT+ Complex, Area LO3 is a specialized visual processing area with a focus on the peripheral visual fields. Its functional profile is characterized by reduced activation or deactivation during a range of cognitive tasks, particularly those involving tools, social interaction, and relational reasoning, highlighting its specialized role within the broader visual processing network ^11^.

Recent evidence suggests that structural alteration in prefrontal regions might be partly linked to perivascular spaces (PVS) pathology. Enlargement of PVS in prefrontal white matter has been associated with DLPFC hypoactivity in autism, with stronger effects beneath the rostral middle frontal gyri and correlating with language impairment, hypersensitivity, and stereotypies ^38^. Moreover, autism severity was associated with PVS enlargement in the right precuneus and cingulate, raising the possibility that such vascular-related changes contribute to the cortical thinning observed in our DLPFC clusters. Alterations involved DLPFC-associated regions implicated in social communication and emotion regulation in autism ^39,40^.

Further, area 55b was identified as a signature region within cortical thickness signature. Beyond its language-related role ^11^, right 55b has been identified as a music perception hub ^35^ and shows strong connectivity with the mirror neuron system, implicating it in action observation and movement inhibition ^41^. This supports its role as a modulatory hub linking motor, social cognition, and inhibition in autism.

Visual regions also emerged prominently in autism, where LO3 and VIP, both parts of the visual 2 network, have been consistently implicated in altered perceptual processing in autism. For example, autistic adolescents show hyperactivation in the lateral occipital complex and reduced top-down modulation during object recognition ^42^, while early fMRI studies identified atypically low BOLD responses in ventral V2 in autism ^43^. These findings align with theories of local-oriented perceptual bias and suggest that cortical thinning in these regions may reflect disruptions in visual integration.

These findings reveal distributed cortical thinning across multiple functional networks in autism, revealing a distinct morphological signature.

###### ADHD cortical thickness signature ROIs

**ADHD-specific cortical thickness differences are particularly pronounced in the Right Orbitofrontal Complex (R_OFC), Left Ventromedial Prefrontal Cortex (L_10v), and Left Anterior Rostral Prefrontal Cortex (L_10r) (Table S 7). These regions, affiliated with the DMN, are implicated in spontaneous cognition, attentional control, and executive functions such as decision-making. Structurally, these signature areas are subdivisions of the Anterior Cingulate and Medial Prefrontal Cortex—specifically areas 10v and 10r—as well as the Orbital and Polar Frontal Cortex, notably the R_OFC** ^11^**.**

Table S 7. ADHD Cortical Thickness Signature

| ROI | Cortex Location | Network/System |
| --- | --- | --- |
| Left Hemisphere | |  |
| 10v | Anterior Cingulate and Medial Prefrontal Cortex [within the depths of the inferior anterior most region of superior frontal gyrus] | **DMN** |
| 10r | Anterior Cingulate and Medial Prefrontal Cortex | **DMN** |
| Right Hemisphere | |  |
| OFC | Orbital and Polar Frontal Cortex | **DMN** |

**The L_10v is characterized as a thick, lightly myelinated region within the medial prefrontal cortex, situated anterior to the motor and premotor areas. It is typically deactivated during cognitively demanding tasks (e.g., math vs. story processing), yet activated during face-shape contrast tasks. In contrast, the L_10r is thinner and more heavily myelinated than L_10v, and shows increased activation during language-math contrasts, while being deactivated in working memory and motor control contexts** ^11^**.**

**Overall, individuals with ADHD exhibit greater cortical thickness in these regions relative to both autistic and typically developing controls. The ADHD cortical thickness signature, defined by this regional profile, suggests atypical developmental trajectories in circuits critical for core ADHD features, including impulsivity, altered reward sensitivity, sensory integration and regulation, and possibly the processing of emotional states in relation to bodily sensations.**

###### All Group Signature Cortical Thickness ROIs

Across all groups, a shared cortical thickness signature was identified in two regions within the left hemisphere: the Posterior Superior Parietal Lobule (area 7Pm) and the Posterior Cingulate Cortex (area POS2). These regions demonstrated significant cortical thickness differences in all pairwise group comparisons, indicating robust transdiagnostic effects.

Table S 8. All Groups Cortical Thickness Signature

| Area | Thickness Signature Regions for Groups | system |
| --- | --- | --- |
| Left Hemisphere | |  |
| 7Pm | Posterior Superior Parietal cortex | FPN |
| POS2 | Posterior Cingulate Cortex | FPN |

Both 7Pm and POS2 are part of the FPN and are associated with high-level cognitive functions. Specifically, L_POS2 is considered an integrative hub involved in a broad array of cognitive tasks, while L_7Pm is implicated in complex cognitive processing.

Although the right hemisphere 7Pm was previously identified as unique to autism, the left hemisphere counterpart (L_7Pm) exhibits a distinct functional profile. According to Baker et al. (2018)^37^, L_7Pm is involved in visual motion and shape processing, spatial representation, attention, and working memory. In contrast, R_7Pm is involved in vision motion, space, vision shape, working memory, motor learning, execution, and attention. Area 7Pm is also involved in episodic memory retrieval and saccade-related activity ^33^.

Area POS2, located within the posterior cingulate cortex, is a structurally and functionally distinct region. It diverges from adjacent cortical areas in terms of myelination, cortical thickness, intrinsic functional connectivity, and task-related activation patterns. POS2 is particularly specialized in integrating information across multiple cognitive domains, particularly those separate from excluding those associated with face processing and narrative-mathematical reasoning ^11^.

#### 3.2 HCP Cortical Curvature results in original and harmonized data

**In the following section, ANCOVA results for cortical curvature ROIs are presented, beginning with pairwise group comparisons, followed by a summary of signature regions, regions that significantly differ in one group and distinguish it from the other two.**

##### 3.2.1 HCP Curvature: Pairwise group comparisons

Cortical curvature findings are present as follows for each pairwise group comparison (Figure S 14) and then distinct brain curvature signatures for each group.

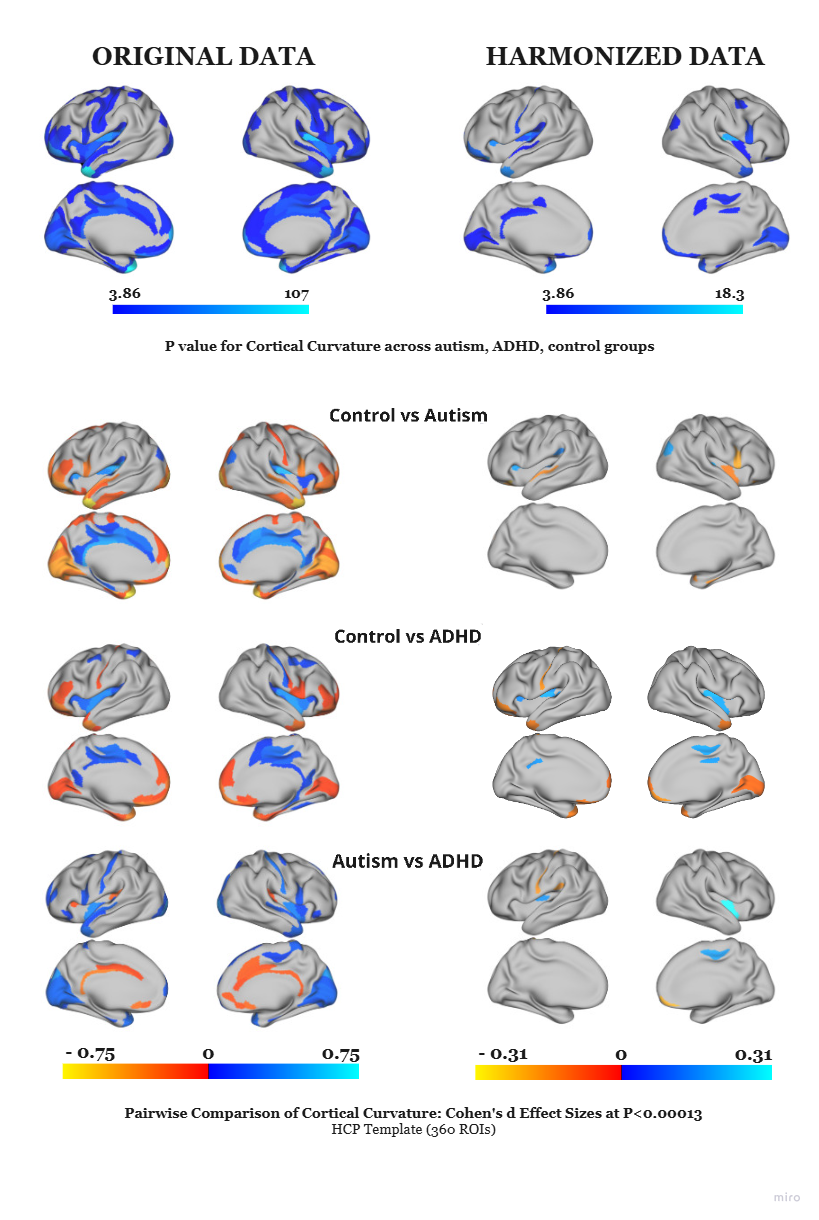

Figure S 14. HCP Cortical Curvature In Main Group and Pairwise Group Comparison

Results from ANCOVA are displayed, with the first row showing the main group effect as log_10_(p) values (p<0.05), corrected for comparisons). Subsequent rows present pairwise post-hoc comparisons for the significant regions, reported as Cohen’s d effect sizes. The first column of brain maps displays results from the original (non-harmonized) data, and the second column displays results after harmonization. Harmonization notably reduced false positives and inflated effect sizes, indicating its importance for robust cross-site analysis.

###### Control vs Autism

Significant differences in cortical curvature were observed between autistic and typically developing individuals (Table S 9), particularly within the **CO network**. Regions including R_6r and R_PoI2 were deeper in controls, whereas R_FOP3, L_PFcm, L_FOP4, and L_FOP3 were deeper in the autism group. Additional differences emerged in the **Auditory** network (R_52, R_Mbelt, L_Pbelt; deeper in autism), **DMN** regions (R_47m, L_47m; deeper in control; R_EC; more prominent in autism), and **Somatomotor** sulci (R_FOP2, L_FOP2; deeper in autism). Visual network differences (Visual 2 network) included R_V3B and L_V3B (deeper in control), and further differences appeared in **DAN** (R_PGp; more prominent in control), **FPN** (L_13l; deeper in control), **Language** (L_A5; more prominent in autism), and **Orbito-Affective** regions (R_Pir; deeper in control).

Table S 9. Significantly Different Cortical Curvature ROIs in Control Vs, Autism comparison

| Network | N | min_Cohens’ d | max_Cohens’ d | ROIs |
| --- | --- | --- | --- | --- |
| CO | 6 | 0.14198 | 0.201731 | R_6r, R_PoI2, R_FOP3, L_PFcm, L_FOP4, L_FOP3 |
| Auditory | 3 | 0.160868 | 0.253905 | R_52, R_MBelt, L_PBelt |
| DMN | 3 | 0.163453 | 0.249994 | R_47m, R_EC, L_47m |
| Somatomotor | 2 | 0.160143 | 0.167774 | R_FOP2, L_FOP2 |
| Visual 2 | 2 | 0.140243 | 0.184402 | R_V3B, L_V3B |
| DAN | 1 | 0.193893 | 0.193893 | R_PGp |
| FPN | 1 | 0.169771 | 0.169771 | L_13l |
| Language | 1 | 0.158128 | 0.158128 | L_A5 |
| Orbito-Affective | 1 | 0.15595 | 0.15595 | R_Pir |

These sulcal curvature findings align with prior evidence indicating localized shape alterations in autism. For instance, deeper right intraparietal sulcus and reduced sulcal length in the left central and medial frontal sulci have been documented in autism relative to controls ^44^. Further, atypical sulcal morphology has been reported in the anterior insula/frontal operculum and temporoparietal junction in autistic children, with inferior frontal gyrus shape contributing to anterior insular depth ^10^.

In terms of gyrification, autistic individuals demonstrate increased gyrification in regions supporting visual memory and sensorimotor function, with stronger effects in the left hemisphere, including the right inferior temporal and middle occipital gyri ^19,20^. Other studies report elevated gyrification in left parietal, temporal, and right frontal regions in autism^45^. with age-associated reductions in gyrification more pronounced in autism ^45^. A systematic review highlights early frontotemporal hypo-gyrification in autism, potentially underlying core social-communicative and sensory processing challenges ^46^.

Regarding sulcal trajectory, our finding of deeper PFcm within the Sylvian fissure (SF) in autism is consistent with prior work demonstrating anterior and superior displacement of the SF, superior frontal sulcus, and inferior frontal sulcus in autistic individuals ^47^. Additional work indicates that typical children show matched asymmetry patterns between SF and parietal cortices, whereas autistic children often show mismatched asymmetries, suggesting disrupted large-scale structural coordination ^48^.

Notably, researchers reported increased gyrification in the right superior temporal gyrus associated with greater autism-related behaviors, overlapping with regions where sulcal depth was similarly related to symptom severity. Conversely, decreased sulcal depth in the middle temporal gyrus was also linked to greater autism traits, highlighting a functional relevance of cortical morphology patterns^20^.

While some studies report no significant gyrification differences between autism, ADHD, and typically developing individuals ^49–51^, others indicate subtle, region-specific differences, particularly when accounting for behavioral and cognitive profiles. For example, typical controls show reduced gyrification in bilateral central and parietal opercula ^20^, contrasting with the higher gyrification observed in autism.

###### Control vs ADHD

Significant group differences in cortical curvature between individuals with ADHD and controls were most pronounced within the **DMN**. ADHD individuals exhibited greater gyrification in regions such as R_10d , R_10v , R_TGd , L_10d, L_10pp , L_TGd whereas R_23d was more prominent in controls. Sulcal differences also emerged, with L_d23ab deeper in ADHD and L_47m deeper in controls, further supporting distinct DMN morphological patterns across groups. Beyond the DMN, widespread sulcal depth differences were observed in additional systems. In the **Auditory Network**, sulci including R_A1, R_52, L_A1, L_52, R_Mbelt and L_Lbelt were deeper in ADHD compared to control. The **Somatomotor Network** showed deeper sulci in ADHD in regions such as R_24dd, R_Ig, L_OP2-3, and L_Ig, deeper sulcus in control is L_3a. Likewise, in the **CO network**, sulci including R_PoI2, R_FOP3, L_FOP4, and L_FOP3 were significantly deeper in the ADHD group (Table S 10).

Table S 10.Significantly Different Cortical Curvature ROIs in Control Vs, ADHD comparison

| Network | n | min_Cohens’ d | max_Cohens’d | ROIs |
| --- | --- | --- | --- | --- |
| DMN | 9 | 0.118332 | 0.178282 | R_23d, R_10d, R_10v, R_TGd, L_d23ab, L_47m, L_10d, L_10pp, L_TGd |
| Auditory | 6 | 0.11819 | 0.19786 | R_A1, R_52, L_A1, L_52, L_LBelt, R_MBelt |
| Somatomotor | 5 | 0.140182 | 0.187264 | R_24dd, R_Ig, L_3a, L_OP2-3, L_Ig |
| CO | 4 | 0.118559 | 0.167661 | R_PoI2, R_FOP3, L_FOP4, L_FOP3 |
| FPN | 2 | 0.155911 | 0.167236 | L_a47r, L_OFC |
| Visual 1 | 1 | 0.124787 | 0.124787 | R_V1 |

Although these regional effects point to morphological alterations in ADHD, particularly within networks implicated in self-referential processing, sensory integration, and motor control, findings across the literature remain mixed. Some studies report no significant differences in sulcal depth or overall curvature between individuals with ADHD and typically developing peers ^49,50^. Similarly, while the current results suggest subtle variations in gyrification, others have found no consistent differences in gyrification between ADHD, autism, and control groups—even when accounting for cognitive ability and symptom profiles such as inattention, social communication, or repetitive behaviors ^49–51^.

However, it is noteworthy that typically developing individuals exhibit lower gyrification in bilateral central and parietal opercular areas^20^, a pattern that diverges from the elevated gyrification observed in ADHD and autism in some studies. This suggests that while global measures may not always detect group differences, localized cortical folding alterations may still reflect distinct neurodevelopmental trajectories across diagnostic groups.

###### Autism vs ADHD

Group differences in cortical curvature were primarily localized to the **DMN, Somatomotor**, and **CO network**s (Table S 11). ADHD showed greater sulcal depth in R_24dd and L_Ig, while L_3a was deeper in autism. In the **CO network**, R_PoI2 was deeper in ADHD, whereas L_PFcm was deeper in autism. Additionally, ADHD exhibited greater gyrification in the DMN region R_10v. These findings suggest distinct morphometric alterations in regions supporting motor control, salience processing, and self-referential cognition, reflecting divergent neurodevelopmental trajectories in autism and ADHD.

Table S 11. Significantly Different Cortical Curvature ROIs in Autism Vs, ADHD comparison

| Network | n | min_Cohens’ d | max_Cohens’ d | ROIs |
| --- | --- | --- | --- | --- |
| Somatomotor | 3 | 0.169277 | 0.191952 | R_24dd, L_3a, L_Ig |
| CO | 2 | 0.173093 | 0.299514 | R_PoI2, L_PFcm |
| DMN | 1 | 0.213689 | 0.213689 | R_10v |

##### 3.2.2 Curvature signature

The distinct cortical curvature regions that showed significant differences between diagnostic groups are presented in Figure S 18, accompanied by brief functional and structural description. As detailed descriptions of these signature regions are provided in the main manuscript, this section offers a general overview. Supplementary tables include a comprehensive list of all curvature-based signature ROIs and their corresponding cortical areas for each group. Mean curvature values are reported to indicate whether a region corresponds to a **sulcus** (negative values) or a **gyrus** (positive values).

Structurally, the curvature signature regions were distributed across functionally distinct cortical territories. In autism, the signature region was confined to the Posterior Opercular Cortex (PFcm). In ADHD, signature regions included Somatosensory and Motor Cortex (3a), Insular and Frontal Opercular Cortex (Ig), Anterior Cingulate and Medial Prefrontal Cortex (10v), and Paracentral Lobular and Mid Cingulate Cortex (24dd). In controls, curvature signature regions were found in the Insular and Frontal Opercular Cortex (FOP3, FOP4, Area 52), Orbital and Polar Frontal Cortex (47m), and Early Auditory Cortex (MBelt). The only region common to all groups was PoI2, located within the Insular and Frontal Opercular Cortex. In the following section, we provide further details.

###### Autism cortical curvature signature ROIs

For the autism group, a unique curvature (Table S 12) alteration was observed in the CO, specifically in the left PFcm sulcus (Posterior Frontal Cortex, caudal medial). This region, located within the folds of the Sylvian fissure in the posterior opercular cortex, exhibited a distinctly deeper sulcus in autism (mean curvature = -0.357) relative to both ADHD and control groups. PFcm is implicated in sensorimotor integration and language processing ^11^, and its distinct microstructural and functional properties suggest that the observed curvature difference may reflect atypical cortical organization in autism, with potential implications for motor and cognitive function.

Table S 12. Autism Curvature Signatures

| ROI | Cortex Location | Values | Specification | Network/System |
| --- | --- | --- | --- | --- |
| Left Hemisphere | |  |  |  |
| PFcm | Posterior Opercular Cortex (within the folds of the Sylvian fissure) | -0.357 | Deeper sulcus in autism | CO |

Our analysis identifies a curvature signature in autism within the CO network, specifically in the left PFcm sulcus, a language processing & sensory motor-related region situated within the posterior opercular cortex along the Sylvian fissure ^11^, which was significantly deeper in autistic individuals compared to both ADHD and typically developing groups.

This focal alteration aligns with prior evidence of atypical sulcal morphology in autism, including depth differences along the Sylvian fissure and atypical sulcal trajectories ^47,48,52^. These findings point to early deviations in cortical folding patterns, aligning with evidence of frontotemporal hypo-gyrification in early childhood autism ^46^.

###### ADHD cortical curvature signature ROIs

In the ADHD group, four cortical regions exhibited unique curvature alterations relative to both autism and control groups: left Area 3a, left Area Ig, and right Area 24dd—all affiliated with the Somatomotor network—and right Area 10v, affiliated with the DMN. Area 3a sulcus (mean curvature = –0.6), located in the fundus of the central sulcus within the primary somatosensory cortex (S1), plays a key role in processing proprioceptive and deep sensory input. Its altered curvature suggests atypical folding within early sensorimotor integration areas. Area Ig (mean = –1.015), a granular subdivision of the insular and frontal opercular cortex, is structurally defined by high myelination and a thinner cortex, and functionally exhibits a somatotopic body map. Area 24dd (mean = –0.313), located in the dorsal anterior cingulate within the paracentral lobular and mid-cingulate cortex, is a cingulate motor area involved in contralateral upper limb motor control. This region is highly myelinated and functionally distinct from adjacent areas supporting lower limb or visceral motor processes.

Table S 13. ADHD Curvature Signature

| ROI | Cortex Location | Values | Specification | Network |
| --- | --- | --- | --- | --- |
| Left Hemisphere | |  |  |  |
| 3a | Somatosensory and Motor Cortex (Primary Somatosensory cortex (S1) | -0.6 | shallower sulcus in ADHD | Somatomotor |
| Ig | Insular and Frontal Opercular Cortex | -1.01 | Deeper sulcus in ADHD | Somatomotor |
| Right Hemisphere | |  |  |  |
| 10v | Anterior Cingulate and Medial Prefrontal Cortex [within the depths of the inferior anterior most region of superior frontal gyrus] | 0.57 | More prominent gyrus in ADHD | DMN |
| 24dd | Paracentral Lobular and Mid Cingulate Cortex | -0.31 | Shallower sulcus in ADHD | Somatomotor |

In contrast, Area 10v (mean = 0.576), situated in the ventromedial prefrontal cortex and part of the DMN, is structurally defined by low myelin content and variable cortical thickness. Functionally, it shows heterogeneous activation—often deactivated during externally oriented cognitive tasks and selectively activated during affective or introspective processes ^11^. Though data quality near the orbitofrontal border imposes some limitations^11^, the distinct curvature profile of Area 10v in ADHD may reflect atypical development of affective and self-referential processing systems. These alterations suggest atypical folding signatures across motor and DMNs, consistent with emerging evidence linking ADHD to atypical maturation in these same networks ^53^.

###### Control cortical curvature signature ROIs

In the control group, distinct sulcal curvature patterns were observed in regions affiliated with the DMN, Auditory, and CO networks. Within the auditory network, the right Area 52 (mean in control= –1.132) and right MBelt (mean = –0.324) exhibited shallower sulci in controls compared to both autism and ADHD, with MBelt being notably deeper in the autism group. In the CO network, bilateral FOP3 (right: –0.888; left: –0.878) and left FOP4 (–0.425) also showed shallower curvature in controls. In the DMN, left Area 47m (mean = –0.191), located within the orbital and polar frontal cortex, showed a deeper sulcus in controls and a shallower sulcus in autism.

Table S 14. Control Curvature Signature

| ROI | Cortex Location | Values | Specification | Network |
| --- | --- | --- | --- | --- |
| Bi-hemispheres | |  |  |  |
| FOP3 | Insular and Frontal Opercular Cortex | R_FOP3 [-0.888] & L_FOP3 [-0.878] | shallower sulci in control | CO |
| Left Hemisphere | |  |  |  |
| FOP4 | Insular and Frontal Opercular Cortex | 0.425 | shallower sulcus in control | CO |
| 47m | Orbital and Polar Frontal Cortex | -0.191 | deeper sulcus in control | DMN |
| Right Hemisphere | |  |  |  |
| MBelt | Early Auditory Cortex | -0.324 | Shallower sulcus in control | Auditory |
| 52 | Insular and Frontal Opercular Cortex | -1.132 | Shallower sulcus in control | Auditory |

Functionally, these curvature signature regions reflect distinct neuroanatomical profiles. FOP3 is a lightly myelinated and thinner region that is generally deactivated during cognitive and motor tasks, except for specific language-related contrasts (e.g., STORY–MATH), whereas FOP4 shows more consistent motor activation and higher myelination^11^. Area 47m, located within ventral prefrontal cortex, demonstrates strong face-related activation and task-driven deactivation during language processing ^11^, consistent with its role in social-affective processing, which may help explain altered face-related responses reported in autism. Furthermore, its structural connectivity with Broca’s area ^54^ underscores its contribution to affective language and communicative function.

MBelt is a highly myelinated region responsive to both auditory and emotional stimuli ^11^. In contrast, Area 52—situated between MBelt and the anterior insula—is less myelinated and functionally characterized by deactivation during language tasks and activation in bodily-related tasks ^11^. These regions, collectively, delineate the structural-functional curvature profile specific to the control group.

###### Transdiagnostic Curvature Signature Regions Across All Groups

cross all three diagnostic groups, only one cortical region—Right Posterior Insular Area 2 (PoI2)—showed statistically significant sulcal curvature differences in a transdiagnostic manner. PoI2 is located within the insular and frontal opercular cortex and is functionally affiliated with the CO network. Structurally, this region is characterized by relatively high cortical thickness and low myelination. Functionally, PoI2 is consistently activated during motor tasks, particularly those involving tongue movement (e.g., T-AVG contrast), and demonstrates variable engagement during cognitive tasks including working memory, relational reasoning, gambling, and language processing.

Table S 15. All Groups Curvature Signature

| Area | Groups Curvature Signature | Values | specification | Network |
| --- | --- | --- | --- | --- |
| Right Hemisphere | | | | |
| PoI2 | Insular and Frontal Opercular Cortex | **-0.309** | deeper sulcus in ADHD | CO |

Curvature values for the PoI2 sulcus were comparable yet significantly different across all pairwise group comparisons: autism = –0.281, ADHD = –0.309, and control = –0.294, with the deepest sulcus observed in the ADHD group. The consistent alterations across groups suggest a potential transdiagnostic structural feature relevant to sensorimotor integration or articulatory function, domains in which all three groups may show subtle differences.

However, interpretation of curvature metrics in PoI2 requires caution. As noted by Glasser et al. (2016)^11^, surface reconstruction in this region—especially in the right hemisphere—is susceptible to segmentation artifacts due to spatial overlap between cortical white matter and the adjacent putamen. These artifacts can introduce inaccuracies in the generation of cortical surfaces, directly impacting curvature estimates derived from white-to-pial geometry. Thus, while PoI2 presents a unique cross-group structural feature, findings should be considered in light of these methodological limitations.

#### 3.3 HCP Cortical Surface results in original and harmonized data

**In the following section, ANCOVA results for cortical surface ROIs are briefly presented for pairwise group comparisons.**

##### 3.3.1 HCP Surface: pairwise group comparison

Cortical surface findings are present as follows for each pairwise group comparison (Figure S 15) and then distinct brain surface signatures for each group.

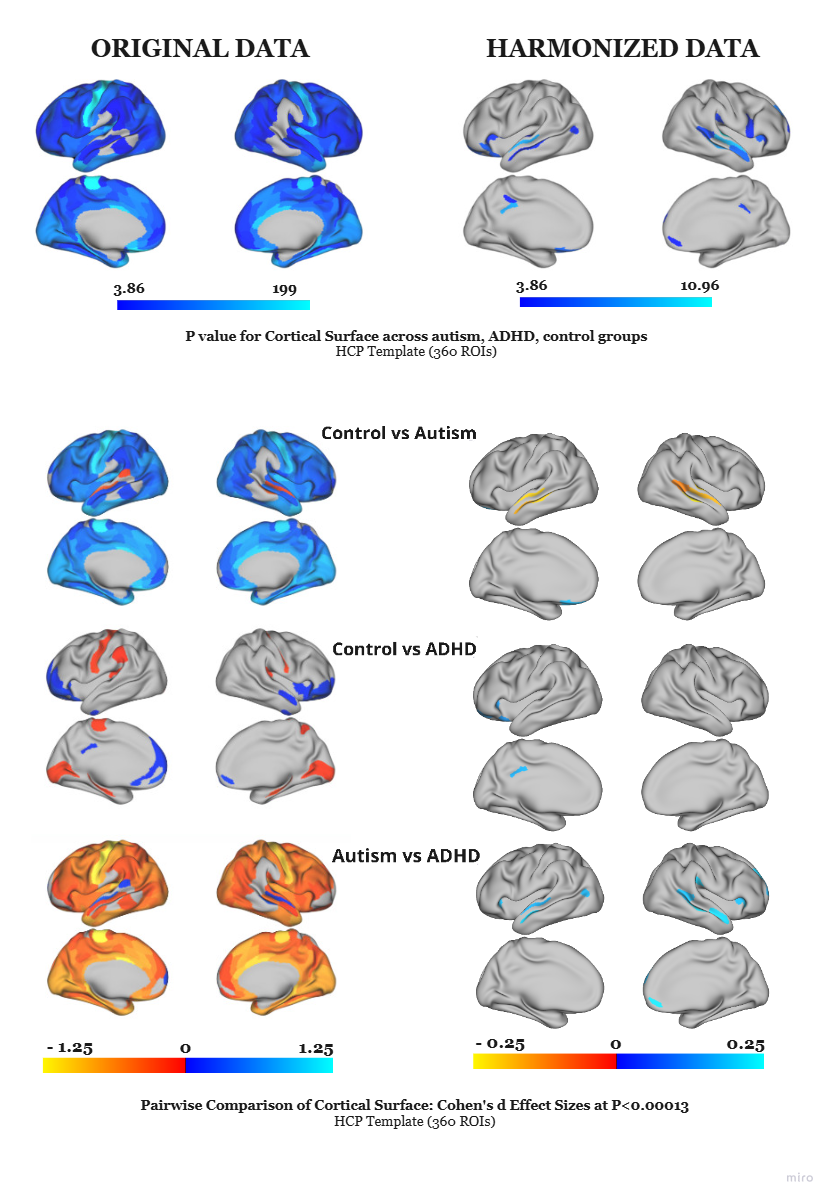

Figure S 15.HCP Cortical Surface in Main Group and Pairwise Group Comparison

Results from ANCOVA are displayed, with the first row showing the main group effect as log_10_(p) values (p<0.05), corrected for comparisons). Subsequent rows present pairwise post-hoc comparisons for the significant regions, reported as Cohen’s d effect sizes. The first column of brain maps displays results from the original (non-harmonized) data, and the second column displays results after harmonization. Harmonization notably reduced false positives and inflated effect sizes, indicating its importance for robust cross-site analysis.

###### Control vs Autism

Compared with controls, the autism group show greater surface area in DMN regions, language network regions, and auditory network (Table S 16). The only reduction is observed in the FPN. All regions except L_OFC exhibited enlarged surface area in autism. STSvp, STSva, and A5 are located within the auditory association cortex; PBelt is part of the early auditory cortex. The orbitofrontal complex (OFC) is located within the Orbital and Polar Frontal Cortex.

Table S 16.Significantly Different Cortical Surface ROIs in Control Vs, Autism comparison

| network | N | min_Cohens’ d | max_Cohens’ d | ROIs |
| --- | --- | --- | --- | --- |
| DMN | 3 | 0.165979 | 0.233688 | R_STSvp, L_STSvp, L_STSva |
| Language | 3 | 0.135709 | 0.19773 | R_A5, R_TPOJ1, L_A5 |
| Auditory | 1 | 0.200534 | 0.200534 | R_Pbelt |
| FPN | 1 | 0.164545 | 0.164545 | L_OFC |

###### Control vs ADHD

Compared with controls, individuals with ADHD show localized reductions in surface area across multiple cognitive control networks (Table S 17). Within the DMN, smaller surface areas were observed in the left dorsal 23a+b (L_d23ab) and L_47s, both located in the posterior cingulate cortex and orbital/polar frontal cortex ^11^, respectively. A similar reduction was detected in the FPN at the left lateral orbitofrontal cortex (area 11l), part of the orbital and polar frontal cortex. Additionally, the CO network exhibited reduced surface area in the left frontal opercular 5 (FOP5).

Dorsal 23a+b corresponds to Brodmann’s area 23 in the posterior cingulate cortex, where 23a and 23b are combined into 23ab and further subdivided into inferior (v23ab) and superior (d23ab) divisions. Area 47s is a subdivision of Posterior Cingulate Cortex and area 11l is a subdivision of Orbital and Polar Frontal Cortex^11^.

Table S 17. Significantly Different Cortical Surface ROIs in Control Vs, ADHD comparison

| network | N | min_Cohens’ d | max_Cohens’ d | ROIs |
| --- | --- | --- | --- | --- |
| CO | 1 | 0.122 | 0.122 | L_FOP5 |
| DMN | 2 | 0.151 | 0.151 | L_d23ab, L_47s |
| FPN | 1 | 0.127 | 0.127 | L_11l |

###### Autism vs ADHD

Compared with ADHD, autistic individuals exhibit consistently larger surface areas across multiple large-scale networks (Table S 18). Within the DMN, autism show greater surface area in the right rostral area 10 (10r), right area 9p, right area 9a, right ventral posterior superior temporal sulcus (STSvp), and left ventral anterior STS (STSva).

Table S 18. Significantly Different Cortical Surface ROIs in Autism Vs, ADHD comparison

| network | N | min_Cohens’ d | max_Cohens’ d | ROIs |
| --- | --- | --- | --- | --- |
| DMN | 5 | 0.160341 | 0.222212 | R_10r, R_9p, R_9a, R_STSvp, L_STSva |
| CO | 4 | 0.161245 | 0.199311 | R_PFcm, R_FOP4, R_FOP5, L_FOP5 |
| Language | 3 | 0.153474 | 0.208117 | R_STSda, R_TPOJ1, L_A5 |
| Visual 2 | 1 | 0.160566 | 0.160566 | L_MST |

In the CO network, larger surface areas were observed in the right PFcm, right frontal opercular area 4 (FOP4), and bilaterally in frontal opercular area 5 (FOP5). In the Language Network, the autism group show increased surface area in right-hemisphere language regions, including the dorsal anterior STS (STSda), temporo-parieto-occipital junction 1 (TPOJ1), and A5. One additional region of larger surface area was observed within the Visual 2 Network—the left medial superior temporal area (MST). MST is located within the MT+ complex and neighboring visual areas; area 10r lies within the anterior rostral prefrontal cortex; areas 9p and 9a are part of the dorsolateral prefrontal cortex (DLPFC); STSvp and STSva are subdivisions of the auditory association cortex; FOP4 and FOP5 are part of the insular and frontal opercular cortex; PFcm, TPOJ1, and A5 are subdivisions of the temporo-parieto-occipital junction. Autism shows larger surface areas in the all following regions compared to ADHD (Table S 18).

##### 3.3.2 Surface signature

The distinct cortical surface regions are presented in the Figure S 18 including brief functional and structural description. A comprehensive list of all surface-based signature ROIs and their corresponding cortical areas for autism group is presented in Table S 19. Further details are presented as follows.

###### Autism cortical surface signature ROIs

In autism, cortical surface area signatures are localized to right STSvp and left STSva in the auditory association cortex, left A5 in the temporo-parieto-occipital junction, and right TPOJ1 at the temporal–parietal–occipital interface (Table S 19). These regions, which are key nodes in language, social cognition, and multimodal integration networks, show enlarged surface area relative to ADHD and controls. Collectively, this distributed temporo-parietal network may reflect developmental adaptations supporting the integration of complex auditory, linguistic, and social information in autism. These signature regions are described further as follows.

Table S 19. Autism Surface Signature

| ROI | Cortex Location | Network/System |
| --- | --- | --- |
| R_STSvp | Auditory Association Cortex | DMN |
| L_STSva | Auditory Association Cortex | DMN |
| L_A5 | Temporo-Parieto-Occipital Junction | Language |
| R_TPOJ1 | Temporo-Parieto-Occipital Junction | Language |

The Right STSvp region, part of the auditory association cortex and DMN, is characterized by relatively higher myelination and strong functional coupling with *language and social cognition* networks. It consistently shows robust activation during narrative comprehension and theory of mind tasks, as well as during complex auditory–linguistic processing ^11^. Its functional profile suggests a central role in integrating social, emotional, and linguistic cues within the broader temporal-parietal network.

The Left STSva region, located anterior to STSvp, exhibits comparatively thinner cortex and distinct myelination patterns, with functional connectivity linked to higher-order language and social cognition regions. It is prominently activated during *narrative processing and social inference* tasks, while showing modulation during working memory, relational reasoning, and emotional decision-making ^11^. These can indicate STSva as a key node for integrating complex social-linguistic information with broader cognitive control systems.

The left A5 region, located within the temporo-parieto-occipital junction and affiliated with the language network, is characterized by distinct functional connectivity and relatively higher myelination. Functionally, A5 shows robust activation during language processing tasks, particularly narrative comprehension and semantic integration, as well as during theory of mind conditions. It demonstrates reduced engagement during working memory and relational reasoning tasks, indicating a task-selective profile optimized for high-level auditory–linguistic processing rather than executive or visuospatial functions ^11^. These properties position A5 as a key temporal language hub specialized for integrating complex speech and social communicative cues.

The right TPOJ1 region, located within the temporo-parieto-occipital junction, is a highly myelinated and structurally distinct association area, larger in the right hemisphere. Functionally, TPOJ1 shows strong activation during motor-related tasks, including motor cueing and complex movement planning, as well as during social–perceptual conditions such as face processing and theory of mind. In contrast, it demonstrates reduced activation during language–semantic contrasts, reflecting task selectivity that favors visuomotor integration and multimodal sensory processing over linguistic computation ^11^. Its connectivity profile links occipital, parietal, and superior temporal regions, supporting its role as a multimodal hub for integrating motion, body perception, and socially relevant visual information.

The evidence refers temporoparietal junction & superior temporal sulcus are the regions part of social cognition ^55^. STS and TPOJ serve as core perceptual–integration hubs for social signals, whose functional connectivity with frontal and limbic systems strengthens over development, enabling more sophisticated social cognition ^55^. Pelphrey & Perlman propose a connectivity-driven developmental model of social cognition, moving beyond “one brain region bring equal to one function” views. They argue that increasingly complex social abilities emerge from progressive integration of multiple brain systems—particularly through enhanced functional connectivity over development (via processes like myelination and synaptic pruning). In this model, regions such as the STS and TPOJ are part of the early “building blocks” of social perception—processing biological motion, gaze, and social cues—that feed forward into higher-order networks for theory of mind, narrative understanding, and social reasoning. Importantly, these areas do not act in isolation: their function and maturation are shaped by both bottom-up sensory input and top-down modulation from more advanced social cognitive systems ^55^.

###### ADHD cortical surface signature ROIs

Our findings highlight left FOP5 as the only identified surface signature region for ADHD (Table S 20). FOP5 is an anterior subdivision of the frontal operculum within the CO network, functionally distinguished by its connectivity profile and activation patterns. It shows strong engagement in motor-related processing and is consistently responsive during primary motor cues. While it participates in language-related tasks, its activation is more variable—often showing reduced activity during high-level relational or working memory demands ^11^. These functional characteristics suggest FOP5 plays a role in motor planning with broader cognitive control.

Table S 20. ADHD Surface Signature

| ROI | Cortex Location | Network/System |
| --- | --- | --- |
| Left hemisphere |  |  |
| FOP5 | insular and frontal opercular cortex | CO |

In ADHD, the reduced surface area of FOP5—a region within the CO Network—provides a surface-based signature of compromised attentional control. The CO, in close interplay with the Salience Network (SN), orchestrates the dynamic switching between the Central Executive Network (CEN) and DMN to flexibly meet task demands ^56^. A structurally diminished FOP5 may weaken this hub’s capacity to detect and prioritize behaviorally relevant stimuli and to sustain goal-directed task sets over time. Such alterations align with the ADHD phenotype, in which inefficient salience detection promotes distractibility and unstable task-set maintenance disrupts sustained attention and cognitive control.

#### 3.4 MIDB Cortical Network Thickness results in original and harmonized data

**In the following section, ANCOVA results for cortical thickness networks are briefly presented for pairwise group comparisons.**

##### 3.4.1 MIDB Network Thickness: pairwise group comparison

Consistent with trends reported in our HCP findings, our MIDB brain atlas comparisons (Figure S 16) reveal a similar directional pattern of cortical thickness across groups: autism is characterized by the thinnest cortices in several networks, ADHD by the thickest, and controls occupy an intermediate position. Several large-scale networks in the MIDB atlas show significant group differences. In the control–autism comparison, cortical thickness differences are most pronounced in the DMN, FPN, DAN, Sensorimotor Medial, and Parietal Medial networks. In the control–ADHD comparison, significant alterations are observed in the DMN, FPN, DAN, and Parietal Medial networks. The autism–ADHD comparison reveals widespread differences across the DMN, FPN, DAN, Sensorimotor Medial, and Parietal Medial networks.

##### 3.4.2 Thickness Network signature

Analysis of cortical thickness using the MIDB brain atlas reveal a distinct network-level signature only in the autism group. Specifically, autism show a unique signature in the Sensorimotor Medial network, characterized by the thinnest cortex in this area. No comparable network-level signatures were identified for ADHD or control groups. However, all three groups demonstrated significant shared differences across several large-scale networks, including the DMN, DAN, FPN, and Parietal Medial networks.

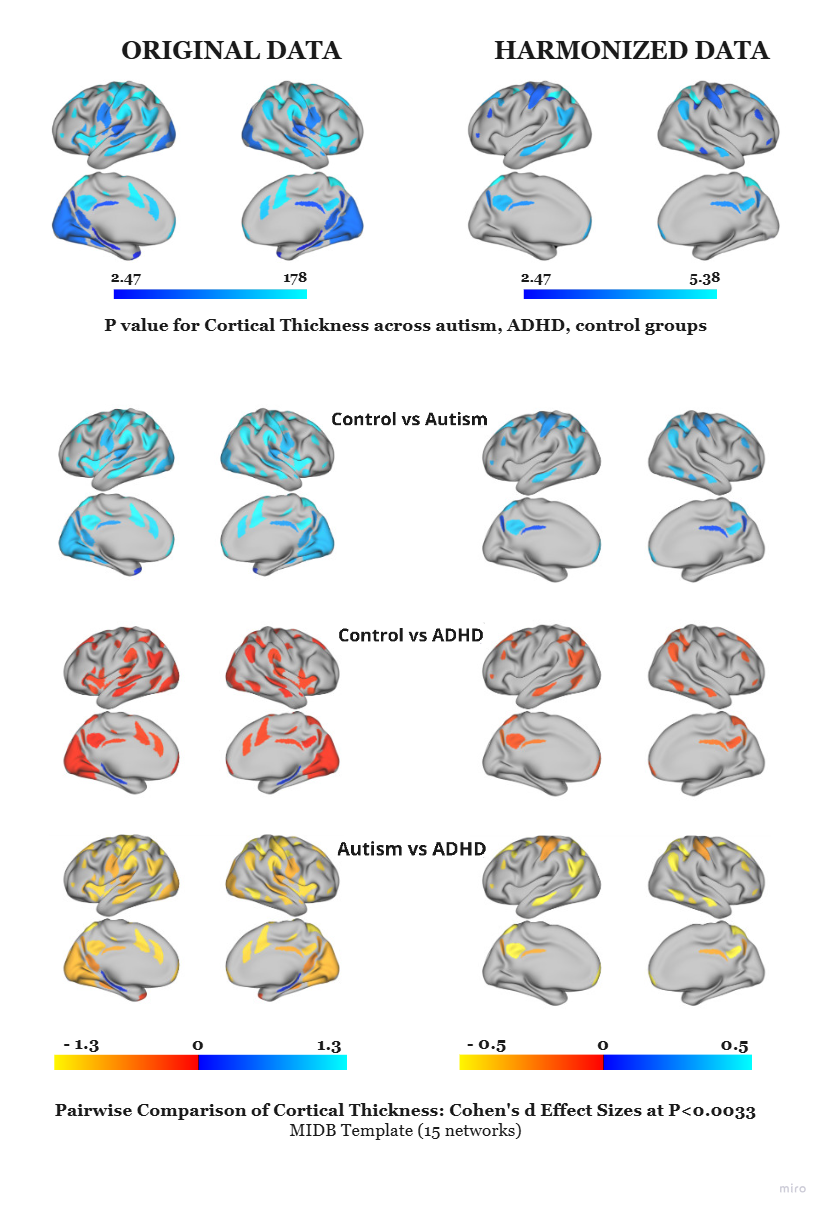

Figure S 16. MIDB Cortical Thickness Network in Main Group and Pairwise Group Comparison

Results from ANCOVA are displayed, with the first row showing the main group effect as log_10_(p) values (p<0.05), corrected for comparisons). Subsequent rows present pairwise post-hoc comparisons for the significant regions, reported as Cohen’s d effect sizes. The first column of brain maps displays results from the original (non-harmonized) data, and the second column displays results after harmonization. Harmonization notably reduced false positives and inflated effect sizes, indicating its importance for robust cross-site analysis.

#### 3.5 MIDB Cortical Thickness ROIs results in original and harmonized data

**In the following section, ANCOVA results for cortical thickness ROIs are briefly presented for pairwise group comparisons.** our MIDB brain atlas ROIs (Figure S 17) reveal a similar directional pattern of cortical thickness across groups: autism is characterized by the thinnest cortices in several regions and ADHD by the thickest regional cortex. The findings visually look similar areas to our findings in networks.

##### 3.5.1 MIDB ROI Thickness: pairwise group comparison

Cortical thickness analyses across MIDB ROIs revealed consistent group-specific patterns with MIDB network parcels and HCP ROIs, where ADHD participants exhibit greater cortical thickness, while autistic participants demonstrated the thinnest cortex, replicating the trend across parcellations.

**Control vs. Autism**: Significant ROI thinning in autism was observed within the DMN Network (DMN2, DMN3, DMN6), Dorsal Attention Network (DAN1, DAN2, DAN4, DAN5), Cingulo-Opercular Network (CO7, CO8), Sensorimotor Medial (SMm1), and Parietal Medial Network (PmN2, PmN3).

**Control vs. ADHD**: Significant ROI differences were detected in DMN (DMN4), DAN (DAN2), and Parietal Medial (PmN1, PmN2).

**Autism vs. ADHD**: Autism was characterized by thinner cortex relative to ADHD within ROIs corresponded to DMN (DMN2, DMN3, DMN4, DMN6), DAN (DAN1, DAN2, DAN4, DAN5), CO (CO7, CO8), SMm1, and PmN2.

These findings highlight distinct ROI-specific cortical thickness profiles, with ADHD showing regional thicker cortex and autism showing widespread cortical thinning relative to controls and ADHD. Cortical thickness analyses revealed a robust gradient (ADHD > Control > Autism) across MIDB ROIs, with autism showing widespread thinning in DMN, DAN, CO, parietal & Sensorimotor Medial networks, and ADHD exhibiting focal thickening relative to controls.

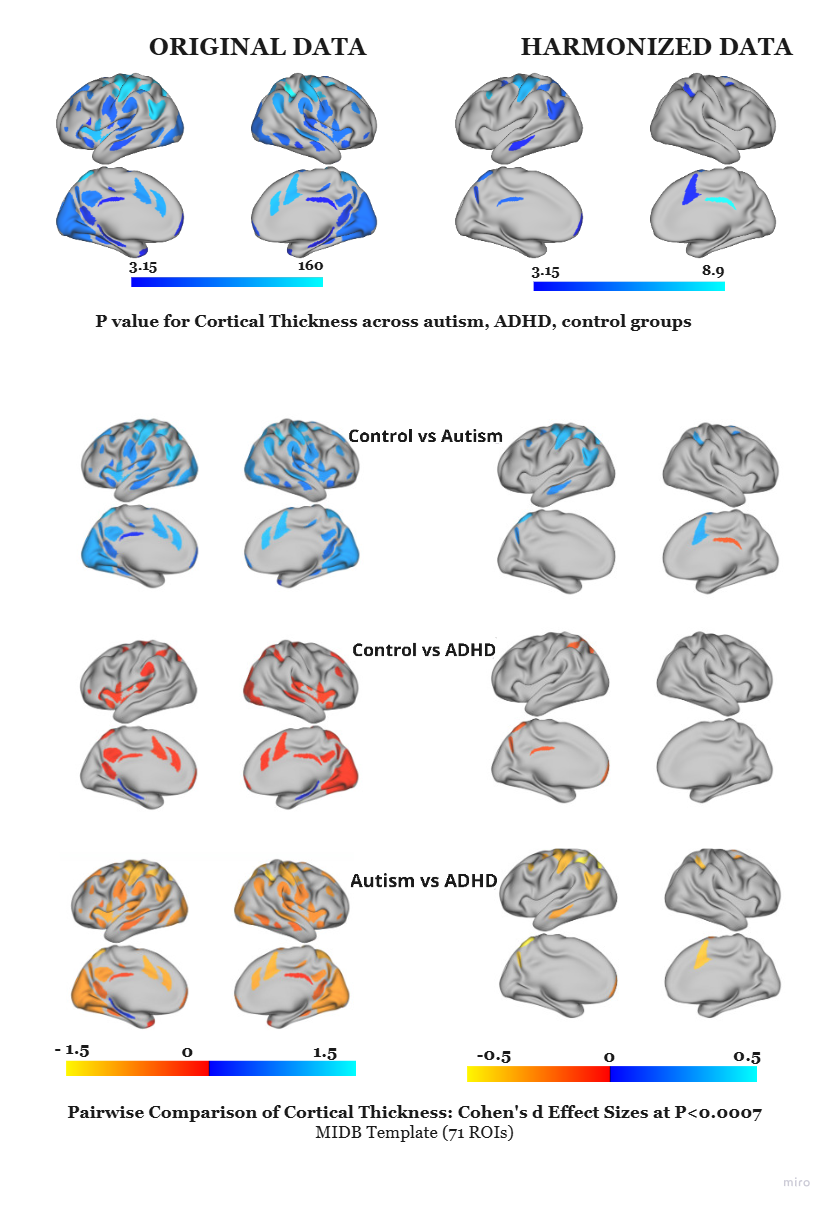

Figure S 17. MIDB Cortical Thickness ROIs in Main Group and Pairwise Group Comparison

Results from ANCOVA are displayed, with the first row showing the main group effect as log_10_(p) values (p<0.05), corrected for comparisons). Subsequent rows present pairwise post-hoc comparisons for the significant regions, reported as Cohen’s d effect sizes. The first column of brain maps displays results from the original (non-harmonized) data, and the second column displays results after harmonization. Harmonization notably reduced false positives and inflated effect sizes, indicating its importance for robust cross-site analysis.

##### 3.5.2 MIDB Thickness ROI signature

Cortical thickness analyses using the MIDB ROI template identified distinct signatures for Autism, ADHD, and shared alterations across groups (Table S 22).

**Autism-specific signature:** Autism was characterized by a multi-network pattern of cortical thickness differences (Table S 21). Within the Sensorimotor Medial network, SMm1 (LH) showed significant differences in thickness. In the DMN, differences were seen in DMN2 (LH), DMN3 (LH), and DMN6 (LH). The DAN showed alterations in DAN1 (LH), DAN4 (RH), and DAN5 (RH). The CO network exhibited differences in CO7 (RH) and CO8 (RH).

Table S 21. MIDB ROIs Signature In Autism, ADHD & All Groups

| Group | MIDB Template Network | ROI (MIDB) | Corresponding HCP ROIs (hemisphere \| areas) |
| --- | --- | --- | --- |
| Autism | Sensorimotor Medial | SMm1 | LH 1, 2, 4, 7PC, 3b, 6d |
|  | DMN | DMN2 | LH PFm, PGs, PGi |
|  |  | DMN3 | LH TE1a, TE1m, TE1p, TE2a |
|  |  | DMN6 | LH 8Av |
|  | DAN | DAN1 | LH 6a |
|  |  | DAN4 | RH 6a |
|  |  | DAN5 | RH AIP |
|  | CO network | CO7 | RH a24pr, p32pr, p24pr, R_p32pr, SCEF |
|  |  | CO8 | RH 6ma |
| ADHD | DMN | DMN4 | LH 10v, 10d |
| Shared (All groups) | DAN | DAN2 | Bilateral AIP, LIPv, 7PC, VIP, 7AL, 7Am, 7PL, IPS1, MIP |
|  | Parietal Medial | PmN2 | LH POS2 |

**ADHD-specific signature:** ADHD showed a more localized pattern, with cortical thickness differences restricted to DMN4 (LH) in the DMN. **Shared signature across groups:** Overlapping alterations in Control, Autism and ADHD were observed in DAN2 (LH) within the DAN and PmN2 (LH) within the Parietal Medial network. This pattern highlights that Autism has a broader, multi-network spanned cortical ROI thickness signature, whereas ADHD differences are more focal (appeared in DMN), with convergence in DAN and parietal networks across all groups.

### 4. Signature of Cortical Morphometry Across groups

Here, we briefly summarize all morphometric signatures identified using the HCP parcellation for each diagnostic group (autism, ADHD, and controls). The detail explanations of signature findings for cortical thickness and curvature are presented in the main manuscript in result and discussion section. Figure S 18 presents all signature regions including all morphometric features for each group, visualized on brain, highlighting significant ROIs identified from both the HCP and MIDB templates.

Table S 22. Autism Signature ROIs Across Features

| ROIs | Cortex Location | Network/System |
| --- | --- | --- |
| Thickness |  |  |
| 6a | Superior Premotor Cortex | DAN |
| 8Av | Dorsolateral Prefrontal Cortex (DLPFC) | DMN |
| AIP | Superior Parietal Cortex | DAN |
| PeEc | Inferior Medial Temporal Cortex | Ventral-Multimodal |
| 2 | Somatosensory Cortex (S1) | Somatomotor |
| 4 | Motor Cortex (M1, Primary Motor Cortex) | Somatomotor |
| 7m | Posterior Cingulate Cortex | DMN |
| 8Ad | Dorsolateral Prefrontal Cortex (DLPFC) | DMN |
| SFL | Dorsolateral Prefrontal Cortex (DLPFC) | Language |
| VIP | Superior Parietal Cortex | Visual network (Visual 2) |
| 47s | Orbital and Polar Frontal Cortex | DMN |
| 55b | Premotor Cortex | Language |
| 6mp | Paracentral Lobular and Mid Cingulate Cortex (Supplementary Motor Area) | Somatomotor |
| 6ma | Paracentral Lobular and Mid Cingulate Cortex (Supplementary Motor Area) | CO |
| 7Pm | superior parietal cortex \| Posterior Superior Parietal lobule | FPN |
| i6-8 | Dorsolateral Prefrontal Cortex (DLPFC) | FPN |
| s6-8 | Dorsolateral Prefrontal Cortex (DLPFC) | FPN |
| LO3 | lateral occipital and posterior temporal cortex\| MT+ Complex and Neighboring Visual Areas | Visual (Visual 2) |
| Curvature |  |  |
| PFcm | Posterior Opercular Cortex | CO |
| Surface |  |  |
| R_STSvp | Auditory Association Cortex | DMN |
| L_STSva | Auditory Association Cortex | DMN |
| L_A5 | Temporo-Parieto-Occipital Junction | Language |
| R_TPOJ1 | Temporo-Parieto-Occipital Junction | Language |

**Autism.** In the autism cohort, signature regions are identified in all cortical morphometric values, distributed across multiple large-scale functional networks, with distinct patterns for thickness, curvature, and surface area (Table S 22). Thickness differences are widespread, spanning the DMN, Language Network, FPN, CO network, Somatomotor Network (SMN), DAN, Ventral-Multimodal Network (VMN), and Visual Network (Visual II). Anatomically, these effects localized primarily to dorsolateral prefrontal cortex (DLPFC), superior and premotor cortices, parietal association areas, posterior cingulate, motor and somatosensory cortices, inferior medial temporal cortex, and lateral occipital cortex.

Curvature alterations in autism are more spatially circumscribed, involving the posterior opercular cortex (PFcm) within the folds of the Sylvian fissure, mapping to the CO network. Surface area differences were concentrated in temporo-parietal association cortices, including right STSvp, left STSva (auditory association cortex), left A5 (temporo-parieto-occipital junction), and right TPOJ1 (temporal–parietal–occipital interface).

**ADHD**. In ADHD, we identify signature regions across all morphometric features (Table S 23). In this regard, cortical thickness alterations are localized primarily to the anterior cingulate and medial prefrontal cortex (Areas 10v, 10r) and the orbital and polar frontal cortex (OFC), aligning with the DMN. Curvature signatures are distributed across the Somatomotor and DMNs. Notable regions included primary somatosensory cortex (Area 3a), insular/frontal opercular cortex (Area Ig), dorsal anterior cingulate within the paracentral lobular and mid-cingulate cortex (Area 24dd), and ventromedial prefrontal cortex (Area 10v). These regions collectively implicate motor integration, interoception, and affective control systems in ADHD-related folding differences. Surface area effects were observed in FOP5 (CO network), a frontal operculum subdivision engaged in motor cue processing and integrative control.

Consistent with our observed ADHD cortical signatures across the DMN, Somatomotor, and CO networks, meta-analytic evidence ^57^ shows that ADHD involves persistent dysregulation across these same systems—manifesting as hypoactivation in executive-control circuits and hyperactivation in DMN and sensorimotor networks from childhood into adulthood, independent of comorbidities or medication.

Table S 23. ADHD Signature ROIs Across Features

| ROI | Cortex Location | Network/System |
| --- | --- | --- |
| Thickness |  |  |
| 10v | Anterior Cingulate and Medial Prefrontal Cortex [within the depths of the inferior anterior most region of superior frontal gyrus] | DMN |
| 10r | Anterior Cingulate and Medial Prefrontal Cortex | DMN |
| OFC | Orbital and Polar Frontal Cortex | DMN |
| Curvature |  |  |
| 3a | Somatosensory and Motor Cortex (Primary Somatosensory cortex (S1) | Somatomotor |
| Ig | Insular and Frontal Opercular Cortex | Somatomotor |
| 10v | Anterior Cingulate and Medial Prefrontal Cortex | DMN |
| 24dd | Paracentral Lobular and Mid Cingulate Cortex | Somatomotor |
| Surface |  |  |
| FOP5 | anterior subdivision of the frontal operculum | CO |

**Control**. In the control group, distinct morphometric features are restricted to curvature differences (Table S 24), with no unique thickness or surface area signatures. Curvature alterations were observed in regions affiliated with the Auditory, CO, and DMNs. Anatomically, the differences are concentrated in the **ventral prefrontal cortex, frontal operculum,** and **auditory association cortex**.

Table S 24. Control Signature ROIs

| ROI | Cortex Location | Network/System |
| --- | --- | --- |
| Curvature |  |  |
| FOP3 | Insular and Frontal Opercular Cortex | CO |
| FOP4 | Insular and Frontal Opercular Cortex | CO |
| 47m | Orbital and Polar Frontal Cortex | DMN |
| MBelt | Early Auditory Cortex | Auditory |
| 52 | Insular and Frontal Opercular Cortex | Auditory |

**All groups**. Across all three diagnostic groups, shared morphometric differences emerged in both curvature and thickness (Table S 25). Curvature alterations were confined to the CO network within the posterior insular and frontal opercular cortex. Thickness differences were observed in the FPN, localized to the posterior superior parietal cortex and posterior cingulate cortex.

Table S 25. All groups Signature ROIs Across Features

| ROIs | Cortex Location | Network/System |
| --- | --- | --- |
| Thickness |  |  |
| 7Pm | Posterior Superior Parietal cortex | FPN |
| POS2 | Posterior Cingulate Cortex | FPN |
| Curvature |  |  |
| PoI2 | Insular and Frontal Opercular Cortex | CO |

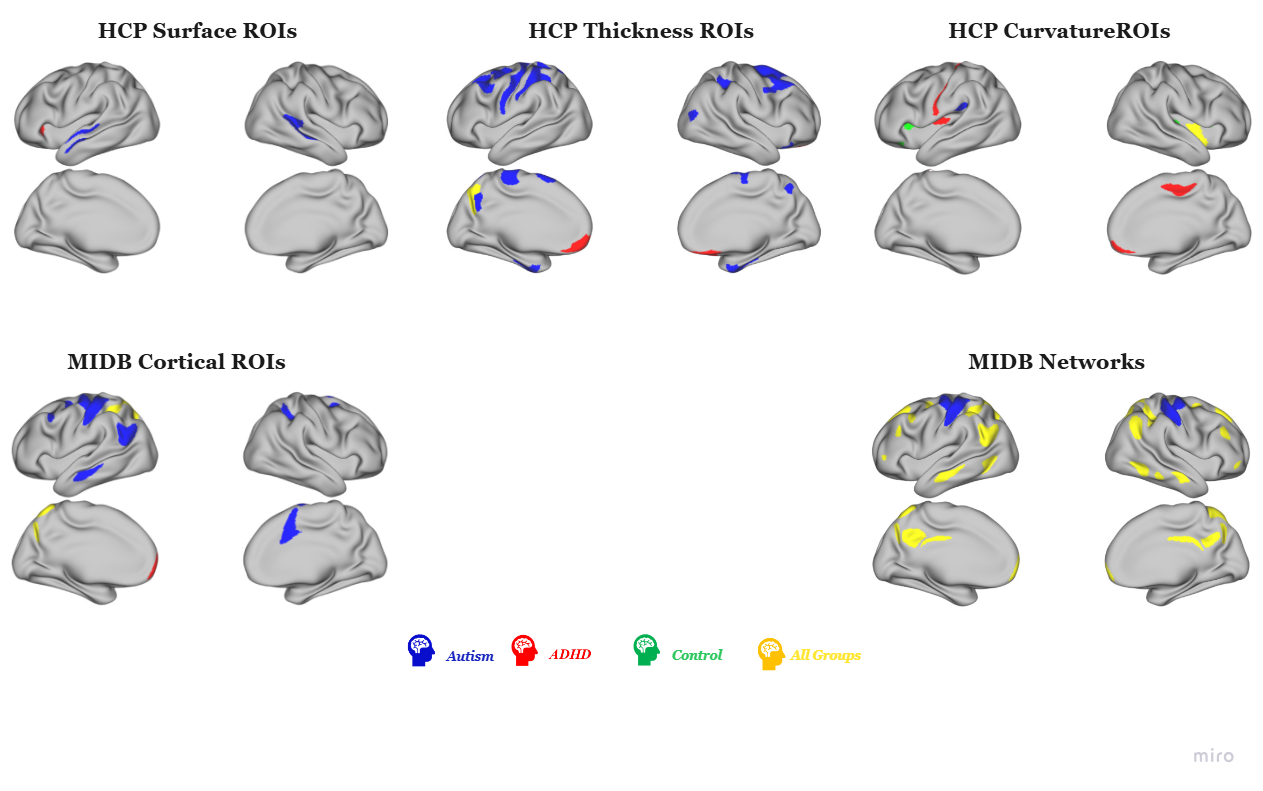

Figure S 18. Brain Signature Regions | Networks Across Morphometric Features & Templates

This figure depicts cortical signatures for HCP morphometric features (surface area, cortical thickness, and curvature) and MIDB cortical thickness, at both the parcel (ROI) and network levels in the MIDB template. Autism (blue) signatures were identified across all features and templates. ADHD (red) signatures were present across all features, although no ADHD-specific networks were detected, whereas autism showed distinct network-level signatures. Control (green) signatures were observed only in curvature. Yellow regions represent transdiagnostic areas—regions differing across all pairwise group comparisons—appearing in all features and templates except surface area.

### 5. Discussion

This supplementary discussion provides further neurobiological interpretation of the cortical thickness and curvature signatures identified in our primary analysis. We expand on the unique patterns of cortical alteration specific to the Autism and ADHD cohorts, describe the distinct structural profile of the neurotypical control group and highlight the transdiagnostic findings that point to a shared neurobiology across these conditions.

#### 5.1 Autistic Cortical Signature Regions

Our cortical thickness and curvature findings reveal a broad pattern of cortical alterations in autism spanning multiple large-scale functional networks. Thickness differences were most pronounced across dorsolateral prefrontal (DLPFC), premotor, sensorimotor, superior parietal, posterior cingulate, temporal, and visual association cortices.

Autism unique cortical thickness signature was characterized by widespread cortical thinning primarily within the higher-order transmodal system, including DMN, FPN, DAN, Language, and Ventral Multimodal, along with signature regions corresponding to sensory-related networks, including Somatomotor and Visual networks.

A central feature of autism signature is atypical morphometry in these higher-order associative network systems, providing a structural basis for the *Triple Network Model of psychopathology* ^56,58^. Specifically, alterations in the key nodes related to DMN and FPN, such as DLPFC (areas 8Av, 8Ad, i6–8, SFL) and posterior cingulate, may underlie the functional disintegration and atypical connectivity frequently reported in autism (^59^, potentially contributing to executive and social-cognitive difficulties ^39,40^.

Additionally, the CO network, critical for the hierarchical control of goal-directed behavior ^60^, showed significant alterations in both thickness and curvature. We observed sulcal depth in the left PFcm sulcus, a key CO node. This atypical folding may reflect early developmental perturbations in axonal tension or neuroinflammatory processes ^52^ and aligns with prior evidence on frontotemporal hypo-gyrification in autism ^46^.

Given the PFcm region involvement in language & sensorimotor processing ^11^, the convergence of cortical thinning and altered curvature within the CO network could represent a structural substrate for neurodevelopmental changes such as motor inflexibility in autism.

Structural changes in regions linked to social-affective processing also correspond with functional evidence. In a large fMRI study of Theory of Mind (ToM), both autism and ADHD showed atypical activation in the bilateral middle cingulate, supramarginal gyrus, and superior temporal gyrus in response to social valence conditions ^61^. Autistic children, in particular, exhibited atypical responses in the middle temporal and anterior cingulate cortices, alongside group-level differences in fusiform activation, a region central to face perception. These findings support the idea that structural alterations in autism reflect broader disruptions in brain systems supporting mentalizing and social cognition.

Furthermore, our results align with the *embodiment theory of autism*. Prior work suggests that impaired integration between sensorimotor and conceptual systems arising from subtle motor planning deficits may disrupt language acquisition and social cognition ^62,63^. Our findings of altered morphometry in premotor, sensorimotor, and frontoparietal regions support the hypothesis that reduced cortical integration in these areas may compromise information flow between sensory, motor, and cognitive domains.

Overall, the autism cortical signature reflects multilevel disruptions spanning widespread alterations across whole brain. These patterns provide structural insight into the developmental and clinical heterogeneity of the condition.

#### 5.2 ADHD Cortical Signature Regions

Our findings reveal a distinct morphometric signature in ADHD, characterized primarily by increased cortical thickness in medial prefrontal and orbitofrontal regions, alongside curvature alterations in somatosensory, insular, frontal opercular, and cingulate territories. These structural deviations were localized mainly to the DMN and Somatomotor networks.

The observed increases in cortical thickness are consistent with neurodevelopmental models positing delayed synaptic pruning, neuronal hyperexcitability, and atypical sensory processing in ADHD ^30^. In particular, thickening in the anterior insula and primary somatosensory regions may indicate disrupted maturation of circuits involved in interoception, pain perception, and proprioceptive feedback ^30^, domains frequently altered in ADHD.

Within the DMN, thickness increases in the right orbitofrontal cortex (R_OFC), left area 10r, and left area 10v underscore the potential dysmaturation in self-referential and cognitive control processes. These regions are implicated in emotional regulation and decision-making, and their altered structure aligns with functional studies reporting atypical reward processing and impaired behavioral inhibition in ADHD ^64^. Notably, these deficits have been attributed to dysregulated catecholaminergic signaling—particularly involving dopamine and norepinephrine—within prefrontal cortical circuits that underlie top-down regulation ^65^.

While increased cortical thickness was also observed in early visual regions, these areas did not contribute to the ADHD-specific morphometric signature. Interestingly, sulcal depth alterations in visual cortex were exclusive to comparisons involving autism and controls, suggesting that such alterations may be more characteristic of autism. However, emerging evidence indicates that stimulant treatment history may influence cortical maturation in visual areas, with untreated ADHD adults exhibiting reduced gray matter volumes ^53^. These findings might highlight a potential modulatory effect of medication and reinforce the importance of disentangling core anatomical alterations from treatment-related changes. These results underscore the heterogeneity of ADHD's structural neurobiology and support a model wherein both intrinsic developmental delays and extrinsic environmental factors shape cortical architecture.

A potential cellular correlate for these macrostructural atypicalities comes from experimental work in animal models, where enhancing parvalbumin-positive interneuron activity in the anterior cingulate cortex (ACC) mitigated core ADHD-like symptoms including hyperactivity, impulsivity, and attentional deficits ^66^. The altered thickness and curvature observed within medial prefrontal and ACC regions in our study may represent correlates of excitatory–inhibitory imbalances.

At the network systems level, these results are consistent with established functional disruptions in ADHD involving the cingulo-frontal-parietal (CFP) attention network, DMN, and reward/motivation circuits ^67,68^. Hypoactivation of the ACC during attentionally demanding tasks^69^ and reduced connectivity within the CFP network have been linked to cognitive control impairments, consistent with the regions altered in our results.

Alterations in the somatosensory cortex (S1) further illuminate sensorimotor integration in ADHD. Electrophysiological studies have shown atypical somatosensory-evoked potentials (SEPs), particularly changes in the N18 and N30 components, which reflect altered communication between brainstem-cerebellar structures and prefrontal cortex ^70^. While S1 activation itself may remain intact during motor tasks in ADHD, disrupted integration with higher-order regions could contribute to impairments in motor learning and sensorimotor processing. Structurally, the shallower sulcus observed in S1 among individuals with ADHD—contrasting with the deeper sulcus seen in autism—may reflect reduced anatomical differentiation or cortical specialization, possibly due to atypical developmental pressures on cortical folding. Given that sulcal depth is influenced by the tension of long-range connectivity, these findings may indicate disrupted sensorimotor-prefrontal connectivity contributing to morphological divergence.

Developmentally, increased cortical thickness in ADHD, particularly in sensory and prefrontal regions may reflect delayed cortical maturation, slower synaptic pruning or prolonged reliance on lower-order processing mechanisms (primary sensory processing), a pattern frequently observed in ADHD studies.

Finally, ADHD’s cortical architecture appears shaped by both intrinsic developmental factors and extrinsic influence such as medication exposure. Our results are consistent with meta-analytic evidence of frontoparietal dysfunction ^71^ and reduced neural flexibility and atypical organization in DMN ^72^, which has been linked to polygenic risk for ADHD ^73^.

Overall, our findings reveal a complex cortical architecture in ADHD involving prefrontal, cingulate, somatosensory, and opercular regions. These alterations appear to converge on dysfunction in attentional control, reward processing, and sensorimotor integration, core behavioral features of ADHD.

#### 5.3 Typical Brain: A Unique Signature of Cortical Curvature

The typical control group is characterized not by a unique thickness profile, but by a distinct signature of sulcal curvature. These specific curvature patterns emerge primarily within regions of the auditory, DMN, CO networks. This suggests that typical neurodevelopment may foster a more efficient or differentiated cortical folding architecture in systems supporting sensory integration and higher-order cognition. Although controls lacked a unique thickness profile relative to the clinical groups, their distinct curvature features underscore the value of curvature as a complementary morphometric feature for characterizing typical brain organization.

#### 5.4 Transdiagnostic Signatures: Evidence for Shared Neurobiology

Across all three groups, we identified convergent regions of divergence in both cortical thickness and sulcal curvature, suggesting transdiagnostic alterations. The left posterior superior parietal lobule (7Pm) and posterior cingulate cortex (POS2), key frontoparietal nodes showed significant thickness differences in all pairwise comparisons. These regions support executive function and high-level cognitive integration ^11,37^. Their altered morphometry in both neurodevelopmental groups relative to controls may reflect disruptions in normative synaptic pruning and cortical fine-tuning during adolescence.

Additionally, the posterior insular region (PoI2), part of the CO network, involved in motor processes such as tongue movement, demonstrated curvature differences across all group comparisons, with the deepest sulcus in ADHD. However, given known surface reconstruction challenges in this area (particularly in the right hemisphere) ^11^, this finding should be interpreted with caution.

#### Predictive Coding Perspective on Divergent Architectures

The *predictive coding framework* ^74^ offers a powerful lens to translate these distinct structural signatures into cognitive and behavioral phenotypes. This theory conceptualizes cognition as arising from hierarchical interactions between top-down predictions and bottom-up sensory signals, with perception and action guided by the minimization of prediction errors.

Within this model, the thinner cortex in autism within sensory and executive networks may reflect difficulties forming precise predictive models (priors), leading to an over-reliance on incoming sensory input and difficulty filtering irrelevant stimuli. This aligns with hallmark features such as sensory hypersensitivity and behavioral inflexibility. Conversely, the thicker, under-pruned cortex in ADHD may yield unstable or noisy internal models. This imprecision in top-down modulation could result in heightened sensitivity to environmental change, distractibility, and deficits in sustained attention.

#### A Roadmap Toward Translational Morphometry

Translating these morphometric findings into clinical applications requires a clear, multi-pronged research strategy. An immediate next step is to move beyond single-modality analyses. Future work should investigate interactions between morphometric features, such as cortical thickness-to-curvature ratios, and integrate these cortical findings with subcortical and white matter morphology. Future work should also clarify the relationship between the cortical signatures we identified and subcortical morphology; while subcortical volume alone may have limited diagnostic value ^75^, its interaction with cortical structure needs investigation.

As cortical morphometry reflects complex microstructural processes, future studies should integrate myelin-sensitive imaging measures, such as the T1w/T2w ratio ^76^ for elucidating the neurobiological mechanisms underpinning variations in cortical thickness and curvature, particularly within the transmodal association cortices that are characterized by protracted maturation and heightened developmental plasticity. Integrating AI and deep learning approaches is another critical step to combine cortical thickness, surface and curvature measures to capture the bigger picture on the morphometric landscape and the interrelated patterns that distinguish autism and ADHD.

### 6. Side Note Boxes

Multiple developmental theories have been proposed to explain cortical structural changes observed across childhood and adolescence. Natu et al. (2019) describe three principal mechanisms: (1) synaptic pruning, involving the elimination of redundant or inefficient synapses, dendrites, and neurons, which contributes to cortical thinning and promotes circuit efficiency; (2) myelination, wherein the increasing insulation of axons enhances transmission speed but also reduces gray–white matter contrast; and (3) cortical morphology, whereby mechanical forces associated with surface area expansion induce folding patterns that may be misinterpreted as cortical thinning. Critically, high-resolution microstructural imaging using quantitative MRI (qMRI) has shown that cortical thinning, particularly in high-level visual areas, often reflects increased intracortical myelination rather than actual tissue loss (Natu et al., 2019).

Gyrification is a hallmark of cortical development and a mechanical outcome of complex biological processes. While historically attributed to neuron proliferation, more recent models attribute folding patterns to a combination of intrinsic cytoarchitecture and extrinsic forces such as differential tangential expansion, axonal tension, dendritic arborization, gliogenesis, and neuropil growth (Essen, 1997; Rash et al., 2019; Ronan & Fletcher, 2015). Mota & Herculano-Houzel (2015) quantified this relationship, demonstrating that cortical folding scales proportionally with surface area and the square root of thickness—a principle conserved across individuals and species (Mota & Herculano-Houzel, 2015). Cortical curvature serves as a geometric proxy for fold sharpness, with higher mean curvature indicating more pronounced sulci and gyri. Curvature thus complements metrics like cortical thickness and surface area in characterizing brain morphology (King et al., 2016).

***Box 1****. Structural Mechanisms Underlying Cortical Thickness and Curvature*^77–82^

ADHD cortical thickness signatures are in three prefrontal regions: the Right Orbitofrontal Cortex (R_OFC), Left Ventromedial Prefrontal Cortex (L_10v), and Left Anterior Rostral Prefrontal Cortex (L_10r). Although these regions belong to distinct large-scale networks, they also converge functionally on affective regulation, reward processing, and higher-order cognitive control.

The R_OFC, part of the Orbital and Polar Frontal Cortex as defined by HCP, represents a region encompassing cytoarchitectonic areas 11m, 13m, 13b, and 14r (Ongür et al., 2003). Specifically, area 13m is situated along the lateral bank of the olfactory sulcus and the medial aspect of the middle orbital gyrus, while area 11m lies within the olfactory sulcus and gyrus rectus. Ongür and colleagues identified areas 11m, 13b, and 14r as forming a functional and anatomical junction between orbital regions, implicated in sensory integration and reward processing, and medial prefrontal zones involved in viscero and emotive-motor regulation. This transitional profile positions R_OFC to support the integration and relay of affective, sensory, and motivational signals. Structurally increased thickness in R_OFC may reflect developmental alterations in these integrative processes.

The L_10v and L_10r regions are within the Anterior Cingulate and Medial Prefrontal Cortex network. Area 10v (also known as area 10 or fp2 in HCP; Glasser et al., 2016) is a thick, lightly myelinated area located anterior to motor and premotor cortices. It is functionally engaged during affective tasks and shows activation to face-shape contrast and deactivated during language-based tasks such as math versus story conditions. It has been linked to affective processing systems (Bludau et al., 2014). In contrast, L_10r demonstrates functional activation during abstract reasoning and language processing (e.g., math-story contrast), but is deactivated during tasks involving working memory and motor control (e.g., Tom-Random, Relational-Match contrasts).

These ADHD-specific cortical thickness findings point to differences in prefrontal systems that (functionally) mediate impulsivity, altered reward sensitivity, emotional modulation, and higher-order executive functions. The increased thickness observed in these regions relative to both autism and control groups may reflect atypical maturation patterns in circuits central to ADHD's clinical presentation.

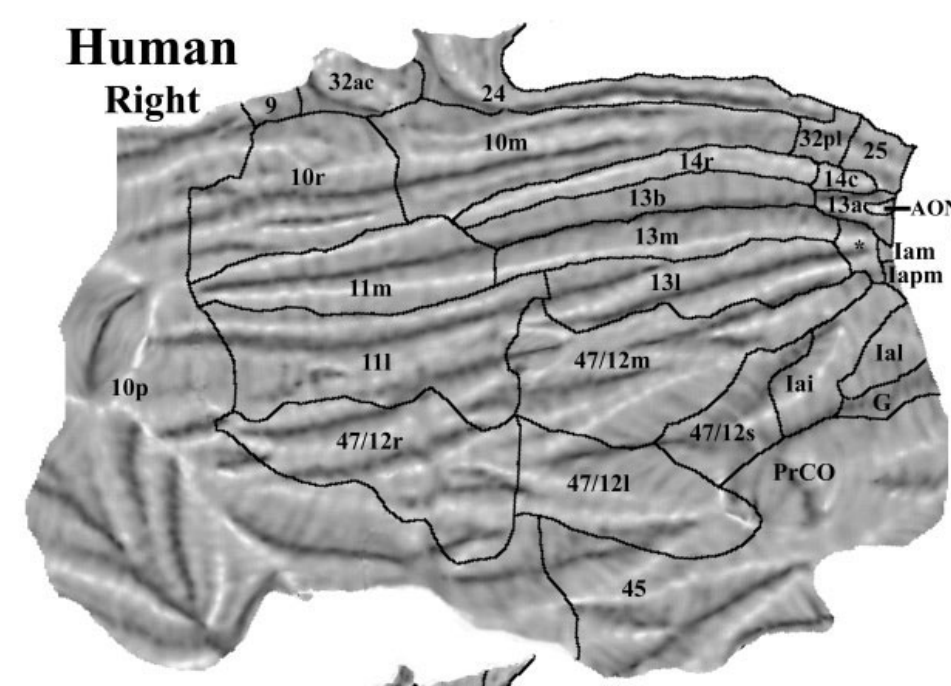

OFC subdivisions image derived from Ongür et al., (2003)

Box 2. **ADHD-Specific Cortical Thickness Alterations in Prefrontal Regions**

**Box 2.** Description of **ADHD Cortical Thickness Signatures in Prefrontal Regions** ^11,83,84^

Table S 26. Acronyms

| **Acronym** | **Full Name / Description** |
| --- | --- |
| **ABCD** | Adolescent Brain Cognitive Development Study |
| **ABIDE** | Autism Brain Imaging Data Exchange |
| **ACC** | Anterior Cingulate Cortex |
| **ADOS** | Autism Diagnostic Observation Schedule |
| **ADI-R** | Autism Diagnostic Interview – Revised |
| **ADHD** | Attention-Deficit/Hyperactivity Disorder |
| **CEN** | Central Executive Network |
| **CO** | Cingulo-Opercular Network |
| **DAN** | Dorsal Attention Network |
| **DMN** | Default Mode Network |
| **DSM-5** | Diagnostic and Statistical Manual of Mental Disorders, Fifth Edition |
| **EEG** | Electroencephalography |
| **fMRI** | Functional Magnetic Resonance Imaging |
| **FPN** | Fronto-Parietal Network |
| **HBN** | Healthy Brain Network |
| **HCP** | Human Connectome Project |
| **HCP-MMP1.0** | Human Connectome Project – Multi-Modal Parcellation (Version 1.0) |
| **KSADS** | Kiddie Schedule for Affective Disorders and Schizophrenia |
| **LH** | Left Hemisphere |
| **MIDB** | Masonic Institute for the Developing Brain |
| **MRI** | Magnetic Resonance Imaging |
| **OHSU** | Oregon Health & Science University |
| **PON** | Parieto-Occipital Network |
| **PMN** | Parietal Medial Network |
| **RH** | Right Hemisphere |
| **ROI** | Region of Interest |
| **Sal / SN** | Salience Network |
| **SCAN** | Somato-Cognitive Action Network |
| **SMd** | Sensorimotor Dorsal Network |
| **SMl** | Sensorimotor Lateral Network |
| **T1w/T2w** | T1-weighted / T2-weighted |
| **ToM** | Theory of Mind |
| **TPJ** | Temporoparietal Junction |
| **VAN** | Ventral Attention Network |
| **Vis** | Visual |

**References**

1.	Alexander LM, Escalera J, Ai L, et al. An open resource for transdiagnostic research in pediatric mental health and learning disorders. *Sci Data*. 2017;4(1):170181. doi:10.1038/sdata.2017.181

2.	ABCD Study. ABCD Study. Accessed June 17, 2024. https://abcdstudy.org/

3.	Feczko E, Conan G, Marek S, et al. Adolescent Brain Cognitive Development (ABCD) Community MRI Collection and Utilities. Published online July 11, 2021:2021.07.09.451638. doi:10.1101/2021.07.09.451638

4.	Fombonne E, Coppola L, Mastel S, O’Roak BJ. Validation of Autism Diagnosis and Clinical Data in the SPARK Cohort. *J Autism Dev Disord*. 2022;52(8):3383-3398. doi:10.1007/s10803-021-05218-y

5.	Cordova MM, Antovich DM, Ryabinin P, et al. Attention-Deficit/Hyperactivity Disorder: Restricted Phenotypes Prevalence, Comorbidity, and Polygenic Risk Sensitivity in the ABCD Baseline Cohort. *J Am Acad Child Adolesc Psychiatry*. 2022;61(10):1273-1284. doi:10.1016/j.jaac.2022.03.030

6.	Nigg JT, Karalunas SL, Mooney MA, et al. The Oregon ADHD-1000: A new longitudinal data resource enriched for clinical cases and multiple levels of analysis. *Dev Cogn Neurosci*. 2023;60:101222. doi:10.1016/j.dcn.2023.101222

7.	Di Martino A, Yan CG, Li Q, et al. The autism brain imaging data exchange: towards a large-scale evaluation of the intrinsic brain architecture in autism. *Mol Psychiatry*. 2014;19(6):659-667. doi:10.1038/mp.2013.78

8.	Di Martino A, O’Connor D, Chen B, et al. Enhancing studies of the connectome in autism using the autism brain imaging data exchange II. *Sci Data*. 2017;4(1):170010. doi:10.1038/sdata.2017.10

9.	ABIDE. Accessed September 15, 2024. https://fcon_1000.projects.nitrc.org/indi/abide/

10.	Dierker DL, Feczko E, John R. Pruett J, et al. Analysis of Cortical Shape in Children with Simplex Autism. *Cereb Cortex N Y NY*. 2015;25(4):1042. doi:10.1093/cercor/bht294

11.	Glasser MF, Coalson TS, Robinson EC, et al. A multi-modal parcellation of human cerebral cortex. *Nature*. 2016;536(7615):171-178. doi:10.1038/nature18933

12.	Ji JL, Spronk M, Kulkarni K, Repovš G, Anticevic A, Cole MW. Mapping the human brain’s cortical-subcortical functional network organization. *NeuroImage*. 2019;185:35-57. doi:10.1016/j.neuroimage.2018.10.006

13.	Hermosillo RJM, Moore LA, Feczko E, et al. A precision functional atlas of personalized network topography and probabilities. *Nat Neurosci*. 2024;27(5):1000-1013. doi:10.1038/s41593-024-01596-5

14.	Gordon EM, Chauvin RJ, Van AN, et al. A somato-cognitive action network alternates with effector regions in motor cortex. *Nature*. 2023;617(7960):351-359. doi:10.1038/s41586-023-05964-2

15.	Fortin JP. Jfortin1/neuroCombat_Rpackage. Published online November 30, 2023. Accessed April 30, 2024. https://github.com/Jfortin1/neuroCombat_Rpackage

16.	Fortin JP, Cullen N, Sheline YI, et al. Harmonization of cortical thickness measurements across scanners and sites. *NeuroImage*. 2018;167:104-120. doi:10.1016/j.neuroimage.2017.11.024

17.	Braden BB, Riecken C. Thinning faster? Age-related cortical thickness differences in adults with autism spectrum disorder. *Res Autism Spectr Disord*. 2019;64:31-38. doi:10.1016/j.rasd.2019.03.005

18.	Ecker C, Shahidiani A, Feng Y, et al. The effect of age, diagnosis, and their interaction on vertex-based measures of cortical thickness and surface area in autism spectrum disorder. *J Neural Transm*. 2014;121(9):1157-1170. doi:10.1007/s00702-014-1207-1

19.	Pereira AM, Campos BM, Coan AC, et al. Differences in Cortical Structure and Functional MRI Connectivity in High Functioning Autism. *Front Neurol*. 2018;9:539. doi:10.3389/fneur.2018.00539

20.	Zoltowski AR, Lyu I, Failla M, et al. Cortical Morphology in Autism: Findings from a Cortical Shape-Adaptive Approach to Local Gyrification Indexing. *Cereb Cortex N Y NY*. 2021;31(11):5188. doi:10.1093/cercor/bhab151

21.	Hadjikhani N, Joseph RM, Snyder J, Tager-Flusberg H. Anatomical differences in the mirror neuron system and social cognition network in autism. *Cereb Cortex N Y N 1991*. 2006;16(9):1276-1282. doi:10.1093/cercor/bhj069

22.	Wallace GL, Dankner N, Kenworthy L, Giedd JN, Martin A. Age-related temporal and parietal cortical thinning in autism spectrum disorders. *Brain*. 2010;133(12):3745-3754. doi:10.1093/brain/awq279

23.	Laidi C, Boisgontier J, de Pierrefeu A, et al. Decreased Cortical Thickness in the Anterior Cingulate Cortex in Adults with Autism. *J Autism Dev Disord*. 2019;49(4):1402-1409. doi:10.1007/s10803-018-3807-3

24.	Richter J, Henze R, Vomstein K, et al. Reduced cortical thickness and its association with social reactivity in children with autism spectrum disorder. *Psychiatry Res Neuroimaging*. 2015;234(1):15-24. doi:10.1016/j.pscychresns.2015.06.011

25.	You W, Li Q, Chen L, et al. Common and distinct cortical thickness alterations in youth with autism spectrum disorder and attention-deficit/hyperactivity disorder. *BMC Med*. 2024;22(1):92. doi:10.1186/s12916-024-03313-2

26.	Zielinski BA, Prigge MBD, Nielsen JA, et al. Longitudinal changes in cortical thickness in autism and typical development. *Brain*. 2014;137(6):1799-1812. doi:10.1093/brain/awu083

27.	Bedford SA, Park MTM, Devenyi GA, et al. Large-scale analyses of the relationship between sex, age and intelligence quotient heterogeneity and cortical morphometry in autism spectrum disorder. *Mol Psychiatry*. 2020;25(3):614-628. doi:10.1038/s41380-019-0420-6

28.	Bedford SA, Lai MC, Lombardo MV, et al. Brain-charting autism and attention deficit hyperactivity disorder reveals distinct and overlapping neurobiology. *Biol Psychiatry*. Published online August 14, 2024. doi:10.1016/j.biopsych.2024.07.024

29.	Almeida Montes LG, Prado Alcántara H, Martínez García RB, De La Torre LB, Ávila Acosta D, Duarte MG. Brain Cortical Thickness in ADHD: Age, Sex, and Clinical Correlations. *J Atten Disord*. 2013;17(8):641-654. doi:10.1177/1087054711434351

30.	Duerden EG, Tannock R, Dockstader C. Altered cortical morphology in sensorimotor processing regions in adolescents and adults with attention-deficit/hyperactivity disorder. *Brain Res*. 2012;1445:82-91. doi:10.1016/j.brainres.2012.01.034

31.	Corbett BA, Constantine LJ. Autism and attention deficit hyperactivity disorder: assessing attention and response control with the integrated visual and auditory continuous performance test. *Child Neuropsychol J Norm Abnorm Dev Child Adolesc*. 2006;12(4-5):335-348. doi:10.1080/09297040500350938

32.	Albajara Sáenz A, Septier M, Van Schuerbeek P, et al. ADHD and ASD: distinct brain patterns of inhibition-related activation? *Transl Psychiatry*. 2020;10(1):1-10. doi:10.1038/s41398-020-0707-z

33.	Baker CM, Burks JD, Briggs RG, et al. A Connectomic Atlas of the Human Cerebrum—Chapter 3: The Motor, Premotor, and Sensory Cortices. *Oper Neurosurg*. 2018;15(Suppl 1):S75. doi:10.1093/ons/opy256

34.	Field DT, Biagi N, Inman LA. The role of the ventral intraparietal area (VIP/pVIP) in the perception of object-motion and self-motion. *NeuroImage*. 2020;213:116679. doi:10.1016/j.neuroimage.2020.116679

35.	Siman-Tov T, Gordon CR, Avisdris N, et al. The rediscovered motor-related area 55b emerges as a core hub of music perception. *Commun Biol*. 2022;5(1):1-13. doi:10.1038/s42003-022-04009-0

36.	Fischl B, Rajendran N, Busa E, et al. Cortical Folding Patterns and Predicting Cytoarchitecture. *Cereb Cortex N Y NY*. 2008;18(8):1973-1980. doi:10.1093/cercor/bhm225

37.	Baker CM, Burks JD, Briggs RG, et al. A Connectomic Atlas of the Human Cerebrum—Chapter 7: The Lateral Parietal Lobe. *Oper Neurosurg*. 2018;15(Suppl 1):S295. doi:10.1093/ons/opy261

38.	Sotgiu S, Cavassa V, Puci MV, et al. Enlarged perivascular spaces under the dorso-lateral prefrontal cortex and severity of autism. *Sci Rep*. 2025;15(1):8142. doi:10.1038/s41598-025-92913-w

39.	Qiu J, Kong X, Li J, et al. Transcranial Direct Current Stimulation (tDCS) over the Left Dorsal Lateral Prefrontal Cortex in Children with Autism Spectrum Disorder (ASD). *Neural Plast*. 2021;2021(1):6627507. doi:10.1155/2021/6627507

40.	Zemestani M, Hoseinpanahi O, Salehinejad MA, Nitsche MA. The impact of prefrontal transcranial direct current stimulation (tDCS) on theory of mind, emotion regulation and emotional-behavioral functions in children with autism disorder: A randomized, sham-controlled, and parallel-group study. *Autism Res Off J Int Soc Autism Res*. 2022;15(10):1985-2003. doi:10.1002/aur.2803

41.	Rech F, Herbet G, Gaudeau Y, et al. A probabilistic map of negative motor areas of the upper limb and face: a brain stimulation study. *Brain J Neurol*. 2019;142(4):952-965. doi:10.1093/brain/awz021

42.	Sapey-Triomphe LA, Boets B, Eylen LV, et al. Ventral stream hierarchy underlying perceptual organization in adolescents with autism. *NeuroImage Clin*. 2020;25:102197. doi:10.1016/j.nicl.2020.102197

43.	Bölte S, Hubl D, Dierks T, Holtmann M, Poustka F. An fMRI-study of locally oriented perception in autism: altered early visual processing of the block design test. *J Neural Transm*. 2008;115(3):545-552. doi:10.1007/s00702-007-0850-1

44.	Auzias G, Viellard M, Takerkart S, et al. Atypical sulcal anatomy in young children with autism spectrum disorder. *NeuroImage Clin*. 2014;4:593. doi:10.1016/j.nicl.2014.03.008

45.	Kohli JS, Kinnear MK, Fong CH, Fishman I, Carper RA, Müller RA. Local Cortical Gyrification is Increased in Children With Autism Spectrum Disorders, but Decreases Rapidly in Adolescents. *Cereb Cortex N Y NY*. 2018;29(6):2412. doi:10.1093/cercor/bhy111

46.	Sasabayashi D, Takahashi T, Takayanagi Y, Suzuki M. Anomalous brain gyrification patterns in major psychiatric disorders: a systematic review and transdiagnostic integration. *Transl Psychiatry*. 2021;11(1):1-12. doi:10.1038/s41398-021-01297-8

47.	Levitt JG, Blanton RE, Smalley S, et al. Cortical sulcal maps in autism. *Cereb Cortex N Y N 1991*. 2003;13(7):728-735. doi:10.1093/cercor/13.7.728

48.	Knaus TA, Tager-Flusberg H, Foundas AL. Sylvian fissure and parietal anatomy in children with autism spectrum disorder. *Behav Neurol*. 2012;25(4):327-339. doi:10.3233/BEN-2012-110214

49.	Forde NJ, Ronan L, Zwiers MP, et al. No Association between Cortical Gyrification or Intrinsic Curvature and Attention-deficit/Hyperactivity Disorder in Adolescents and Young Adults. *Front Neurosci*. 2017;11. doi:10.3389/fnins.2017.00218

50.	Gharehgazlou A, Freitas C, Ameis SH, et al. Cortical Gyrification Morphology in Individuals with ASD and ADHD across the Lifespan: A Systematic Review and Meta-Analysis. *Cereb Cortex*. 2021;31(5):2653-2669. doi:10.1093/cercor/bhaa381

51.	Gharehgazlou A, Vandewouw M, Ziolkowski J, et al. Cortical Gyrification Morphology in ASD and ADHD: Implication for Further Similarities or Disorder-Specific Features? *Cereb Cortex*. 2022;32(11):2332-2342. doi:10.1093/cercor/bhab326

52.	Nordahl CW, Dierker D, Mostafavi I, et al. Cortical Folding Abnormalities in Autism Revealed by Surface-Based Morphometry. *J Neurosci*. 2007;27(43):11725-11735. doi:10.1523/JNEUROSCI.0777-07.2007

53.	Ghozy S, Meiza J, Morsy A, et al. How psychostimulant treatment changes the brain morphometry in adults with ADHD: sMRI Comparison study to medication-naïve adults with ADHD. *Psychiatry Res Neuroimaging*. 2025;349:111992. doi:10.1016/j.pscychresns.2025.111992

54.	Kim SJ, Tanglay O, Chong EHN, et al. Functional connectivity in ADHD children doing Go/No-Go tasks: An fMRI systematic review and meta-analysis. *Transl Neurosci*. 2023;14(1):20220299. doi:10.1515/tnsci-2022-0299

55.	Pelphrey KA, Carter EJ. Charting the typical and atypical development of the social brain. *Dev Psychopathol*. 2008;20(4):1081-1102. doi:10.1017/S0954579408000515

56.	Menon V. Large-scale brain networks and psychopathology: a unifying triple network model. *Trends Cogn Sci*. 2011;15(10):483-506. doi:10.1016/j.tics.2011.08.003

57.	Cortese S, Kelly C, Chabernaud C, et al. Toward systems neuroscience of ADHD: a meta-analysis of 55 fMRI studies. *Am J Psychiatry*. 2012;169(10):1038-1055. doi:10.1176/appi.ajp.2012.11101521

58.	Menon V. The Triple Network Model, Insight, and Large-Scale Brain Organization in Autism. *Biol Psychiatry*. 2018;84(4):236. doi:10.1016/j.biopsych.2018.06.012

59.	Yerys BE, Gordon EM, Abrams DN, et al. Default mode network segregation and social deficits in autism spectrum disorder: Evidence from non-medicated children. *NeuroImage Clin*. 2015;9:223-232. doi:10.1016/j.nicl.2015.07.018

60.	D’Andrea CB, Laumann TO, Newbold DJ, et al. Substructure of the brain’s Cingulo-Opercular network. Published online October 10, 2023:2023.10.10.561772. doi:10.1101/2023.10.10.561772

61.	Vandewouw MM, Safar K, Mossad SI, et al. Do shapes have feelings? Social attribution in children with autism spectrum disorder and attention-deficit/hyperactivity disorder. *Transl Psychiatry*. 2021;11(1):1-11. doi:10.1038/s41398-021-01625-y

62.	Eigsti IM. A Review of Embodiment in Autism Spectrum Disorders. *Front Psychol*. 2013;4:224. doi:10.3389/fpsyg.2013.00224

63.	Moseley RL, Pulvermüller F. What can autism teach us about the role of sensorimotor systems in higher cognition? New clues from studies on language, action semantics, and abstract emotional concept processing. *Cortex*. 2018;100:149-190. doi:10.1016/j.cortex.2017.11.019

64.	Cao A, Hong D, Che C, et al. The distinct role of orbitofrontal and medial prefrontal cortex in encoding impulsive choices in an animal model of attention deficit hyperactivity disorder. *Front Behav Neurosci*. 2023;16. doi:10.3389/fnbeh.2022.1039288

65.	Arnsten AFT. Fundamentals of attention-deficit/hyperactivity disorder: circuits and pathways. *J Clin Psychiatry*. 2006;67 Suppl 8:7-12.

66.	Jendryka MM, Lewin U, van der Veen B, et al. Control of sustained attention and impulsivity by Gq-protein signalling in parvalbumin interneurons of the anterior cingulate cortex. *Transl Psychiatry*. 2023;13(1):1-12. doi:10.1038/s41398-023-02541-z

67.	Bush G. Attention-Deficit/Hyperactivity Disorder and Attention Networks. *Neuropsychopharmacology*. 2010;35(1):278-300. doi:10.1038/npp.2009.120

68.	Bush G. Cingulate, Frontal and Parietal Cortical Dysfunction in Attention-Deficit/Hyperactivity Disorder. *Biol Psychiatry*. 2011;69(12):1160. doi:10.1016/j.biopsych.2011.01.022

69.	Bush G, Frazier JA, Rauch SL, et al. Anterior cingulate cortex dysfunction in attention-deficit/hyperactivity disorder revealed by fMRI and the counting stroop. *Biol Psychiatry*. 1999;45(12):1542-1552. doi:10.1016/S0006-3223(99)00083-9

70.	McCracken HS, Murphy B, Ambalavanar U, Zabihhosseinian M, Yielder PC. Sensorimotor integration and motor learning during a novel visuomotor tracing task in young adults with attention-deficit/hyperactivity disorder. *J Neurophysiol*. 2023;129(1):247-261. doi:10.1152/jn.00173.2022

71.	Wang M, Yu J, Kim HD, Cruz AB. Neural correlates of executive function and attention in children with ADHD: An ALE meta-analysis of task-based functional connectivity studies. *Psychiatry Res*. 2025;345:116338. doi:10.1016/j.psychres.2024.116338

72.	Yin W, Li T, Mucha PJ, et al. Altered neural flexibility in children with attention-deficit/hyperactivity disorder. *Mol Psychiatry*. 2022;27(11):4673-4679. doi:10.1038/s41380-022-01706-4

73.	Sato JR, Biazoli CE, Bueno APA, et al. Polygenic risk score for attention‐deficit/hyperactivity disorder and brain functional networks segregation in a community‐based sample. *Genes Brain Behav*. 2023;22(2):e12838. doi:10.1111/gbb.12838

74.	Marais AL, Roche-Labarbe N. Predictive coding and attention in developmental cognitive neuroscience and perspectives for neurodevelopmental disorders. *Dev Cogn Neurosci*. 2025;72:101519. doi:10.1016/j.dcn.2025.101519

75.	Mooney MA, Bhatt P, Hermosillo RJM, et al. Smaller total brain volume but not subcortical structure volume related to common genetic risk for ADHD. *Psychol Med*. 2021;51(8):1279-1288. doi:10.1017/S0033291719004148

76.	Norbom LB, Syed B, Kjelkenes R, et al. Probing Autism and ADHD subtypes using cortical signatures of the T1w/T2w-ratio and morphometry. *NeuroImage Clin*. 2025;45:103736. doi:10.1016/j.nicl.2025.103736

77.	Natu VS, Gomez J, Barnett M, et al. Apparent thinning of human visual cortex during childhood is associated with myelination. *Proc Natl Acad Sci U S A*. 2019;116(41):20750. doi:10.1073/pnas.1904931116

78.	Essen DCV. A tension-based theory of morphogenesis and compact wiring in the central nervous system. *Nature*. 1997;385(6614):313-318. doi:10.1038/385313a0

79.	King JB, Lopez-Larson MP, Yurgelun-Todd DA. Mean cortical curvature reflects cytoarchitecture restructuring in mild traumatic brain injury. *NeuroImage Clin*. 2016;11:81-89. doi:10.1016/j.nicl.2016.01.003

80.	Rash BG, Duque A, Morozov YM, Arellano JI, Micali N, Rakic P. Gliogenesis in the outer subventricular zone promotes enlargement and gyrification of the primate cerebrum. *Proc Natl Acad Sci U S A*. 2019;116(14):7089-7094. doi:10.1073/pnas.1822169116

81.	Mota B, Herculano-Houzel S. BRAIN STRUCTURE. Cortical folding scales universally with surface area and thickness, not number of neurons. *Science*. 2015;349(6243):74-77. doi:10.1126/science.aaa9101

82.	Ronan L, Fletcher PC. From genes to folds: a review of cortical gyrification theory. *Brain Struct Funct*. 2015;220(5):2475-2483. doi:10.1007/s00429-014-0961-z

83.	Ongür D, Ferry AT, Price JL. Architectonic subdivision of the human orbital and medial prefrontal cortex. *J Comp Neurol*. 2003;460(3):425-449. doi:10.1002/cne.10609

84.	Bludau S, Eickhoff SB, Mohlberg H, et al. Cytoarchitecture, probability maps and functions of the human frontal pole. *NeuroImage*. 2014;93 Pt 2(Pt 2):260-275. doi:10.1016/j.neuroimage.2013.05.052
